# Imputation approaches and quality standards for single-cell epigenetic age predictions

**DOI:** 10.1101/2023.12.14.571557

**Authors:** Zian Liu, Md Abul Hassan Samee

**Affiliations:** Department of Integrative Physiology, Baylor College of Medicine, TX, US

## Abstract

DNA methylation describes the addition of methyl groups, often between CpG dinucleotides. Single-cell bisulfite sequencing technologies allow the measurement of DNA methylation levels within individual cells. Epigenetic clocks are statistical models for computing biological age from DNA methylation levels, and have been used for detecting age variations in various disease contexts. However, there have been no attempts to apply epigenetic clocks to single-cell methylation data in humans. Thus, we questioned whether pre-built epigenetic clocks could be applied to single-cell methylation data; if so, how can we perform data quality control and imputation. We concluded that 1) linear regression-based epigenetic clocks can be applied to bisulfite-sequencing data, 2) data quality control can be used to reach the desired level of prediction accuracy, 3) first-principle imputation strategies could be used for missing data on selected CpG methylation sites, and 4) machine learning-based imputation tools could be used for accuracy-based age predictions. We built the first training-free, reference data-free framework for estimating epigenetic age in human single-cells, which would provide a foundation for future single-cell methylation-based age analyses.

## Introduction

DNA methylation (“DNAm”) denotes the addition of methyl-groups to DNA nucleotides, commonly the cytosine in C-G dinucleotides (CpG sites, CpG loci) (1). DNAm is highly influential on gene regulation and developmental biology, and has been shown to differ under the context of various genetic diseases (1–3). Intriguingly, DNAm has also been used for accurately predicting biological ages of tissue samples (4,5). Approaches for measuring DNAm include (a) sequencing-based approaches, such as whole-genome bisulfite sequencing (WGBS-seq or WGBS) and reduced-representation bisulfite sequencing (RRBS), and (b) array-based approaches, such as Illumina 27k/450k/EPIC arrays. Sequencing approaches provide extensive genomic coverage and are considered to be the “gold standard” (6), but are cost-prohibitive and suffer from limitations in either read-depth and bisulfite treatment-related biases (WGBS) or genome coverage (RRBS). Array-based approaches are affordable, target a pre-selected set of loci from the entire human genome, and result in accurate methylation level measurements at said loci, but assess fewer CpG loci (up to 850k in the current state-of-the-art method) in total. Single-cell DNAm technologies (scMe), such as single-cell bisulfite sequencing (scBS-seq), single-cell RRBS (scRRBS-seq), and single-cell multiomics methods, profile epigenetic alterations at single-cell resolutions with up to whole genome coverage (7–10); this allows researchers to investigate epigenetic variations within tissues. However, scMe technologies also further expose the limitations of sequencing-based approaches, namely lower read depths compared to bulk BS-seq, and sparser coverage of only 5-20% of genomic methylation loci (7–10). Despite recent advances in single-cell methylation technologies, no current method provides both high genome coverage and high read depth at specific CpG loci (6).

Epigenetic clocks are statistical models for predicting a tissue sample’s biological age, or “epigenetic age”, from DNAm values of specific CpG loci (5). These models have been used to accurately predict chronological age in healthy individuals (4,5,11–17). They have also been used reveal disease-related deviations in biological ages, such as increased variation in biological age as seen in tumor samples (4), and elevated biological age (“age acceleration”) as seen in HIV patient samples (18). Several clock models also function as “pan-tissue clocks” that can make reasonably consistent age predictions across many tissue types (4,14,16,17). Recent clock models have attempted to improve the accuracy of epigenetic age predictions by incorporating different CpG loci and/or employing model architectures beyond regularized linear regression (14,15,17,19). See Table 1 for a summary of previously published epigenetic clock models for humans. Hence, epigenetic age has evolved as an independent means of studying the influence of methylation variation, between either different tissues or different individuals.

**Table 1.** Comparison of published methods for human epigenetic age estimation.

| **Method name**<br>**or alias** | **Publication** | **Model**<br>**architecture** | **Data type** | **Additional**<br>**training**<br>**required** | **Non-**<br>**methylation**<br>**data required** |
| --- | --- | --- | --- | --- | --- |
| Hannum | Hannum et al.<br>2013 | Linear<br>regression | Illumina 27k | No | No |
| Horvath | Horvath 2013 | Linear<br>regression | Illumina 27k | No | No |
| Horvath2 | Horvath et al.<br>2018 | Linear<br>regression | Illumina<br>27k/450k | No | No |
| Zhang19 | Zhang et al.<br>2019 | Linear<br>regression | Illumina 450k | No | No |
| RnBeads | Assernov et al.<br>2014 | Linear<br>regression | BS-seq | No | No |
| PhenoAge | Levine et al.<br>2018 | Generalized<br>linear model | Illumina | No | Yes |
| MethylNet | Levy et al.<br>2020 | Variational<br>autoencoder | Illumina * | Yes | No |
| AltumAge | de Lima<br>Camillo et al.<br>2021 | Deep neural<br>network | Illumina<br>27k/450k | No | No |

Recent advances in scMe and epigenetic clocks lead us to ask whether single-cell methylation age analyses could be used to study age variations within tissues, such as for mammalian development and various cancer subtypes (6,20). However, such an analysis has never been attempted on human data, likely due to the following two reasons. First, it has been suggested that up to thousands of samples are required to train an accurate epigenetic clock model (5), but there is no such large-scale BS-seq or scBS-seq data available for humans. Although BS-seq-based epigenetic clocks and single-cell methylation analysis frameworks have been developed in mice (21,22), there is currently no equivalent method in humans. Second, the inherent data quality limitations to scBS-seq imply that scBS-seq and array-based DNAm data may have different structures, so models built on the latter may not be directly applicable to scBS-seq. Namely, scBS-seq data have highly variable coverage of up to only 5-20% of CpG loci across the genome, low read-depth at CpG loci with coverage, as well as biases caused by the harsh bisulfite treatment (6,23). These factors make it challenging to use scBS-seq data for epigenetic clock analyses without additional data quality control and imputation.

Multiple first-principle imputation strategies (4,8,14) and several machine learning (ML)-based imputation methods (24–28) have been used for DNAm imputation. However, the efficacy of these methods has not been tested for making epigenetic age predictions. Given the innate data quality issues observed in scBS-seq (6,8,10,23) and the lack of appropriate methods for data imputation prior to making epigenetic age predictions, it may be advantageous to preprocess the data by filtering out low-quality samples, imputing missing data in the remaining samples, and then apply pre-trained epigenetic clock models on them.

Considering the aforementioned lack of appropriate epigenetic clock models and limitations in scBS-seq data, we first asked whether it is possible to apply epigenetic age analyses to scBS-seq data. If so, we aim to establish quality control guidelines which allows the production of robust outcomes from scBS-seq data, as well as the tools or pipelines that may be necessary to aid such analysis. We have, to our knowledge, built the first pipeline for the application of epigenetic clocks to scBS-seq data, built the first benchmarking study to quantitatively assess quality control standards for scBS-seq data in humans, and evaluated performances of first-principle scBS-seq imputation methods in a setting for predicting epigenetic age. Our benchmarking results suggest that age analyses could be performed on scBS-seq data with appropriate imputation and quality control steps. It is feasible to impute the lowest quality samples within a dataset to ensure the remaining data achieves the desired level of quality standard.

## Results

### A pipeline for simulating scBS-seq data and evaluating the efficacy of state-of-the-art imputation methods

We are aware of the following issues pertinent to any scBS-seq data: 1) lack of coverage or sequencing reads at large numbers of genomic locations, 2) low sequencing read depth, and 3) bisulfite conversion errors (8,23). Our initial goal is to gauge whether and to what extent such issues affect the conclusions of a single-cell epigenetic age study (see below). Then, we want to assess if we can make up for the loss in performance via data quality control and imputation. Finally, we want to develop a workflow for conducting quality control in real scBS-seq data (Figure 1A).

**Figure 1.**
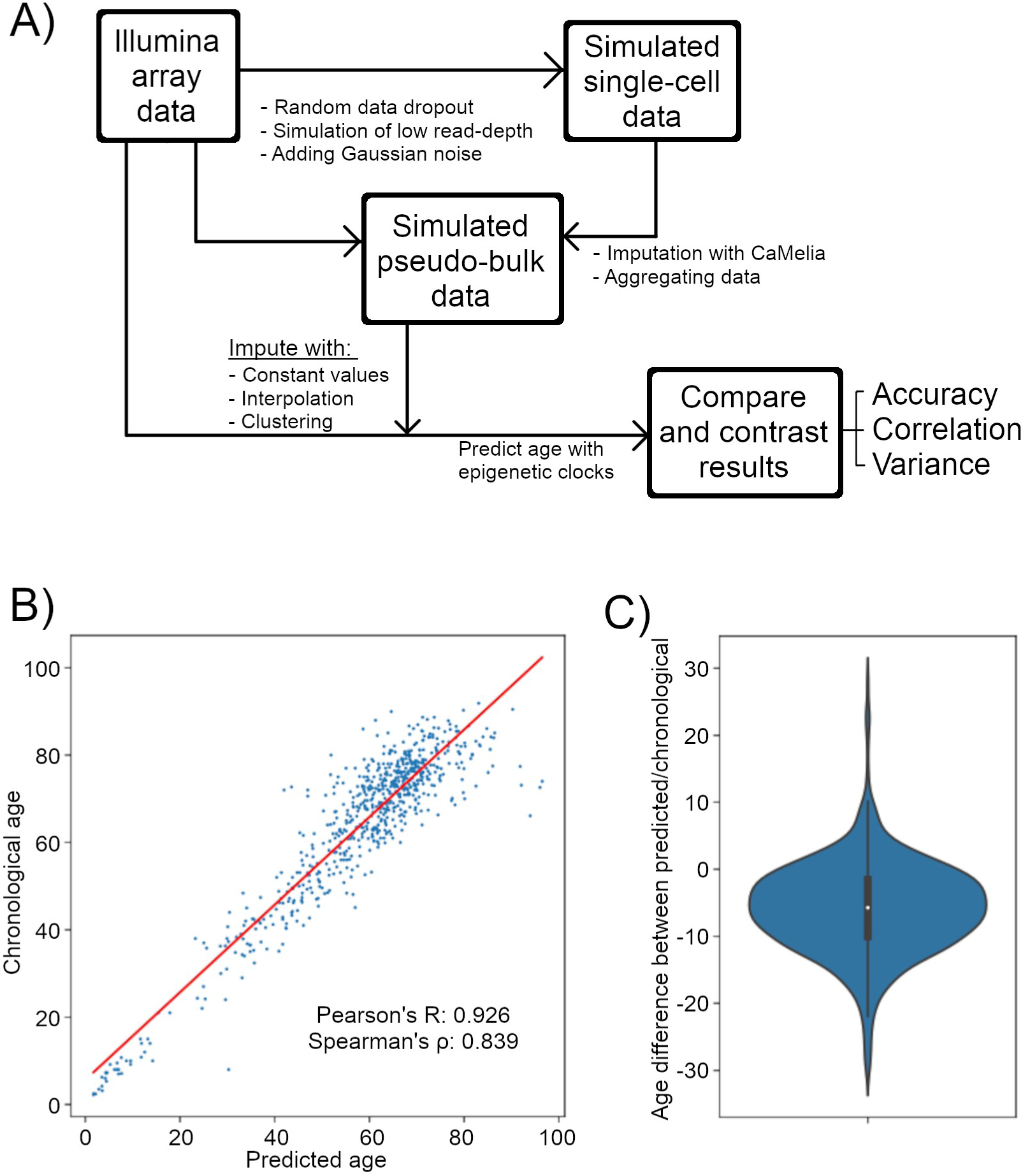
Overview of our study. A) Overall workflow of our benchmarking project. B) Scatterplot and C) violin plot showing results from applying the Horvath clock to the unperturbed Illumina array input data. In B), the x-axis shows age predictions made by the epigenetic clock, the y-axis shows observed chronological/calendar age of samples, and the label shows Pearson/Spearman correlation values between the two axes. In C), the y-axis shows per-sample differences between the predicted and observed age values.

Researchers may be interested in pursuing any of the following hypotheses regarding epigenetic age prediction: 1) to accurately predict the biological age of samples, 2) to compare differences in epigenetic age, or “age accelerations”, between two treatment conditions, or 3) to compare variations in biological age between two treatment conditions. The first and the second hypotheses have been pursued in multiple aging research projects (4,18,29–31). The third hypothesis as a phenomenon has been documented in bulk tumor samples (4,31) and may be of interest for researchers studying heterogeneous diseases. These three sets of hypotheses would require different model performance metrics: 1) high accuracy between predicted results and chronological age values, 2) accurate correlative trends between predicted results and chronological age values, or 3) low variation between predicted results of single-cells of similar chronological age values. We will refer to the performance of the epigenetic clock across all three metrics as model “performance”. The individual metrics will be referred to by the measurements we use, namely “mean/accuracy”, “Pearson correlation”, and “variance” (Figure 1A).

To overcome the data quality issues of scBS-seq data, we aggregated read counts in individual single-cells to create “pseudo-bulk” BS-seq data. Although it may appear preferable to analyze data on a per-cell basis, our analysis of the commonly used scBS-seq and scRRBS-seq dataset suggest that scMe data from individual cells experience extreme data sparsity and low read depth (Figure S1-S2), which can be mitigated to a large extent by aggregating cells into pseudo-bulks. In principle, performing a pseudo-bulk aggregation will ensure 1) more CpG loci to have at least one sequencing read, 2) CpG loci that are commonly sequenced across multiple cells will have increased read-depth, and 3) noise caused by bisulfite conversion errors is minimized. An outline of our approach is as follows. We first select previously generated Illumina array datasets from healthy individuals (32). We then apply the original Horvath clock, a popular pan-tissue epigenetic clock using only 353 CpG loci (4), to the unperturbed data (Figure 1B-C). Then, we apply three strategies to perturb the input data so that they display properties of pseudo-bulk scBS-seq data, and perform grid-search combinations of our three perturbation methods to test for negative effects of perturbation on the pseudo-bulk data.

Most epigenetic clocks are linear models that do not tolerate missing inputs, so they would require data imputation. We have selected several first-principle data imputation approaches based on 1) the relatively stable methylation states of most DNAm sites (33), 2) prior publications related to DNAm imputation strategies and tools (24–28,34), and 3) methods used by published epigenetic clocks (4,14). We also included a recently published ML tool (27) for data imputation. We first compared various first-principle imputation methods for recovering from data perturbation effects. Then, we simulated a single-cell dataset on which we applied both cutting-edge ML imputation tools and our best-performing first-principle imputation strategies, re-created the pseudo-bulk data, and compared the performances between these approaches. See Figure 1A for our workflow and the Methods section for details on our approach. See the Methods section for more rationale behind our selection of data processing methods.

### Data sparsity, read depth, and noise adversely affect age prediction performance

We started our analysis with a reference DNA methylation Illumina 450k array-based blood sample data from 691 healthy individuals aged between 2 and 92 years (Figure S3) (32). We perturbed the input data so that data from each sample would resemble an aggregate of methylation data from individual cells, or “pseudo-bulks”. We first performed data dropout by removing a predetermined portion of randomly selected data across all samples and loci. We then simulated low read-depth by using the original data as binomial distribution priors, where we performed N binomial simulations and recorded the outputs as synthetically generated data. To simulate a more realistic distribution of read-depth values, we also obtained read-depth information from a set of pseudo-bulk data generated from a recent scTrio-seq dataset (35). Based on read-depth distributions in our dataset of interest (Figure S4), we described a list of nine read-depth distributions that recapitulate read-depth distributions of several pseudo-bulk samples (Figure S5, Table S1). Finally, we simulated bisulfite conversion error by introducing Gaussian noise with fixed standard deviation values as suggested by a previous publication (25). See Figure 2A for a graphical illustration and the Methods section for more details.

**Figure 2.**
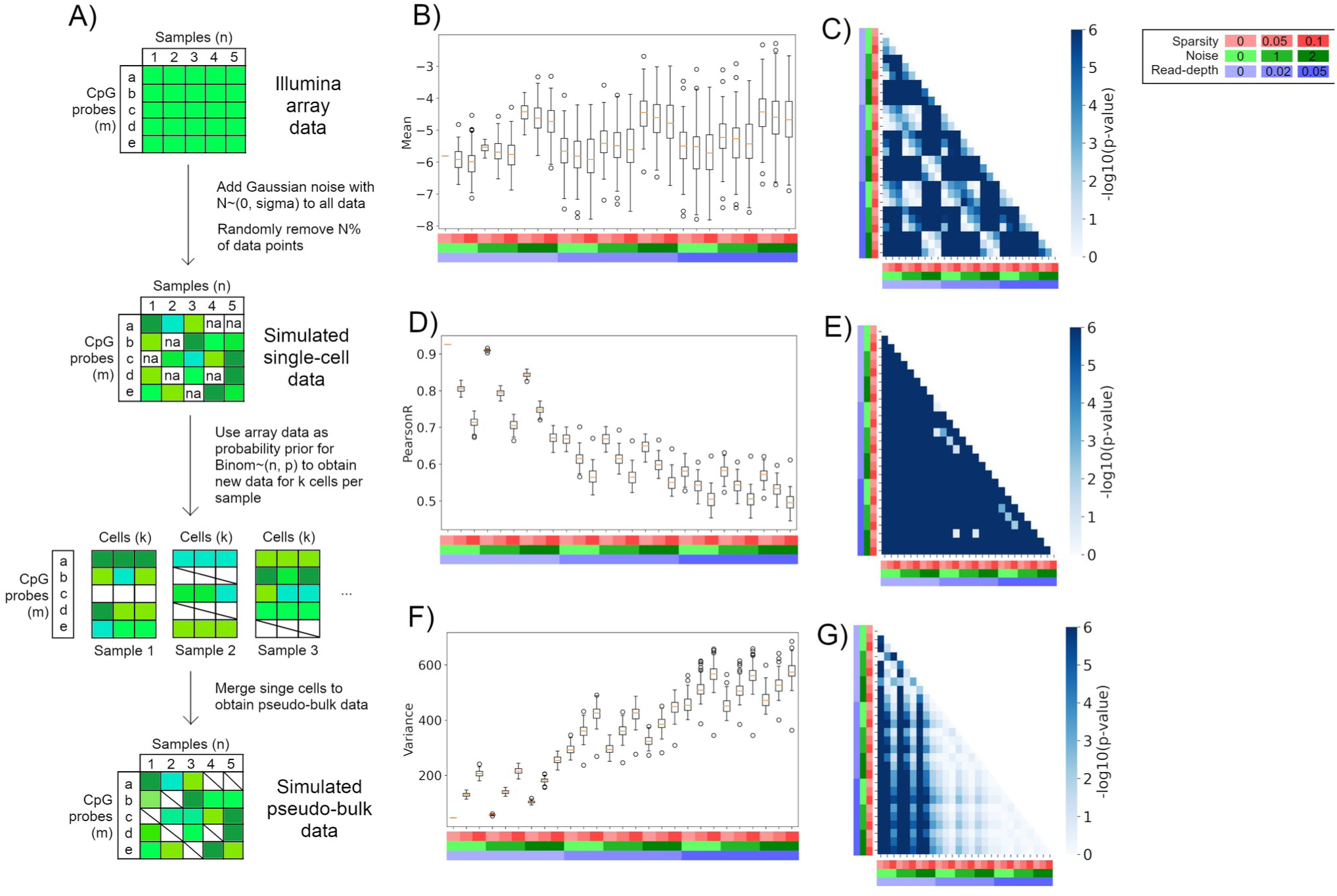
Data perturbation negatively influences prediction quality of epigenetic clocks. A) Graphical overview of our data perturbation strategies as shown on a data matrix with n=5 samples (from different individuals) and an epigenetic clock using 5 CpG loci. The colored blocks represent methylation values, with white/empty blocks representing missing data. B-G) Prediction results from combining data dropout, lowered read-depth, and bisulfite conversion error constraints, with different color combinations representing combinations of data perturbation as seen in the legend. The heatmaps show -log10-transformed p-values in the range of [0, 6], or p=1 to p=1e-6. P-values lower than 1e-6 are capped. B-C) Prediction accuracy/mean as shown by B) box plots or C) pairwise t-tests. D-E) Prediction correlation/Pearson’s p as shown by D) box plots or E) pairwise t-tests. F-G) Prediction precision/variance as shown by F) box plots or G) pairwise F-tests.

Increasing data dropout proportion systematically reduces age prediction performances (Figure S6-S11, Table S2). This is especially the case for mean and Pearson correlation predictions, where high dropout levels result in significantly different model predictions compared to low dropout values (Figure S6-S9, Table S2). For variance predictions, data perturbations significantly reduce predictive qualities but bottom out at ∼20% data dropout (Figure S10-S11, Table S2).

Increasing Gaussian noise also systematically reduces age prediction performances (Figure S12-S17, Table S3). Adding any level of noise significantly influences mean and correlation predictions, but the absolute differences in mean and correlation predictions are not strongly influenced until a standard deviation of 0.02 (Figure S12-S15, Table S3). On the other hand, adding a noise of less than or equal to a standard deviation of 0.05 did not significantly influence variance prediction (Figure S16-S17). Previous work has estimated total bisulfite conversion errors in standard BS-seq to reach ∼0.86% in high-accuracy conditions (23), so our simulated errors may be more rigid than real-world conditions. The influence of bisulfite conversion errors on age prediction may be negligible in real data.

Decreasing read-depth levels also systematically reduces age prediction performances (Figure S18-S29, Table S4-S5), which is the case for both constant read-depth limitations (Figure S18-S23, Table S4) and data-derived read-depth limitations (Figure S24-S29, Table S5). Having a uniform read depth above 20 may have little adverse effects on model performance (Figure S18-S23, Table S4). Similarly, using realistic read depth distributions also significantly adversely influenced model prediction except for having the most lenient read depth settings (Figure S24-S29, Table S5).

Finally, we performed a grid combination of the three types of data perturbations introduced above (Figure 2A, Table S6). Each combination of data perturbation reduces the prediction performances of our selected model in terms of average prediction (Fig 2B-C), correlation (Fig 2D-E), and precision (Fig 2F-G), and higher levels of perturbation result in systematically reduced prediction performances as expected (Fig 2B-G, Table S6). Specifically, the perturbations rarely significantly influence the mean of average predictions, but tend to make predictions much farther from the mean (Fig 2B-C), whereas correlation and variance systematically decrease prediction performances (Fig 2E, F) with most levels of perturbations being significantly different from each other (Fig 2E, G).

We have implemented the above framework as a modular pipeline, which makes it possible to input any linear epigenetic clock of interest and a set of quality standards and obtain results on how the clock model would perform given these metrics. The generated results could then be used for data quality control purposes.

### Simulating quality standards using a data-centric approach allows the identification of specific low-quality data to remove

Aside from conducting quality control via a pre-determined cutoff, it is also possible to gradually remove or penalize the lowest-quality samples until the remaining samples reach a desired quality-control level. This approach is particularly useful when specific samples have low quality or are outliers, and has been performed in tasks such as RNA-seq outlier detection (36,37). To pursue this direction, we have devised a testing framework to identify samples with the lowest quality levels, where we 1) select a set of reference samples from the bulk data, 2) derive a set of quality standards from our scBS-seq data-of-interest, then 3) apply the quality standards to the reference samples and evaluate prediction results. We could then compare our performances on the simulated bulk dataset, and check how many low-quality samples need to be removed before the data reaches a desirable level of prediction performance. Since this approach allows us to evaluate the performances of epigenetic clocks on samples with the same quality levels as our data-of-interest, albeit with the ground truth values available, we will refer to this as the “data-centric approach”. See Figure 3A for a graphical illustration.

**Figure 3.**
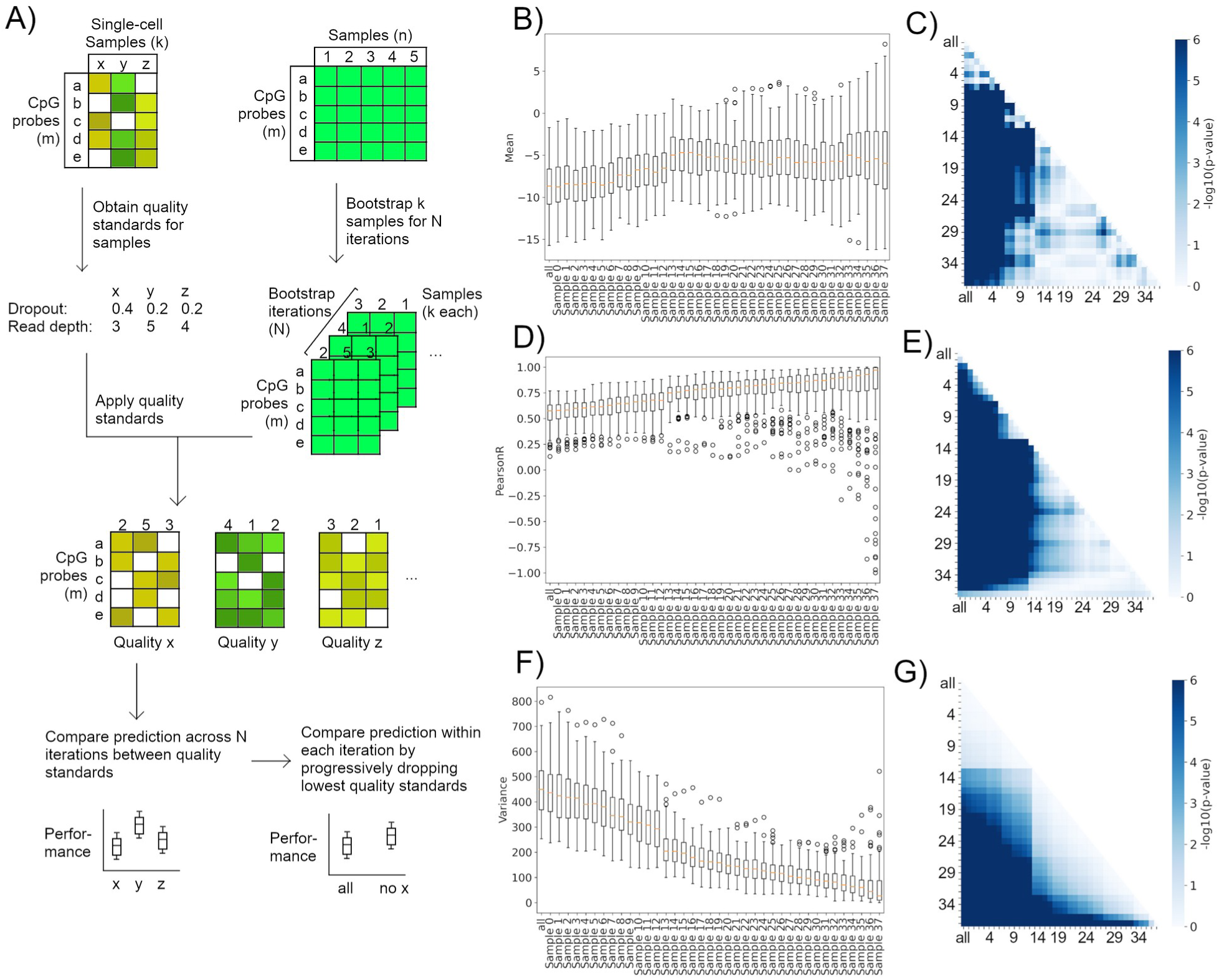
Data-centric quality control reveals optimal number of samples to remove to achieve quality standard. A) Graphical overview of our data-centric quality control strategies as shown on a data matrix with n=5 samples (from different individuals) and an epigenetic clock using 5 CpG loci, as well as three scBS-seq pseudo-bulk samples of various quality standards. The colored blocks represent methylation values, with white/empty blocks representing missing data. B-G) Prediction results from systematically removing the lowest quality samples as measured from high to low variance, with different labels representing the last sample being dropped from the data. The heatmaps show -log10-transformed p-values in the range of [0, 6], or p=1 to p=1e-6. P-values lower than 1e-6 are capped. B-C) Prediction accuracy/mean as shown by B) box plots or C) pairwise t-tests. D-E) Prediction correlation/Pearson’s p as shown by D) box plots or E) pairwise t-tests. F-G) Prediction precision/variance as shown by F) box plots or G) pairwise F-tests.

We have performed our analysis as described, and have progressively removed the worst quality standards, as ranked by 1) variance, 2) Pearson’s correlation, or 3) random selection as control (Table S7). Our rationale is that a low-quality sample would lead to either low prediction correlation with the underlying chronological age or high variability. We did not rank the quality standards by prediction accuracy, as our previous results suggest that data perturbation does not usually influence prediction performance in a statistically significant manner (Figure 2B-C).

Removing low-quality standards by ranking variance or Pearson’s correlation both produced progressively stabilized mean prediction (Fig 3B-C, Figure S30-S31) and progressive, statistically significant Pearson correlation and variance improvements (Fig 3D-G, Fig S32-35, Table S7-8). This is in contrast to removing quality standards by random selection, where we see no significant performance improvements in any of the three metrics (Figure S36-41, Table S9).

Interestingly, removing quality standards ranked by variance also produced statistically significant “blocks”. In the case of our real data-derived quality standards, removing one-third of the most variable standards stabilized prediction mean and correlation such that further reduction of standards did not produce statistically significant differences (Fig 3C, E). Ranking by correlation did not produce such a “statistical significance block” effect (Figure S30-35). This concept may be informative of the optimal number of samples to retain for downstream analyses. For example, if we have 20 pseudo-bulk scBS-seq samples-of-interest, and simulations suggest that removing up to 6 quality standards statistically significantly improves prediction performance but further removal of quality standards does not, then we will remove those 6 samples and retain the remaining 14 for downstream analyses.

We also included tabulated formats of the results (Table S7-10). The data-centric approach makes it possible to specify the desired level of prediction accuracy, correlation, and variance, and then progressively remove the lowest-quality samples until the remaining simulated samples reach the pre-determined prediction results.

### Comparing imputation strategies suggest linear interpolation, clustering, and machine learning tools to be effective at imputing data for different hypotheses

We have compared various first-principle-based imputation strategies across randomly perturbed data with three levels of perturbation: light, medium, or heavy (Figure 4A, Table S11). Our selected imputation strategies include imputing with constant values, imputing with the average methylation level across the sample or the loci, interpolating values by building a local regression using methylation levels of nearest neighboring CpG loci, or clustering samples based on methylation levels and imputing with the average methylation level across samples in the same cluster. We have explained our rationale for the choice of imputation methods in the Methods section, and we have summarized the results in both graphical and tabulated formats (Figure 4, Figure S42-53). The full list of imputation strategies is tabulated in Table 2.

**Figure 4.**
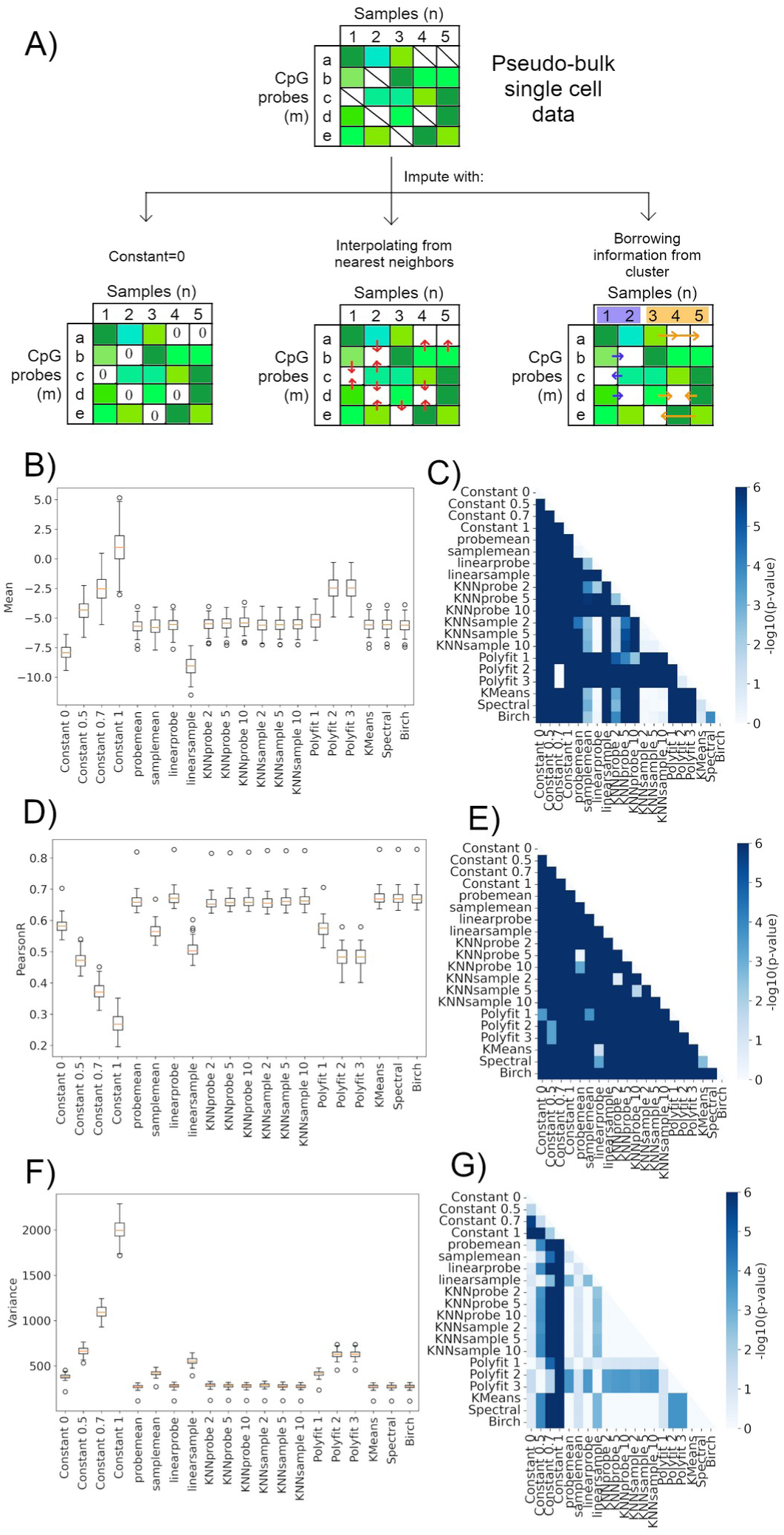
First-principle imputation strategies rescue data with similar performances aside from a few notable exceptions. A) Graphical overview of the application our first-principle-based data imputation strategies as shown on a perturbed data matrix generated from previous steps, with n=5 samples (from different individuals) and an epigenetic clock using 5 CpG loci. The colored blocks represent methylation values, with white/empty blocks representing missing data. In the clustering example, samples 1-2 are categorized as the blue cluster, while samples 3-5 are categorized as the orange cluster. B-G) Prediction results as applied on the lightly perturbed dataset, with different labels representing different data imputation strategies. The heatmaps show -log10-transformed p-values in the range of [0, 6], or p=1 to p=1e-6. P-values lower than 1e-6 are capped. B-C) Prediction accuracy/mean as shown by B) bar charts or C) pairwise t-tests. D-E) Prediction correlation/Pearson’s p as shown by D) bar charts or E) pairwise t-tests. F-G) Prediction precision/variance as shown by F) bar charts or G) pairwise F-tests.

**Table 2.**
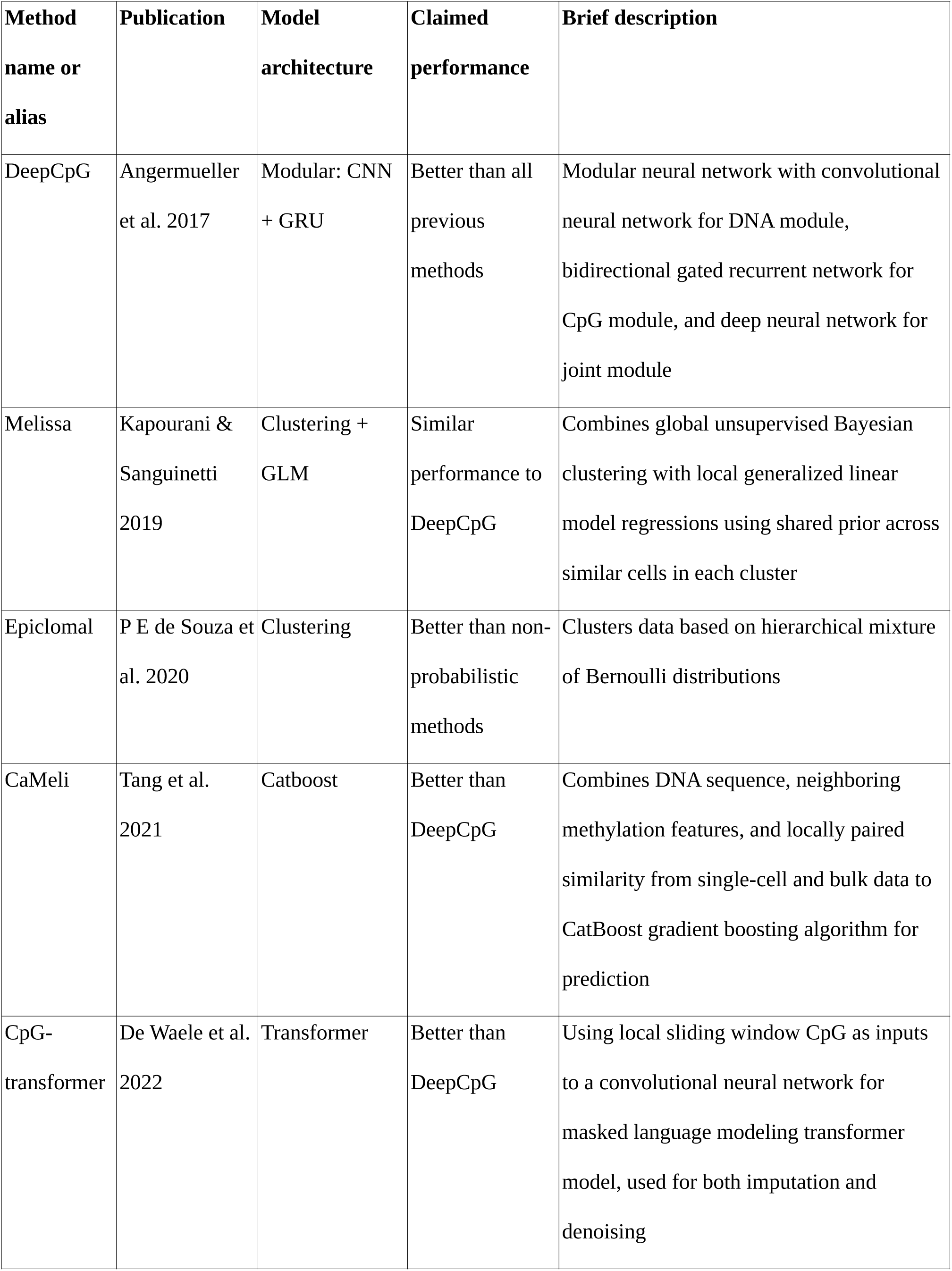
Comparison of published methods for single-cell methylation data imputation.

We have noticed wide gaps in prediction performances of several imputation methods (Figure 4B-G, Figure S42-53, Table S12), especially within the higher-quality datasets (Figure 4B-G). Specifically, the constant imputation methods, polynomial function methods, and across-sample averaging methods (“samplemean” and “linearsample”) show statistically significantly worse performance compared to all other methods. Other methods reached similar performances across all three metrics (Fig 4B, D, F). Some of these similar methods showed statistically significant differences in Pearson correlation values from each other (Fig 4E), but not in accuracy or variance (Fig 4C, G).

We have further inspected the numerical results to find the best-performing methods (Table S12). According to the results, the best-performing methods for predicting correlation are “linear probe” which averages the methylation levels of the two nearest CpG loci without missing values within a particular sample, closely followed by the clustering approaches (“K-means”, “Spectral”, “Birch”, “Ward”). The non-K-means clustering methods outperformed “linear probe” when data quality is poor (Figure S48-53, Table S12). The best-performing methods for predicting variance are the various clustering approaches along with “probe mean” which averages the methylation levels of all non-NaN CpG loci within a particular sample (Table S12). Overall, it appears that Spectral clustering (“Spectral”) and hierarchical clustering (“Birch”, “Ward”) produced the most robust results across all data perturbation qualities (Table S12). Nevertheless, these differences may be minimal as performances of most methods did not show statistically significant differences (Fig 4C, E, G).

Interestingly, the polynomial function methods performed poorly, but reached equivalent performance with “linear probe” when genomic distances between CpG loci are considered (Table S12). This is somewhat surprising, as prior literature have suggested methylation levels at a particular CpG locus to be strongly influenced by methylation levels of neighboring loci (34). We believe the poor performances of the polynomial function methods are likely due to our dataset being limited to 353 loci which are far from each other in terms of genomic locations. This limits our ability to build an accurate local basis function without taking distance information into account. This also may explain why performances of polynomial function imputation tools become comparable to other methods when genomic distances are considered.

The constant imputation methods are some of the earliest adopted imputation methods for survey data, and are, for our purposes, applied based on the assumption that most CpG loci are stably methylated (33). Constant imputation methods performed statistically significantly worse compared to nearly all other methods under most circumstances (Fig 4B-G, Table S12). Nevertheless, should it be necessary to employ them, it appears that imputing with the value 0 works best for preserving prediction correlation and variance (Fig 4D, F, Table S12), while imputing with 0.5 may be best for preserving prediction accuracy (Fig 4B, Table S12). In the context of linear regressions, imputing with 0 implies “removing” a particular predictor (in this case, CpG loci) completely, as the imputed loci will no longer contribute to the final predicted age. Imputing with 0.5 may approximate the average methylation levels across all CpG loci (33). However, considering that imputing with any of the constant values significantly skewed either prediction accuracy, Pearson correlation, or variance (Figure 4B-G, Table S12), imputing with a constant should only be used as a last resort and will likely not result in high prediction performances.

Finally, we compared our first-principle-based imputation methods with CaMelia (27), a recently published scMe imputation tool that has outformed other state-of-the-art models like DeepCpG. To ensure valid comparisons, we created synthetic datasets of 25 single cells each without creating pseudo-bulks following our outlined approach (Figure 1A), performed CaMelia imputation on these data, and compared age prediction results between using or not using CaMelia. Since CaMelia was unable to impute all missing CpG loci, we used secondary, first-principle imputation methods for the remaining missing loci.

The results suggest that applying CaMelia rescued certain CpG loci with zero sequencing reads (Figure 5A), and increased read depth at the majority of CpG loci (Figure 5B) as expected. However, data rescued with CaMelia showed mixed results in age prediction. Although CaMelia improved prediction accuracy by reducing the differences between predicted and chronological age (Figure 5C, D, Table S13), it also reduced prediction Pearson correlation and increased prediction variance between predicted and chronological age (Figure 5E, F, Table S13). The samples we selected for testing CaMelia are representative of the broader age distribution observed across all used samples (Figure S54), and performance trends between the other, non-CaMelia methods are consistent with our imputation results above (Figure 4, Figure 5C-F, Table S13), suggesting that the results are likely not statistical artifacts. It appears that the usage of CaMelia may be best suited for situations when obtaining an accurate proxy for chronological age is desired.

**Figure 5.**
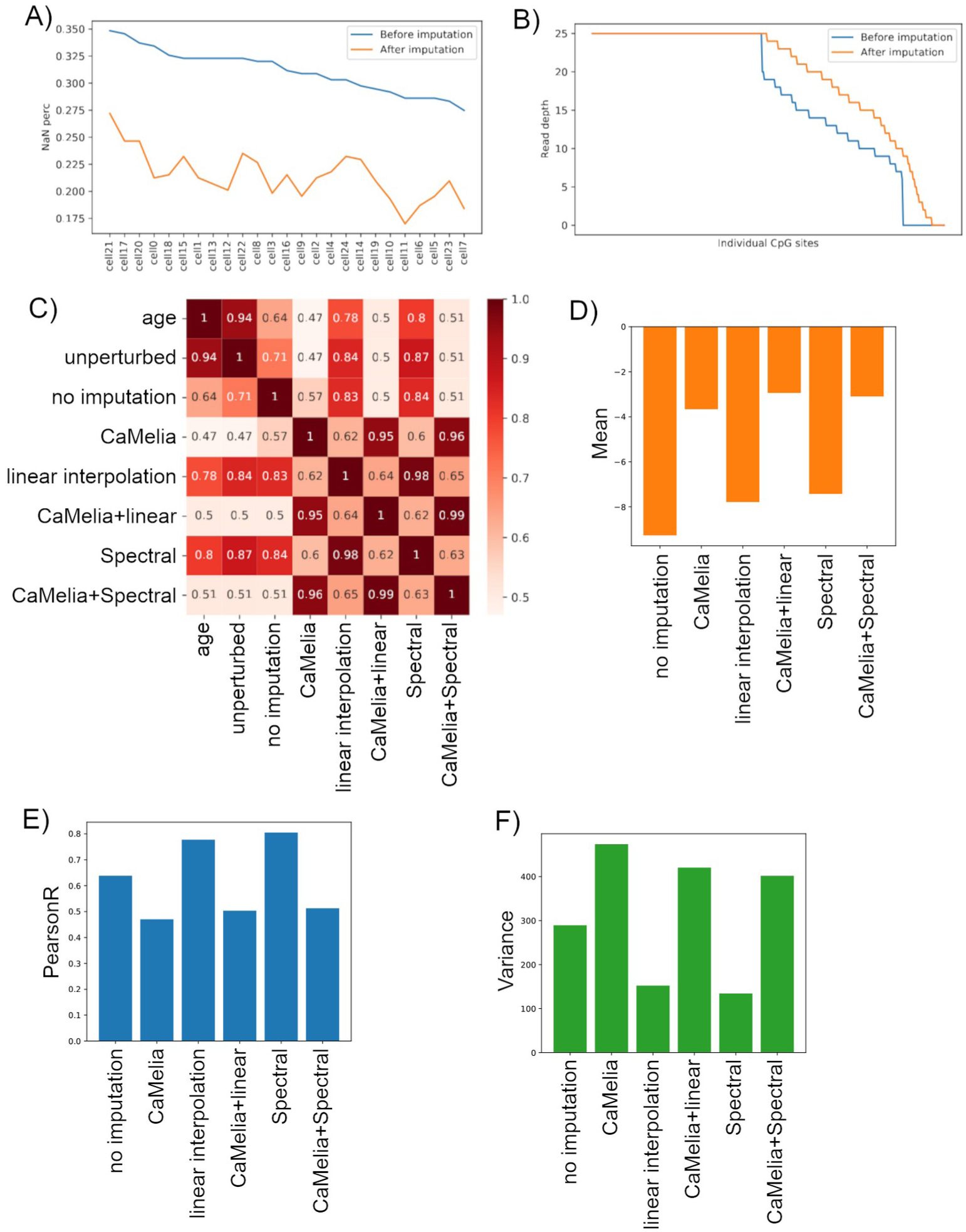
Imputing with CaMelia showed mixed results compared to first-principle imputation strategies. A-B) Lineplots comparing data qualities of single-cells before and after CaMelia imputation in an exemplar sample. A) Percentage of CpG loci with missing values within each cell, B) total number of reads of each CpG loci summed across all cells. C-F) Heatmap showing pairwise Pearson correlation values between chronological age, age predictions on unperturbed data, and three imputation approaches (no imputation or impute with 0, linear interpolation, Spectral clustering) with or without CaMelia imputation. D-F) Detailed comparisons between the aforementioned three imputation approaches combined with or without CaMelia in terms of D) prediction mean, E) prediction Pearson’s correlation, or F) prediction variance.

### Data guidelines for epigenetic age prediction with scBS-seq data

Based on our results, we would recommend the following guidelines for making epigenetic age predictions with scBS-seq data. See Figure 6 for a flowchart of our conclusions.

**Figure 6.**
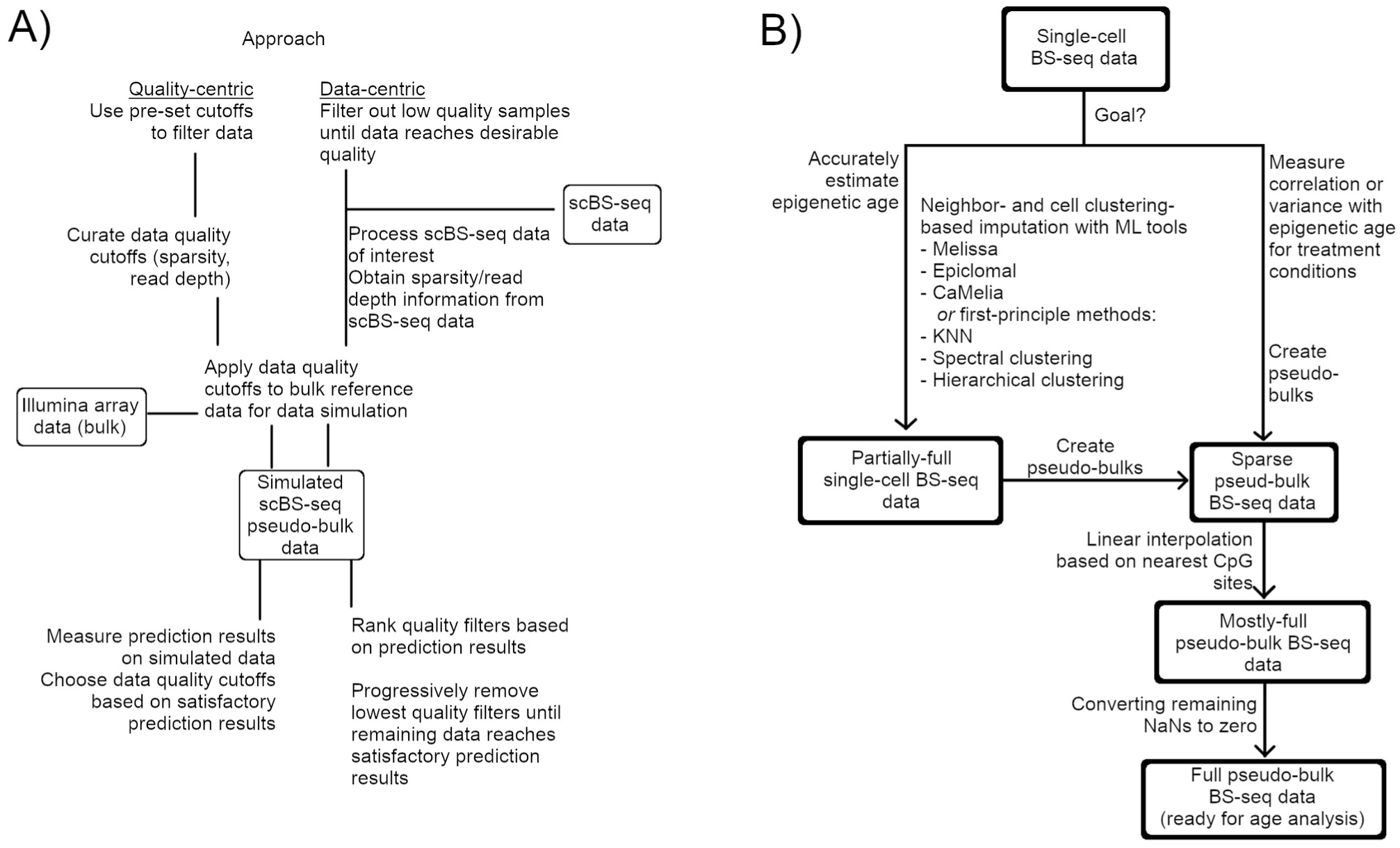
Our data processing recommendations summarized from our findings. A) Our recommended approach for single-cell data quality control. B) Our recommended approach for scBS-seq data preprocessing for epigenetic age analyses.

#### 1. Rule of thumb for data quality-control

At the current stage of development for single-cell technology and epigenetic clocks, we recommend creating pseudo-bulks of single cells before performing age predictions. Nevertheless, single-cell technologies still provide the advantage of being able to combine cells into different pseudo-bulks which best suit a researcher’s needs.

Our results from perturbations in sparsity, read depth, and bisulfite conversion individually for the Horvath 2013 model (Table S6), as well as from first-principle data imputation strategies (Table S12), can be used as guidelines for quality-control cutoff values specifically for the Horvath 2013 clock. We provided a script for estimating such results for different linear epigenetic clock models. Alternatively, we also provided a script for using a data-centric approach to remove the worst-quality samples within a batch.

#### 2. Strategies to impute data

Our data suggest that first-principle-based approaches alone are sufficient in creating adequate prediction accuracy. Two of the simplest yet reasonably accurate imputation methods are to 1) perform a linear interpolation by averaging the methylation levels of the closest, non-missing upstream and downstream methylation loci, or 2) use Spectral or hierarchical clustering across samples. There is no guarantee that the recently developed imputation tools can be performed for every single CpG loci of interest, so first-principle approaches may be necessary as a fallback regardless of other imputation strategies used.

#### 3. Validity of currently available imputation methods for age prediction

DeepCpG, Melissa, Epiclomal, CaMelia, and cpg-transformer (24–28) are currently available tools for imputing methylation data. We did not benchmark the performances of DeepCpG in our study. As explained by previous papers (25), DeepCpG 1) necessitates having full CpG methylation information of up to 25 neighbors, and 2) is much more computationally heavy than other listed methods. These two conditions make DeepCpG an unlikely choice for data imputation given our current task. It is also worth noting that several of the methods have reached similar or higher performance than DeepCpG in their respective benchmarking results (25–28). Since the remaining methods utilize very similar approaches, we selected one of the newer methods, CaMelia, for our benchmarking study.

## Discussions

The advent of scBS-seq and related technologies, combined with the availability of epigenetic clock models, have inspired us to test the feasibility of performing epigenetic age estimations at single-cell resolutions. We have demonstrated that it is possible to perform epigenetic age analyses on scBS-seq data, without the use of extra data modalities or model training, by using linear regression-based epigenetic clocks with first-principle imputation tools. Our work suggests that scBS-seq data quality control can be done by either performing a routine test using pre-defined quality standards, or by defining quality standards from our data of interest and testing them individually on a bulk reference dataset. We suggested specific first-principle imputation tools that are best suited for imputing DNAm data for age predictions, as well as the fact that ML-based tools could be useful for accuracy-based predictions but not other goals.

We hope our framework could be applied to scenarios where epigenetic age estimates could bring novel insights into the underlying biology of diseases. For example, a researcher may be interested in performing a longitudinal age analysis across patients at multiple age points, assessing accelerated aging patterns within specific cell types in pediatric heart tissues, or comparing epigenetic age variation between two tumor subtypes. Such an analysis would be impossible with the conventional epigenetic clock models as they do not work directly with single-cell datasets. In contrast, a single-cell epigenetic age estimation framework with high performance in terms of accuracy, correlation, and variance would be able to help power such analyses.

### The necessity of imputation and model training

We have assumed that imputation is necessary for epigenetic age predictions, as linear regression models necessitate non-sparse input data. From another perspective, it is possible to avoid or limit imputation by selecting loci with high coverages within a dataset, and use those loci to train a novel epigenetic clock. We did not consider this as an option for the scope of this manuscript, as it has been suggested that training an accurate epigenetic clock requires thousands of samples, with recent clocks utilizing over 10,000 training samples (5,14,17). In contrast, a human pathological dataset may have only up to hundreds of single-cells for the adjacent normal control, and even fewer when aggregated to pseudo-bulks. Nevertheless, the argument for using thousands of training samples assumes that a conventional linear regression framework is used to train models. It is possible that a statistically rigorous framework could be used to train an accurate model even with limited training data, so a novel model could, in theory, largely avoid imputation by utilizing only loci which have very high coverages. This is an idea we wish to explore more in a future project.

### Comparison with scAge

Notably, a recent project, “scAge”, employed the aforementioned model training strategy to profile murine cell aging (22). Compared to our project, scAge 1) builds a novel method and thus requires model training, 2) focuses on using a minimal set of CpG loci that intersect between single-cell and bulk data, and 3) requires an external bulk-data reference. Although we are excited about scAge and its applications, the approach might not currently be feasible for human data for the following reasons. Modeling frameworks such as scAge 1) still necessitates model training, but there are few human single-cell BS-seq datasets available for such purposes; 2) requires bulk reference data, which may not be available for a human dataset of interest. Instead, we focused our strategies on imputing data at loci provided by previously validated epigenetic clocks, which allows us to build an age prediction pipeline that is training-free and reference-free.

### Availability of BS-seq-specific epigenetic clocks for humans

BS-seq-specific epigenetic clocks have been available for mice (21), but there is only one human BS-seq-specific epigenetic clock available as part of the RnBeads package as far as we are aware of (38,39). Horvath and Raj have suggested the importance of having large training data sample sizes, up to thousands of samples, for building an accurate model (5), so the lack of human BS-seq-specific epigenetic clock is likely caused by the limited amounts of human BS-seq data available. Namely, the aforementioned RnBeads clock only used data from 232 samples for training and did not perform model validation (38,39). At the current stage, we do not recommend training or using BS-seq-specific epigenetic clocks for human epigenetic age prediction.

### Usage and selection of single-cell epigenetics imputation programs

We are aware of multiple recently published methods to impute single-cell methylation data (DeepCpg, Melissa, CaMelia, cpg-transformer) (24–28). These programs differ in their ML model architectures, but all rely heavily or exclusively on the similarity of methylation states of a CpG locus of interest to its neighbors or parallels across similar samples. As described earlier, we have used CaMelia in our project (27) which not only outperformed DeepCpG in the authors’ own benchmarking but also provided an easy-to-train framework. Regarding other methods that we did not test: Melissa is designed for location-specific imputation and may fail at genome-wide tasks due to many regions having low read depths, and was thus not suitable for our purpose (25). We are also aware of Epiclomal and cpg-transformer with the latter outperforming CaMelia (26,28); we were unable to incorporate them in our work due to computational constraints, but we believe both models should show promise in practical use and would likely result in similar performances as CaMelia.

## Methods

### Selection and exclusion of epigenetic clocks

We are aware that various epigenetic clocks have been built for humans, the vast majority of which employ a linear regression or similar model training framework (4,11–17,19). We selected the Horvath 2013 model, one of the most popular pan-tissue epigenetic clock models (4), to use for this manuscript, as this model has been demonstrated to make accurate predictions across a wide range of tissue types (5), has a simple model architecture, and has relatively few (353) CpG loci. Nevertheless, our analysis pipeline is built to be directly applicable to any linear regression-based epigenetic clock model.

Regarding why we didn’t select a few other popular epigenetic clocks. The Horvath skin-and-blood clock (13) is trained only on skin and blood cells, so it is not considered for this particular project. The Hannum clock (11) has been suggested to be based on cell proportions (5), which would make it not suitable for single-cell-based analyses. Other epigenetic clocks did not reach the widespread popularity of the aforementioned clocks and were thus not considered for the main workflow of this project. Notably, newer epigenetic clocks might not use a conventional linear regression framework. As an example, AltumAge (17) uses a deep neural network, for which we would need to adapt our analysis pipeline before processing as neural networks cannot be interpreted directly via its coefficients.

To run the Horvath 2013 clock, we first downloaded clock coefficients and intercept terms from the original publication, and then programmed a matrix multiplication operation to use said coefficients/intercept in Python. We did not perform BMIQ normalization to our input data as described in the original publication.

### Selection of reported statistics

We measured 1) mean, 2) Pearson correlation, and 3) variance across all of our analyses in response to the three possible goals outlined in our manuscript. For measuring the mean, we measured the relative differences between predicted age and chronological age, and reported the results by finding the mean/average across all relative differences in each experiment. For measuring Pearson correlation, we measured correlation values between predicted age and chronological age for each experiment. For measuring variance, we first measured relative differences as in measuring mean, and then reported the variances as defined by those differences. For pairwise comparisons, we performed pairwise t-tests for 1) mean and 2) Pearson correlation, and pairwise F-tests for 3) variance.

### Additional information on pairwise comparisons

We performed pairwise one-sample t-tests or F-tests between each pair of distributions of data, which result in two heatmaps with all values across the diagonal equal to 1.

Given two data arrays x and y with lengths L each. We performed paired two-sample t-tests using the following function:

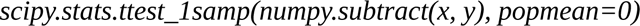

This function executes a one-sample t-test using the differences between two matched arrays. We performed F-tests using the following function:

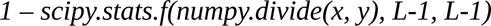

This function obtains p-values using the cumulative distribution function (cdf); the cdf is calculated by obtaining the division between two paired arrays, and then using the length of each array minus one as degrees of freedom. The used functions are implemented in the Python libraries scipy and numpy. The resulting p-values are then adjusted for multiple testing via FDR using the Benjamini-Hochberg procedure as implemented in:

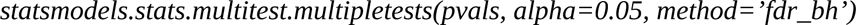

This function is from the Python library statsmodels.

It is worth noting that pairwise t-tests always result in pairwise comparison matrices that are diagonally symmetrical, but pairwise F-tests will not result in symmetrical comparison matrices. If our pairwise comparison matrices are not symmetrical, we will report the lower p-value of the two values across the diagonal.

### Constructing the reference dataset

To construct an unperturbed reference, we first concatenated and rearranged the aforementioned Illumina 450k array data, and then created a reference array containing the chronological age of each individual/patient in the data using clinical information. We then run the epigenetic clock on the native data without data perturbations. Since our pipeline applies to linear regression models, this is performed via:

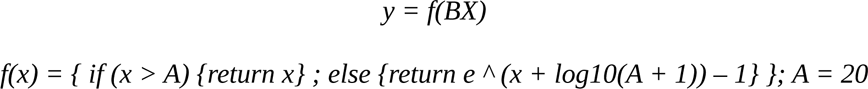

Where B is the set of linear coefficients described by each epigenetic clock model, and f(x) is a nonlinear transformation function present in certain epigenetic clocks to describe the uneven aging patterns between infants and adults (Horvath 2013, Horvath et al. 2018); note that the constant A is used to distinguish the two states of aging, and is defined as A=20 in the Horvath epigenetic clocks.

### Workflow for data perturbation analyses

We perturbed our reference dataset using various perturbation types as outlined in the following, and then predicted epigenetic age and calculated statistics for each perturbed dataset. To ensure that our results are robust, for each strategy we test, we performed the said perturbations for N=100 iterations on the input data. This then allows us to report distributions of statistics for each level of treatment within each perturbation strategy.

### Simulating data sparsity

To simulate data sparsity, we have devised three data dropout strategies to replace a randomly selected subset of data and replace them with NaNs/missing data. For the first approach, we randomly selected M% of the data points across the entire dataset, without ensuring that an equal portion of data is dropped from each sample or each CpG locus. For the second approach, we stratified the data such that within each sample, M% of different selections of CpG probes are removed. For the third approach, we selected M% of the CpG probes and ensured that the same probes would be dropped out across all samples. The value of M is selected from [0.05, 0.1, 0.2, 0.3, 0.4, 0.5, 0.6, 0.7, 0.8, 0.9, 0.95]. The first dropout strategy is used for the majority of the manuscript unless otherwise specified.

### Simulating bisulfite conversion error

It is well known that errors caused by harsh bisulfite treatment conditions could skew the number of reads captured by bisulfite sequencing-based technologies (6,23). To model bisulfite conversion errors, we have added Gaussian noise with various levels of standard deviation S, where the value of S is selected from [0.01, 0.02, 0.05, 0.1, 0.2, 0.5, 1]. To give an example: we have simulated low read-depth for cg000001, which has a methylation level of 0.6 based on the original Illumina array data. We are interested in simulating a read depth of 4 for this CpG probe. We will then perform the simulation. If we get a result of 2 reads, the transformed p would have a value of 0.5. If we get a result of 3 reads, the transformed p would have a value of 0.75. 0 and standard deviations SD in {0.01, 0.02, 0.05, 0.1, 0.2, 0.5, 1}, and then trimmed the resulting values to range of [0, 1].

### Simulating low read depth

To simulate low read depth, we have treated each data point as a prior of a binomial distribution, where the posterior will be the result of

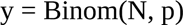

where p is the previously defined prior and n is the read depth. The value of N is selected from [1, 2, 3, 4, 5, 6, 7, 8, 9, 10, 20, 30, 40, 50]. This transformation results in a y value which is correlated with the original p, albeit with a much larger noise due to the transformation.

To give an example: we have simulated low read depth for cg000001, which has a methylation level of 0.6 based on the original Illumina array data. We are interested in simulating a read depth of 4 for this CpG probe. We will then perform the simulation. If we get a result of 2 reads, the transformed p would have a value of 0.5. If we get a result of 3 reads, the transformed p would have a value of 0.75.

We have simulated two types of read depth limitations. For the first approach, we have assumed all probes have a uniform low read depth D, where D ranges from 1 to an arbitrary large number (D=50). For the second and more realistic approach, we inspected a dataset generated by scBS-seq (35) and curated a set of read-depth distributions that resemble real scBS-seq data; we then randomly assigned each probe to have a particular read depth such that the read depth distribution of our synthetic data matches that of real data with low read depth. Our detailed read depth distribution for the second approach is included in graphical and tabular form in Table S1.

### Combining data perturbation strategies

It is unclear how the different data perturbation strategies would interfere with each other, so we chose to perform a grid-combination of data perturbations of the above three types of perturbations. We chose to combine 1) random data dropout, 2) bisulfite conversion error, and 3) real data read depth-based binarization. We chose to combine several settings and performed a grid search. Our approach could be summarized by the following formula:

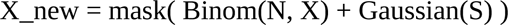

Where X and X_new represent our input data before and after data perturbation. N can either be a constant value n, or a matrix N of read-depths matching the rows/columns of X_array, such that each locus on each pseudo-bulk sample has an associated read-depth value. G is a matrix of random noise values matching the rows/columns of X_array. The mask() function randomly drops a percentage of data. Binom() describes the binomial sampling procedure, where we 1) randomly selects to add a read or not based on the input variable, and then 2) repeat for N times to obtain several reads that would be our final output.

### Testing data-centric quality control

To conduct quality control by removing the lowest quality samples, we first obtained samples from scTrio-seq data as mentioned above (35), and calculated data dropout and read depth constraints from the data. With m samples of interest and m sets of data quality standards, we randomly select m samples from our input data without replacement, and then apply data quality standards to the selected samples. This process is repeated for N=100 iterations.

Then, we compare accuracy, correlation, and precision against chronological age as compared between the data quality standards, which allows us to rank the m samples/quality standards based on which has the lowest correlation or precision values. Finally, we remove the lowest quality samples in a step-wise fashion, thus resulting in m-i samples for each iteration, and measure accuracy, correlation, and precision against chronological age, until (m-i) <= 3, at the point which data prediction results lose meaning. We then perform statistical analyses as mentioned in the above sections.

### Selection of imputation strategies

Several first-principle imputation strategies have been employed by prior publications. Firstly, it has been suggested by prior research that the most important contributor to the methylation level of a CpG site is a combination of the methylation levels of its nearest neighbors, followed by the distance to those neighbors (24,34). Hence, a natural approach would be to build local regression models of varying complexity using values from a missing value’s neighboring loci. However, this approach requires either the missing values’ neighboring values to be known, or having to span arbitrary long distances across the genome to build “local” models that may not reflect local methylation landscapes. We have proposed several implementations of this approach: 1) take the average values of the two nearest non-NaN values from the same sample or 2) from the same probe, or building a polynomial regression with degree = 3-5) 2, 3, or 4 using a certain number of nearest neighboring loci.

In addition, it has been suggested that most CpG loci tend to have consistent methylation levels, and each sample tends to have relatively consistent overall methylation levels (33). Previous publications have used imputation methods following this principle, including averaging methylation levels across all samples at a specific locus, averaging methylation levels across the entire sample, and clustering similar samples using K-nearest-neighbors (KNN) and then finding the averages within each cluster (4,14).

Alternatively, a simple strategy is to impute with a constant value considering that most methylation sites in the human genome are stably methylated (33). However, this method completely ignores the underlying biology and does not produce variation between samples, thus making between-sample comparisons completely irrelevant at the given CpG site. In the context of linear models, it is worth noting that imputing with the value 0 has the added benefit of “dropping” a variable from the model.

To overcome some shortcomings of first-principle approaches, several imputation papers have suggested creating clusters of samples and then using neighborhood data from the entire cluster to predict local missing CpG values. This strategy mitigates the effects of low coverage while simultaneously allowing neighbor-based imputation and have resulted in several high-performing ML-based imputation methods (25–28). In the spirit of these approaches, we have built several implementations that cluster samples in different fashions, and then conducted 1) probe mean imputation, or 2) built polynomial functions using neighborhood data. We have also directly used the CaMelia method in our project.

### Testing data imputation strategies

To test the efficacy of different imputation methods, we have first devised three sets of data perturbation strategies of differing perturbation severity:

1. Light perturbation (10% random dropout, binarization level 3, 0.01 Gaussian noise).
2. Medium perturbation (25% random dropout, binarization level 5, 0.02 Gaussian noise).
3. Heavy perturbation (50% random dropout, binarization level 6, 0.05 Gaussian noise).

We generated N random iterations (N=100) for each perturbation type. We then imputed the data with various strategies outlined below and compared the results.

### Constant value imputation

For our constant value imputation strategy, we have imputed all NaN values with one of the following values: {0, 0.5, 0.7, 1}. 0 reflects that the probe is unmethylated, 1 reflects that the probe is fully methylated across all cells, 0.5 reflects that the probe is methylated in half of the sample, while 0.7 is a rule-of-thumb number selected as roughly 70-80% of all CpG probes in the human genome are permanently methylated (33). It is also worth noting that imputing with a value of 0 has the unique property of “removing” the predictor from the model and is the de-facto way of handling missing data for our programming language of choice.

Note that some of the imputation strategies outlined below will not completely impute all CpG probes. In those cases, unless otherwise specified, the remaining missing values will be imputed with the constant value of 0.

### Mean value imputation

We have two imputation strategies based on means, “sample mean” and “probe mean”. For each instance of NaN, we will replace it with the mean of the methylation levels across that specific sample (“sample mean”), or the mean of the CpG probe across all other samples (“probe mean”), ignoring any cases of NaNs that may occur.

### Linear imputation using nearest neighboring CpGs

We have two imputation strategies based on averaging the methylation levels of the two nearest neighbors of the NaN value. For each instance of NaN, we will find the two nearest non-NaN CpG probes (“linear probe”) or samples (“linear sample”) and average their methylation levels to obtain methylation levels at the current location.

Note that since the samples are not ordered in any way related to CpG methylation, “linear sample” is expected to perform poorly as it fills the missing value with what are essentially CpG methylation levels of two randomly-selected samples.

### Local polynomial fit imputation

We have a set of imputation strategies that rely on identifying the nearest n non-NaN methylation probes upstream and downstream to a particular instance of NaN, and then building a polynomial regression with an arbitrary degree using these data as inputs (“polyfit_n” or “polyfit_n_distance”). It is possible to either construct these regressions using real relative genomic distances between CpG loci, or by assuming a uniform distance between each CpG probe. In the case that it is impossible to identify n nearest neighbors (start/end of a chromosome), we will either fill the neighbors with a constant value (default = 0.5) or remove them from the polynomial equation. We will then predict the local methylation level from the regression line.

### Clustering mean value imputation

We have a set of imputation strategies that rely on conducting a preliminary imputation on the dataset, clustering each sample based on their overall CpG levels, and then performing “probe mean” imputation as mentioned above within each cluster of samples. For the preliminary imputation strategy, we have selected “linear probe” as it is the best-performing simple imputation strategy. Should a particular probe to still be NaN after imputation, we will impute the remaining probes with “linear probe” imputation, followed by imputing with 0.

For this approach, we have used the following methods: K-nearest neighbors (KNN) clustering, KMeans clustering, Spectral clustering, and Birch clustering. For KNN, we also performed CpG probes clustering.

### Creating synthetic single-cell data from bulk methylation data

To create a synthetic single-cell dataset using bulk data, we have followed a similar approach as the aforementioned low read-depth simulations, where we treated each data point as a prior of a binomial distribution. Unlike the previous method, instead of reporting a single value as the summed number of reads, we will instead report Boolean arrays with lengths that correspond to the read depth level we select. We have first assigned a total number of single-cells M for each sample (we chose M=25 for the manuscript); this value will be the “background” read-depth for CpG probes with no read-depth constraints. Then, we performed bisulfite conversion error modeling by adding Gaussian noise to the bulk data as previously described, and performed data sparsity/low read-depth constraints as previously described after adding Gaussian noise. As a result of this, we have mapped M single-cells for each input sample, where each single cell has a read depth of 0 (no data) or 1 for each CpG loci. We can then perform single-cell bisulfite sequencing-based imputation on this synthetic dataset, and/or sum up reads from single cells to recreate the pseudo-bulk samples as seen in most other parts of our manuscript.

To give an example: we simulate low read depths for cg000001 and cg0000002, which have methylation levels of 0.6 and 0.2 based on the original Illumina array data. We are interested in simulating a read depth of 4 and 3. We will then perform the simulation and may get an output formatted as: [[0, 1, 1, 0], [1, 0, NaN, 0]]. To interpret the results: cell 1 has reads in cg000002 but not cg000001, cell 2 has reads in cg000001 but not cg000002, cell 3 has a read in cg000001 and doesn’t have information for cg000002, whereas cell 4 does not have reads for either cg000001 or cg000002. We can then create a pseudo-bulk data by summing across the single-cells, and the result would be [0.5, 0.33], indicating that we have 50% methylation in cg000001 and 33.3% methylation in cg000002. If we impute the data, we may get a result formatted as [[0, 1, 1, 0], [1, 0, 0, 0]], which can be summed to [0.5, 0.25]; this would indicate a 50% methylation in cg000001 and 25% methylation in cg000002.

### Using CaMelia for single-cell imputation

We have used the CaMelia package as described by the authors (27). Note that we have removed several lines of code that have depreciated in Python 3; we did not modify any significant lines of code. We also used several data processing scripts in tandem with the CaMelia package mainly for data re-formatting, which are provided on our git repository.

The CaMelia package returns an “imputation” dataset which can be merged with the original data to perform the desired imputation function. We then compared age prediction results of our synthetic data with or without CaMelia imputation, accompanied by several first-principle imputation tools as described above.

### Programming

Most programming tasks in this project are performed with Python 3.10. The remaining portions are performed with 1) shell scripting with bash and 2) R programming.

### Availability of data and materials

Illumina 450K array data were obtained from GEO with accession codes GSE72772, GSE72773, and GSE72775 (32). Reference scBS-seq data and scRRBS-seq data are obtained from GEO with accession codes GSE56879 and GSE65364 respectively (8,9). Human single-cell Trio-seq data were obtained from GEO with accession code GSE97693 (35). Epigenetic clock coefficients were obtained from supplementary materials of their respective publications and reformatted to fit with our analysis pipelines; the reformatted coefficient tables are provided on our project git repository for ease of access. The CaMelia epigenetics imputation package scripts were obtained from https://github.com/JxTang-bioinformatics/CaMelia. All other data, notebooks, scripts, and utilities for this publication are available in a permanent git repository on the Samee Lab git repository at https://codeberg.org/sameelab/epiclock-singlecell-benchmarking.

## Authors’ contributions

Z.L and M.A.H.S. conceptualized the work. Z.L performed the experiments. Z.L. and M.A.H.S. wrote and reviewed the manuscript.

## Supporting information

All supplemental figures

All supplemental tables

## Acknowledgments

We thank C.-A. Kapourani (first author of the Melissa epigenetics imputation method) and J. Tang (first author of the CaMelia epigenetics imputation method) for inspiring our work, kindly sharing their code, and assisting us with various programming-related tasks.

## References

1. Moore, L. D., Le, T. & Fan, G. DNA Methylation and Its Basic Function. Neuropsychopharmacology 38, 23–38 (2013).

2. Kulis, M. & Esteller, M. 2 - DNA Methylation and Cancer. in Advances in Genetics (eds. Herceg, Z. & Ushijima, T.) vol. 70 27–56 (Academic Press, 2010).

3. Greenberg, M. V. C. & Bourc’his, D. The diverse roles of DNA methylation in mammalian development and disease. Nature Reviews Molecular Cell Biology 20, 590–607 (2019).

4. Horvath, S. DNA methylation age of human tissues and cell types. Genome Biology 14, 3156 (2013).

5. Horvath, S. & Raj, K. DNA methylation-based biomarkers and the epigenetic clock theory of ageing. Nature Reviews Genetics 19, 371–384 (2018).

6. Karemaker, I. D. & Vermeulen, M. Single-Cell DNA Methylation Profiling: Technologies and Biological Applications. Trends in Biotechnology 36, 952–965 (2018).

7. Guo, H., et al. Single-cell methylome landscapes of mouse embryonic stem cells and early embryos analyzed using reduced representation bisulfite sequencing. Genome Research 23, 2126–2135 (2013).

8. Smallwood, S. A. et al. Single-cell genome-wide bisulfite sequencing for assessing epigenetic heterogeneity. Nature Methods 11, 817–820 (2014).

9. Hou, Y. et al. Single-cell triple omics sequencing reveals genetic, epigenetic, and transcriptomic heterogeneity in hepatocellular carcinomas. Cell Research 26, 304–319 (2016).

10. Clark, S. J. et al. scNMT-seq enables joint profiling of chromatin accessibility DNA methylation and transcription in single-cells. Nature Communications 9, 781 (2018).

11. Hannum, G., et al. Genome-wide Methylation Profiles Reveal Quantitative Views of Human Aging Rates. Molecular Cell 49 (2), 359–367 (2012).

12. Levine, M. E. et al. An epigenetic biomarker of aging for lifespan and healthspan. Aging (Albany NY) 10, 573–591 (2018).

13. Horvath, S. et al. Epigenetic clock for skin and blood cells applied to Hutchinson Gilford Progeria Syndrome and ex vivo studies. Aging (Albany NY) 10, 1758–1775 (2018).

14. Zhang, Q. et al. Improved precision of epigenetic clock estimates across tissues and its implication for biological ageing. Genome Medicine 11, 54 (2019).

15. Levy, J. J. et al. MethylNet: an automated and modular deep learning approach for DNA methylation analysis. BMC Bioinformatics 21, 108 (2020).

16. Liu, Z. et al. Underlying features of epigenetic aging clocks in vivo and in vitro. Aging Cell 19, e13229 (2020).

17. de Lima Camillo, L. P., Lapierre, L. R. & Singh, R. A pan-tissue DNA-methylation epigenetic clock based on deep learning. npj Aging 8, 4 (2022).

18. Horvath, S. & Levine, A. J. HIV-1 Infection Accelerates Age According to the Epigenetic Clock. J Infect Dis 212, 1563–1573 (2015).

19. Higgins-Chen, A. T., et al. A computational solution for bolstering reliability of epigenetic clocks: implications for clinical trials and longitudinal tracking. Nature Aging 2, 644–661 (2022).

20. Meacham, C. E. & Morrison, S. J. Tumour heterogeneity and cancer cell plasticity. Nature 501, 328–337 (2013).

21. Hernando-Herraez, I. et al. Ageing affects DNA methylation drift and transcriptional cell-to-cell variability in mouse muscle stem cells. Nature Communications 10, 4361 (2019).

22. Trapp, A., Kerepesi, C. & Gladyshev, V. N. Profiling epigenetic age in single-cells. Nature Aging 1, 1189–1201 (2021).

23. Genereux, D. P., Johnson, W. C., Burden, A. F., Stöger, R. & Laird, C. D. Errors in the bisulfite conversion of DNA: modulating inappropriate- and failed-conversion frequencies. Nucleic Acids Res 36, e150 (2008).

24. Angermueller, C., Lee, H. J., Reik, W. & Stegle, O. DeepCpG: accurate prediction of single-cell DNA methylation states using deep learning. Genome Biology 18, 67 (2017).

25. Kapourani, C.-A. & Sanguinetti, G. Melissa: Bayesian clustering and imputation of single-cell methylomes. Genome Biology 20, 61 (2019).

26. P. E. de Souza, C., et al. Epiclomal: Probabilistic clustering of sparse single-cell DNA methylation data. PLOS Computational Biology 16, e1008270 (2020).

27. Tang, J. et al. CaMelia: imputation in single-cell methylomes based on local similarities between cells. Bioinformatics 37, 1814–1820 (2021).

28. De Waele, G., Clauwaert, J., Menschaert, G. & Waegeman, W. CpG Transformer for imputation of single-cell methylomes. Bioinformatics 38, 597–603 (2022).

29. Lin, Q. & Wagner, W. Epigenetic Aging Signatures Are Coherently Modified in Cancer. PLOS Genetics 11, e1005334 (2015).

30. Zheng, C., Li, L. & Xu, R. Association of Epigenetic Clock with Consensus Molecular Subtypes and Overall Survival of Colorectal Cancer. Cancer Epidemiology, Biomarkers & Prevention 28, 1720–1724 (2019).

31. Zheng, C., Berger, N. A., Li, L. & Xu, R. Epigenetic age acceleration and clinical outcomes in gliomas. PLOS ONE 15, e0236045 (2020).

32. Horvath, S. et al. An epigenetic clock analysis of race/ethnicity, sex, and coronary heart disease. Genome Biology 17, 171 (2016).

33. Ziller, M. J. et al. Charting a dynamic DNA methylation landscape of the human genome. Nature 500, 477–481 (2013).

34. Zhang, W., Spector, T. D., Deloukas, P., Bell, J. T. & Engelhardt, B. E. Predicting genome-wide DNA methylation using methylation marks, genomic position, and DNA regulatory elements. Genome Biology 16, 14 (2015).

35. Bian, S. et al. Single-cell multiomics sequencing and analyses of human colorectal cancer. Science 362, 1060–1063 (2018).

36. Robinson, M. D., McCarthy, D. J. & Smyth, G. K. edgeR: a Bioconductor package for differential expression analysis of digital gene expression data. Bioinformatics 26, 139–140 (2010).

37. George, N. I., Bowyer, J. F., Crabtree, N. M. & Chang, C.-W. An Iterative Leave-One-Out Approach to Outlier Detection in RNA-Seq Data. PLOS ONE 10, e0125224 (2015).

38. Assenov, Y. et al. Comprehensive analysis of DNA methylation data with RnBeads. Nat Methods 11, 1138–1140 (2014).

39. Müller, F. et al. RnBeads 2.0: comprehensive analysis of DNA methylation data. Genome Biology 20, 55 (2019).

