## Supplementary material for "Imputation approaches and quality standards for single-cell epigenetic age predictions": All supplemental figures

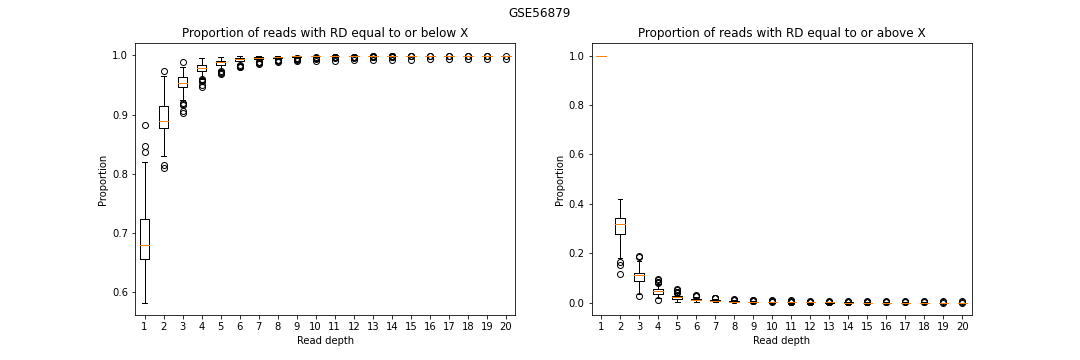

**Figure S1**. Boxplots showing the cumulative proportion of cells with read depth equal to or below (left), or equal to or above (right) a certain count for the dataset GSE56879.

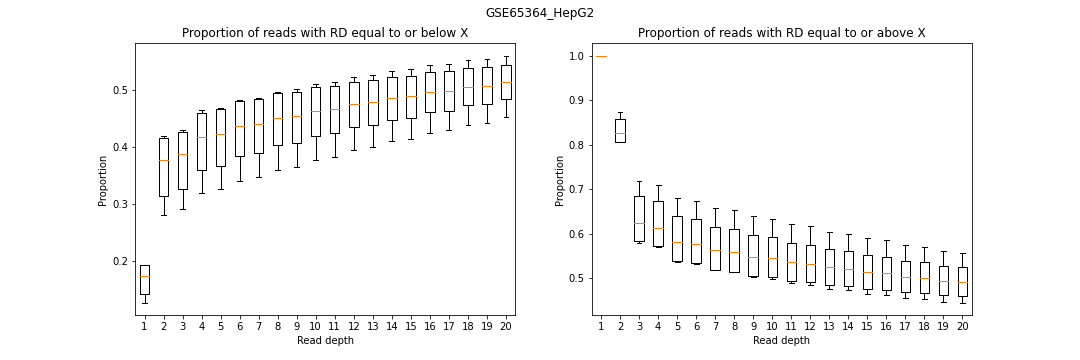

**Figure S2**. Boxplots showing the cumulative proportion of cells with read depth equal to or below (left), or equal to or above (right) a certain count in HepG2 cells for the dataset GSE65364.

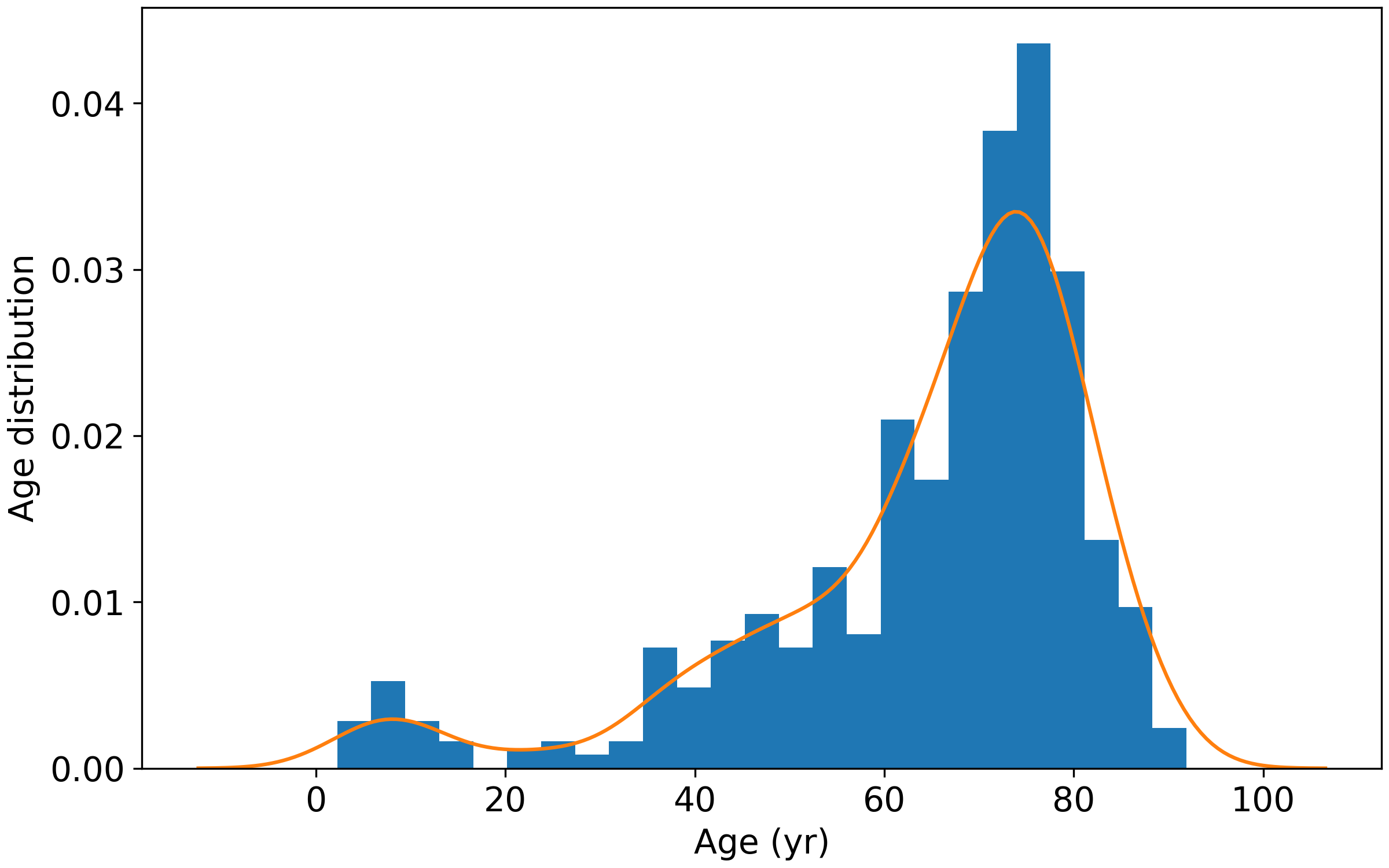

**Figure S3**. Age distribution of samples in input data.

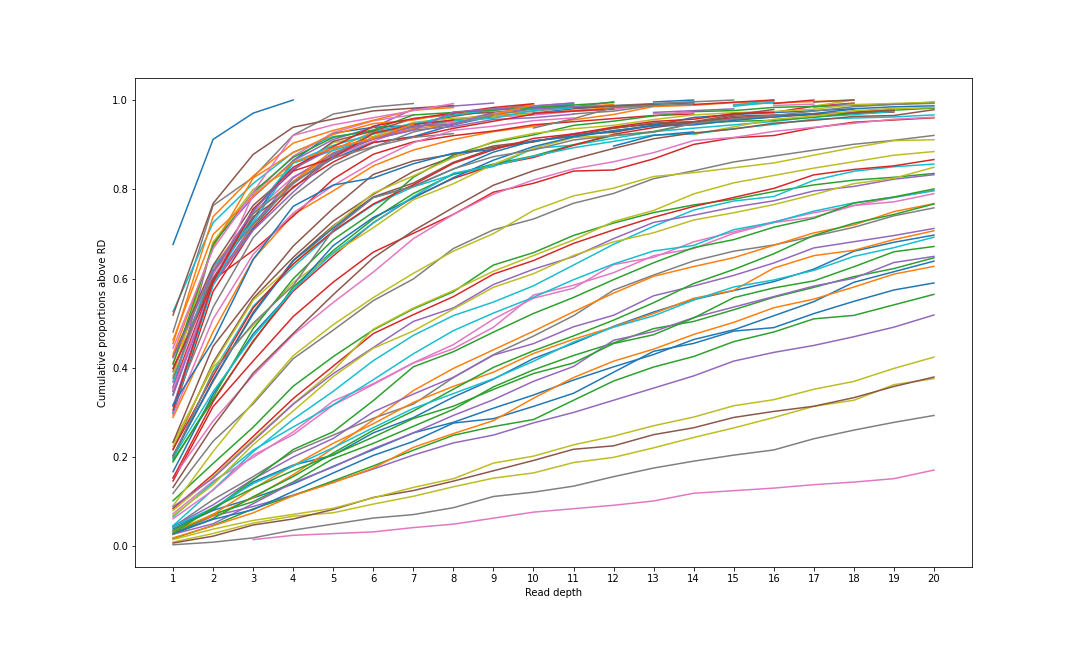

**Figure S4**. Cumulative read depth distributions as observed in scTrio-seq data.

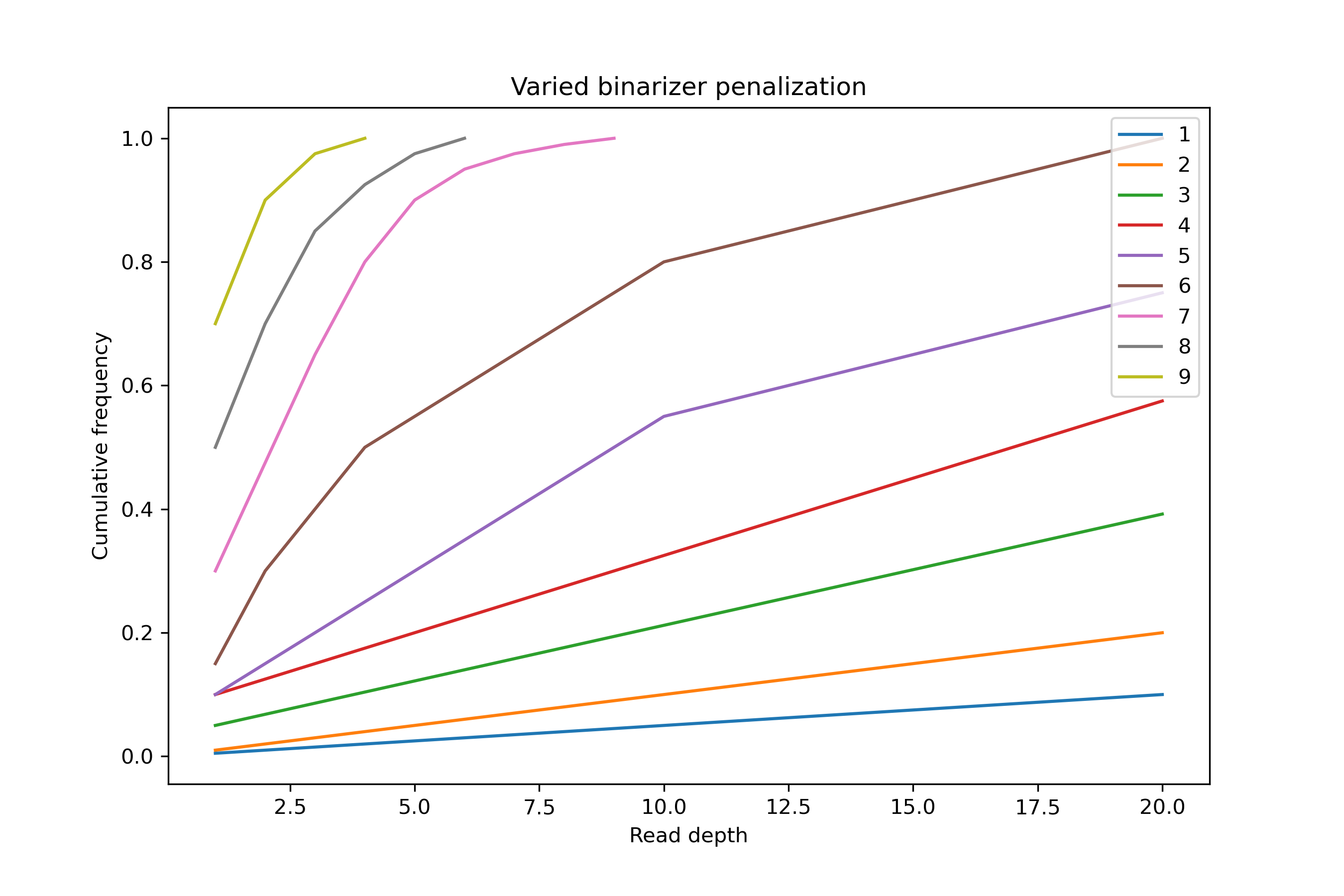

**Figure S5**. Curated read depth distributions based on distributions as observed in scTrio-seq data.

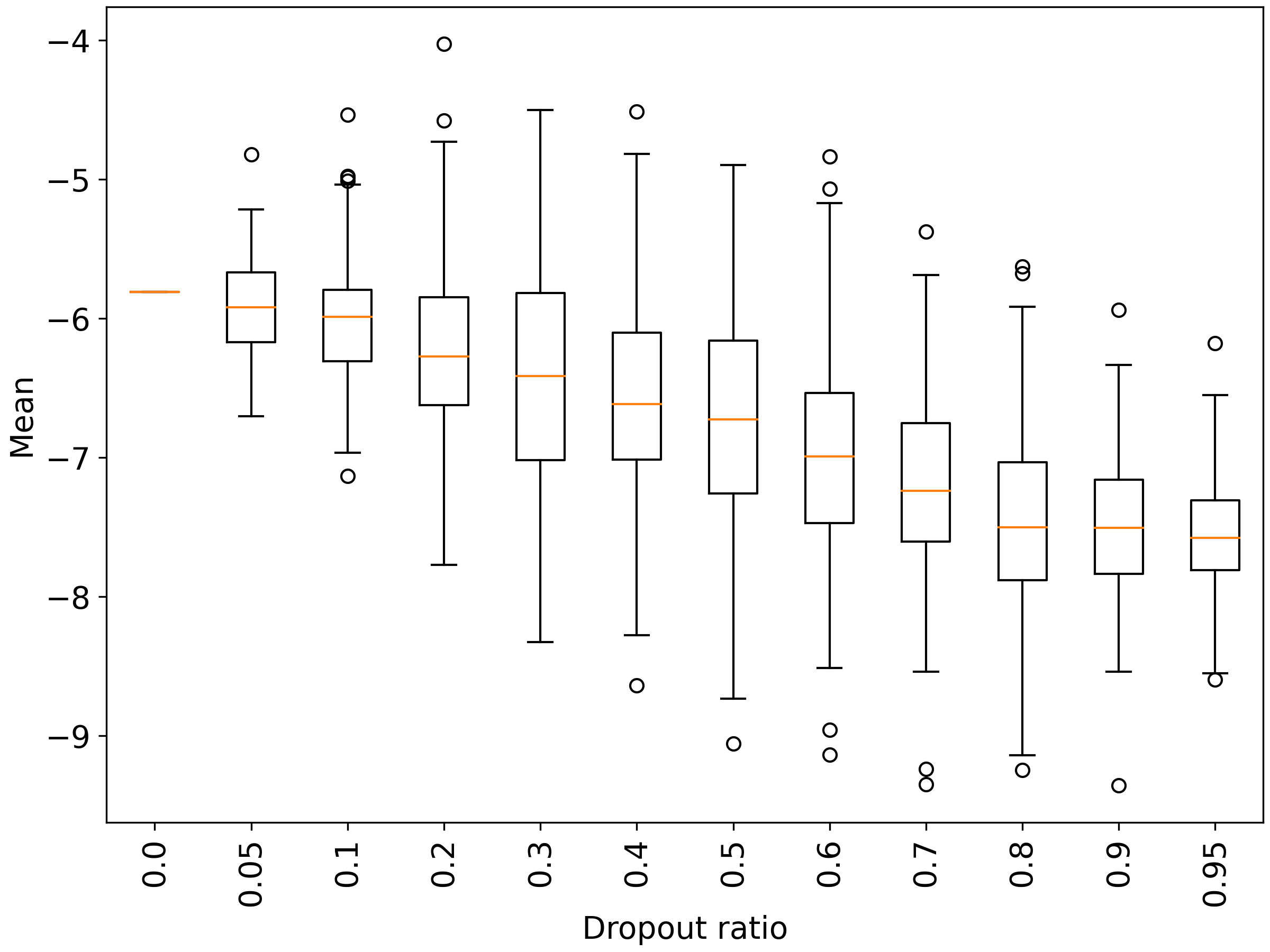
**Figure S6**. Boxplot showing distributions of mean at various degrees of data dropout.

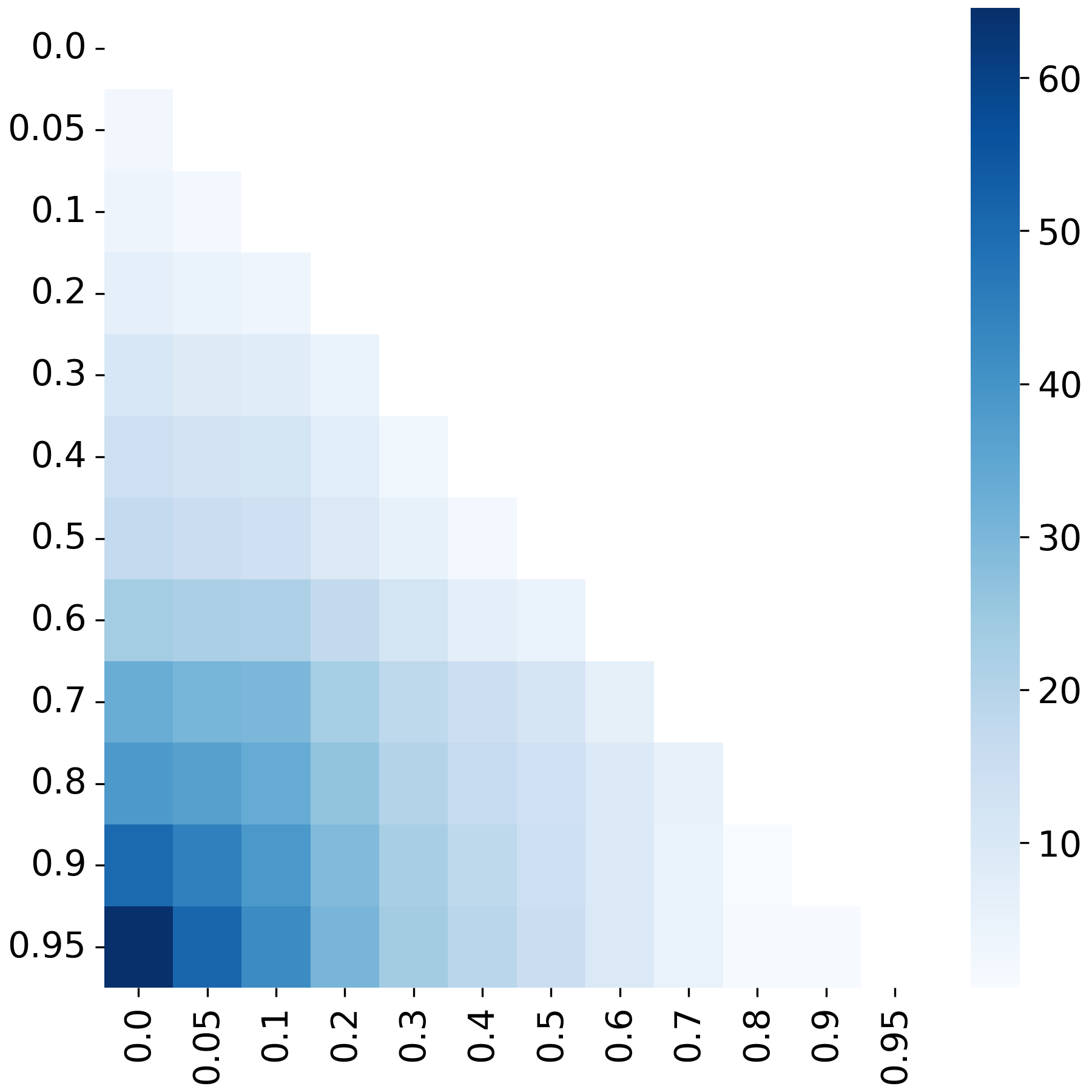

**Figure S7.** Heatmap showing pairwise t-tests of mean at various degrees of data dropout. The colorbar shows -log10-transformed p-values.

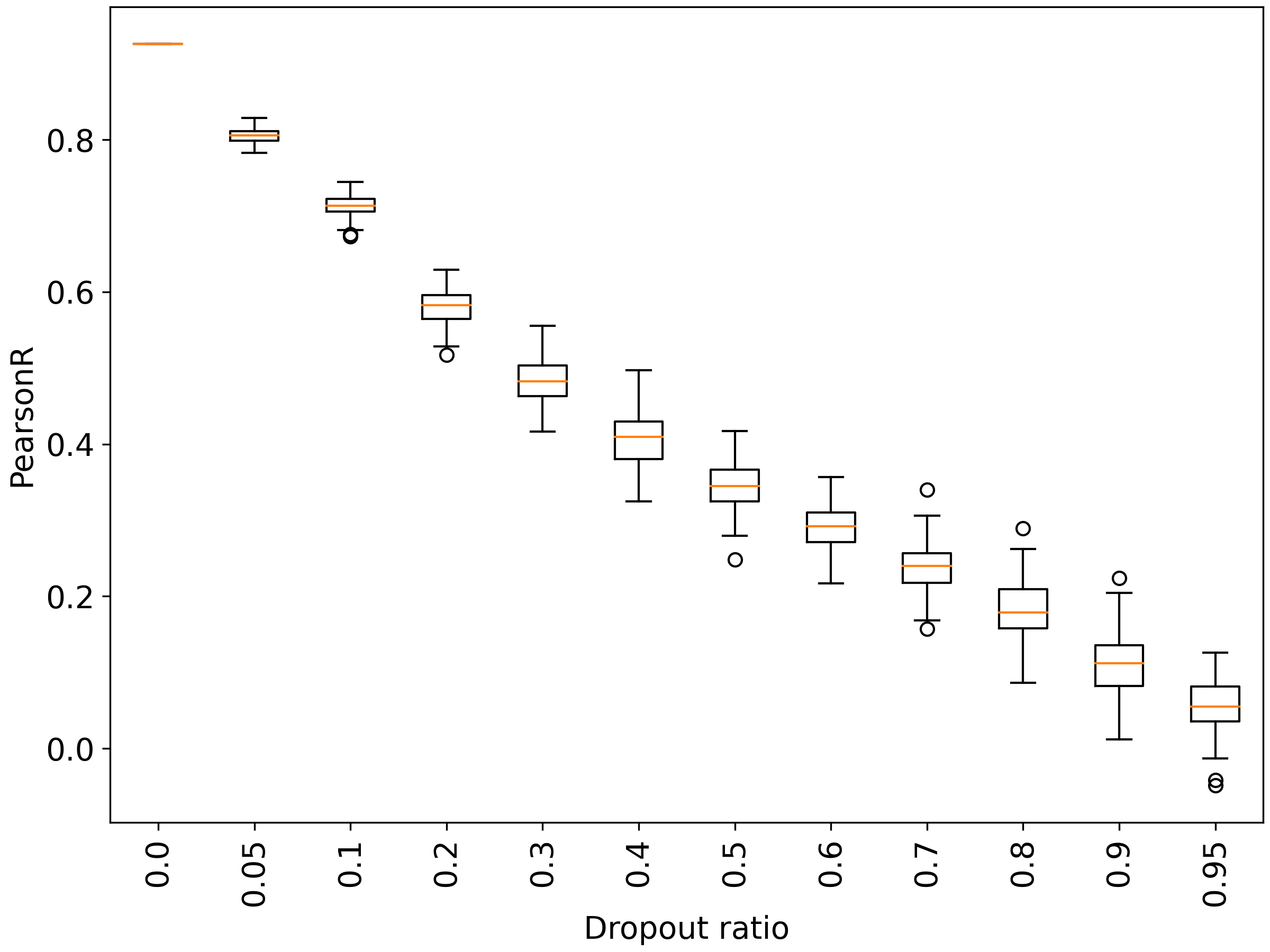

**Figure S8.** Boxplot showing distributions of Pearson correlation at various degrees of data dropout.

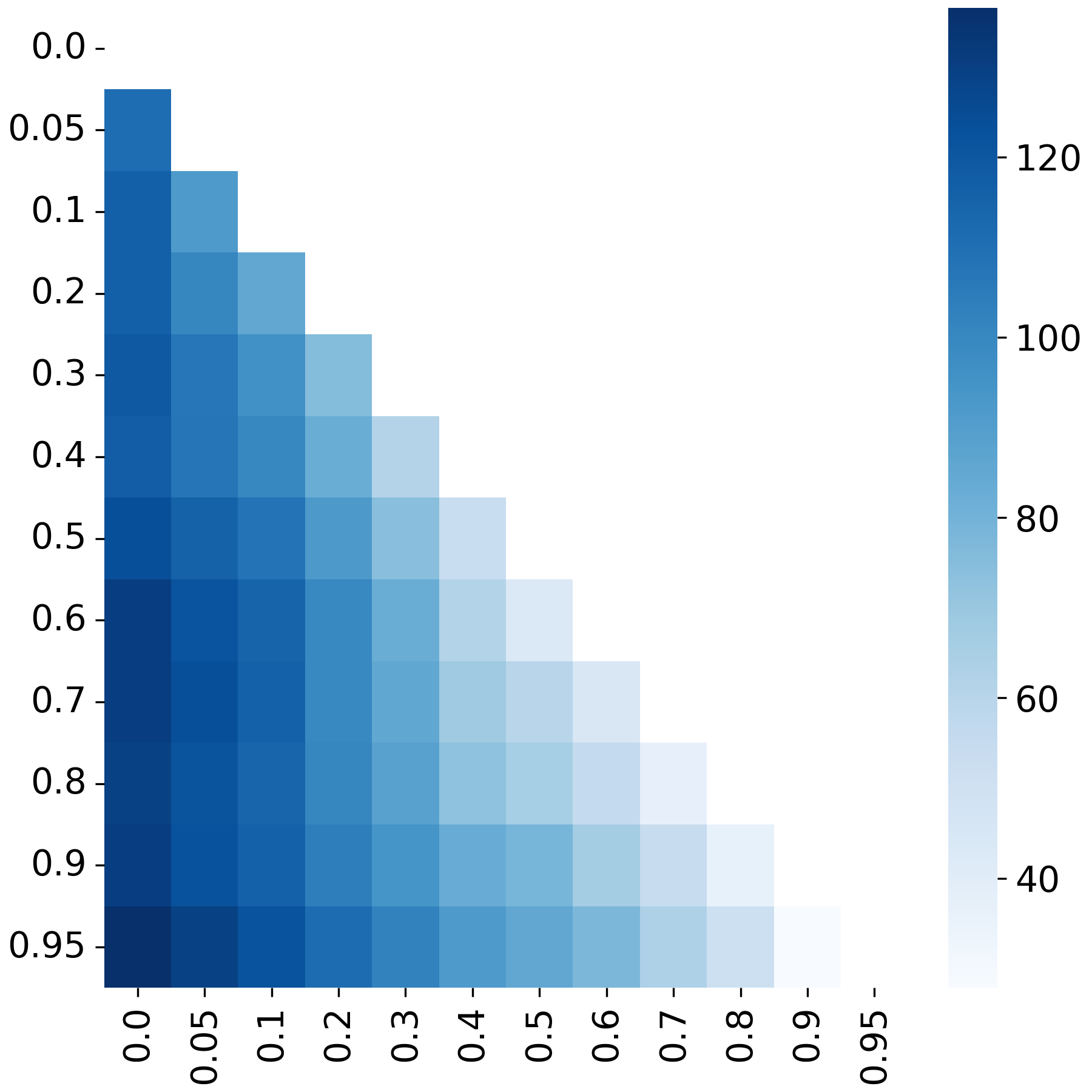

**Figure S9.** Heatmap showing pairwise t-tests of Pearson correlation at various degrees of data dropout. The colorbar shows -log10-transformed p-values.

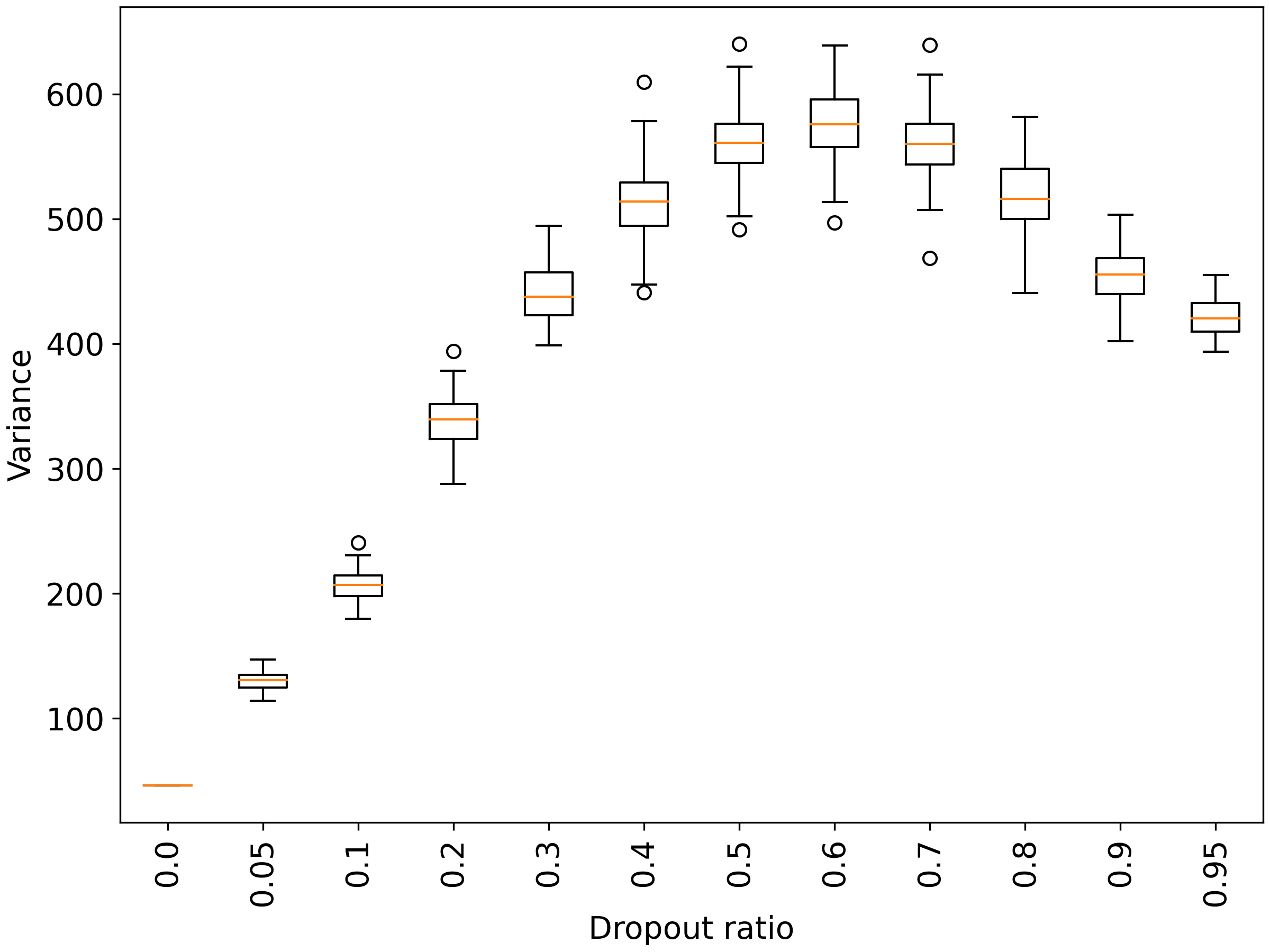
**Figure S10**. Boxplot showing distributions of variance at various degrees of data dropout.

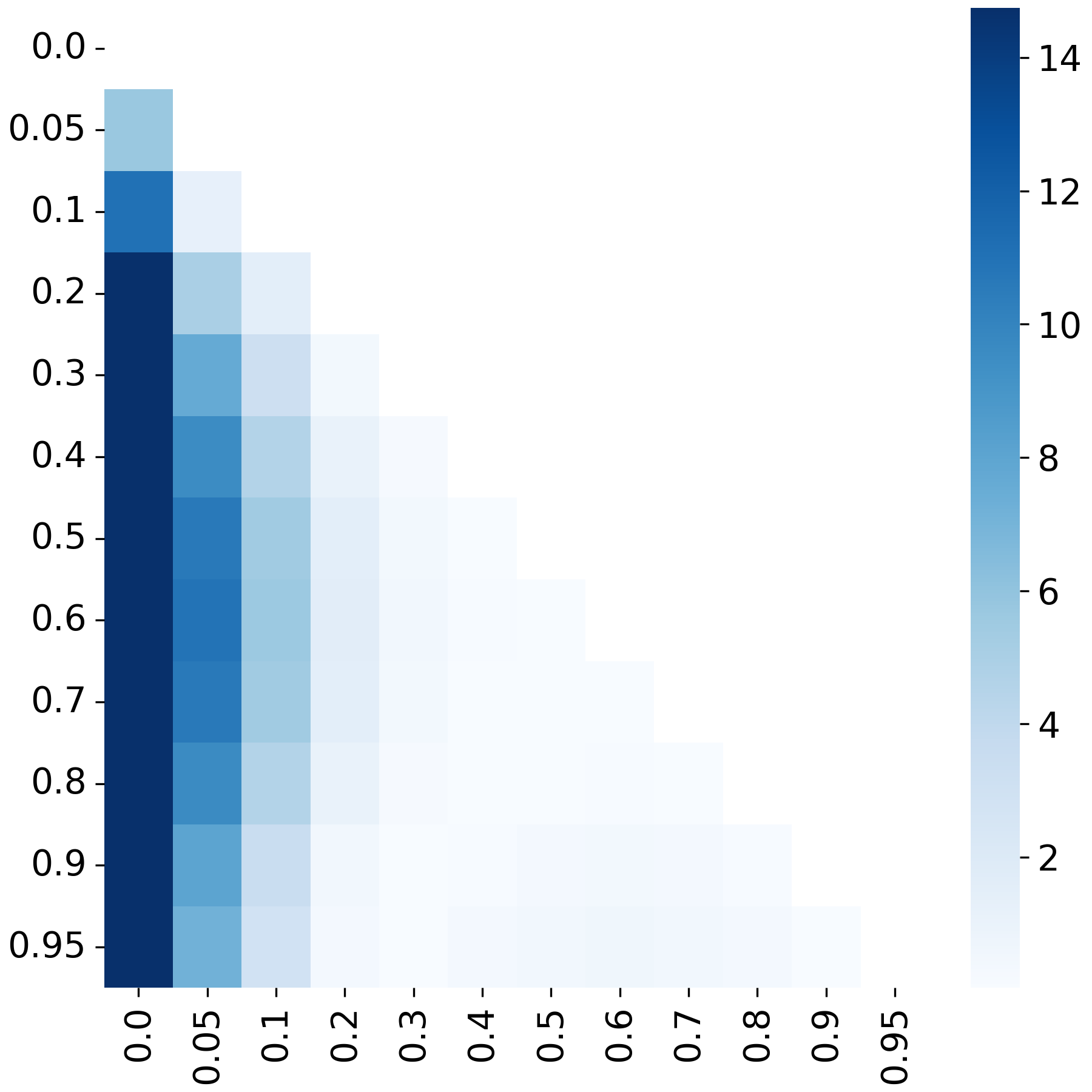

**Figure S11**. Heatmap showing pairwise F-tests of variance at various degrees of data dropout. The colorbar shows -log10-transformed p-values.

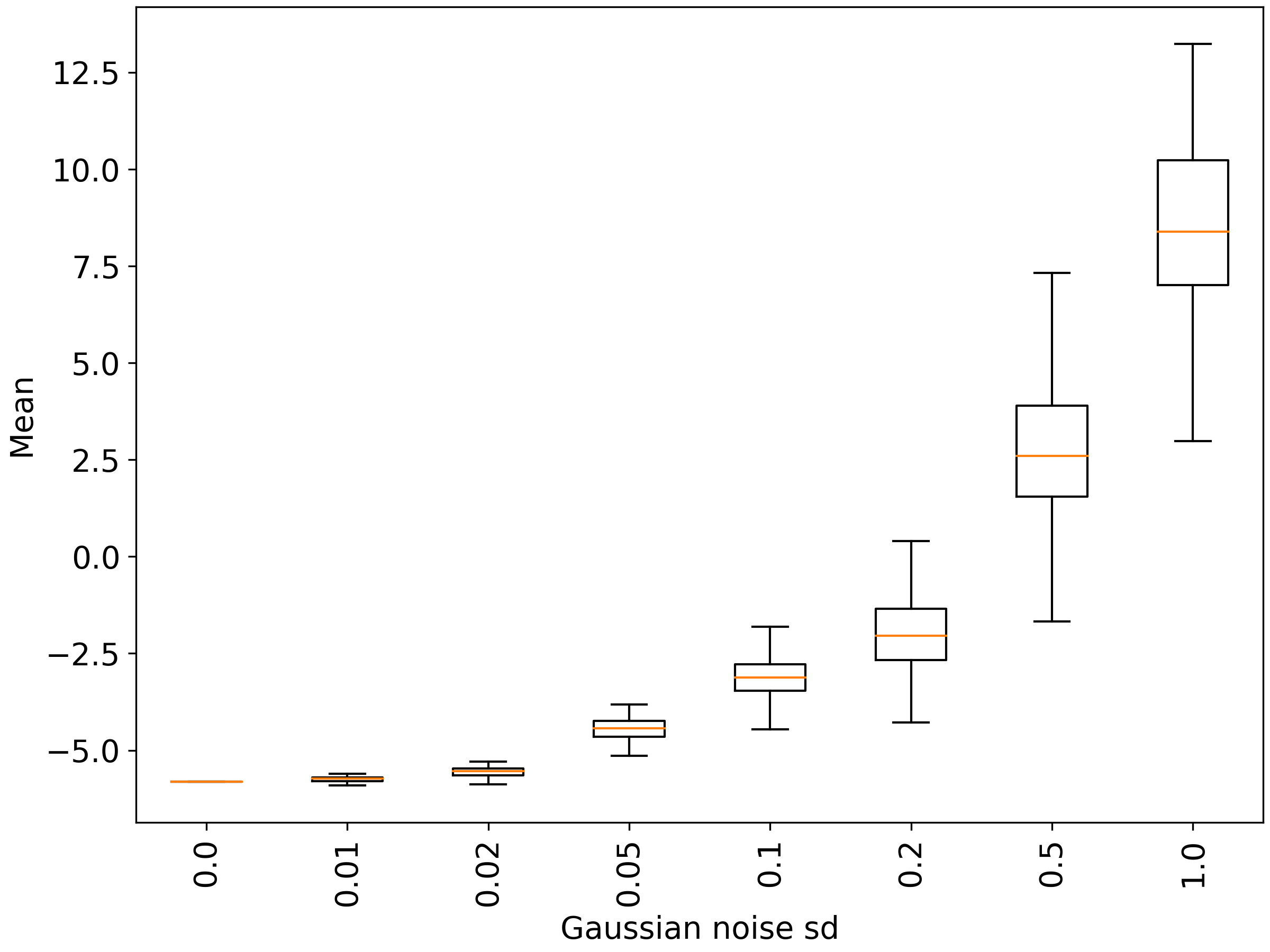

**Figure S12**. Boxplot showing distributions of mean when adding Gaussian noise with various levels of standard deviation.

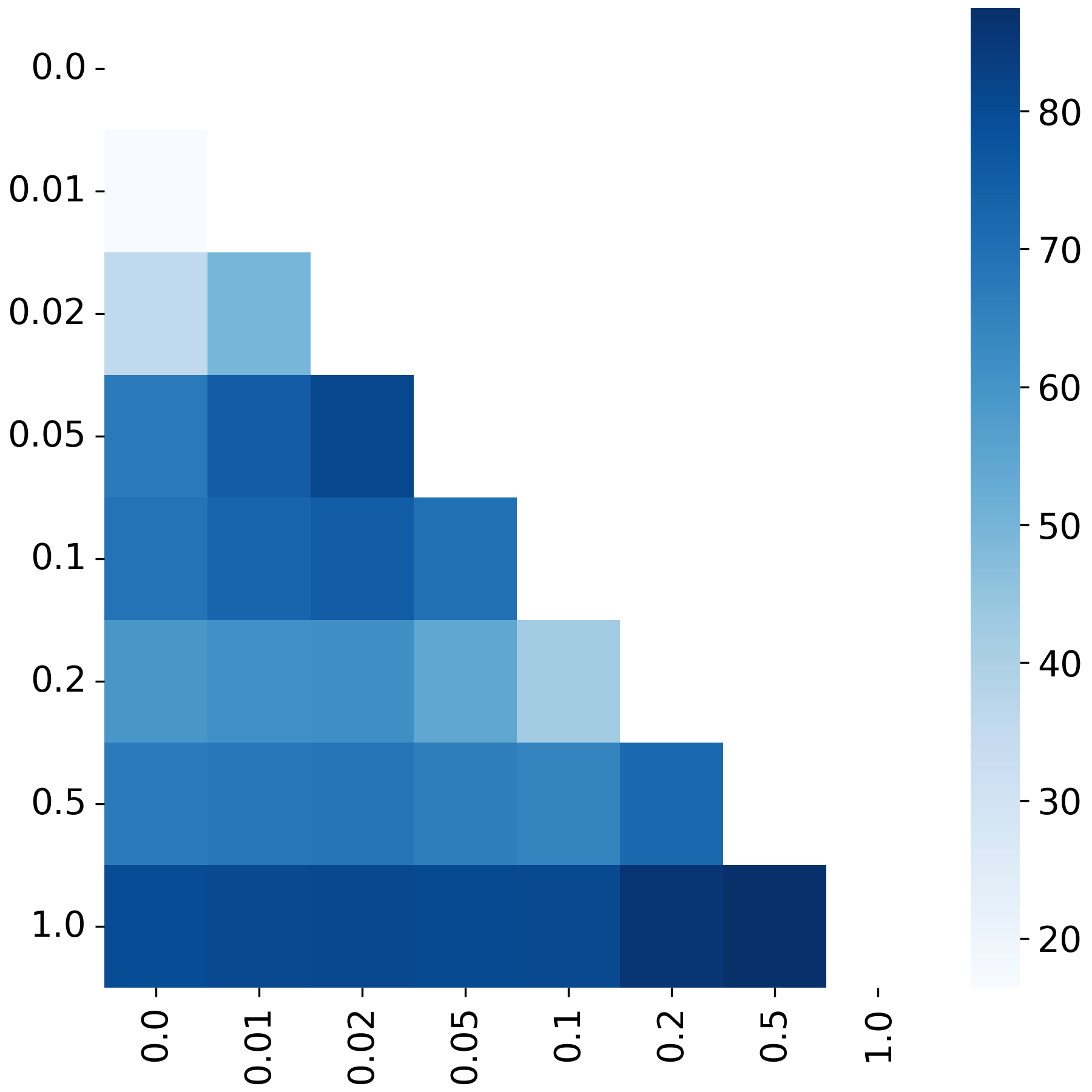

**Figure S13**. Heatmap showing pairwise t-tests of mean when adding Gaussian noise with various levels of standard deviation. The colorbar shows -log10-transformed p-values.

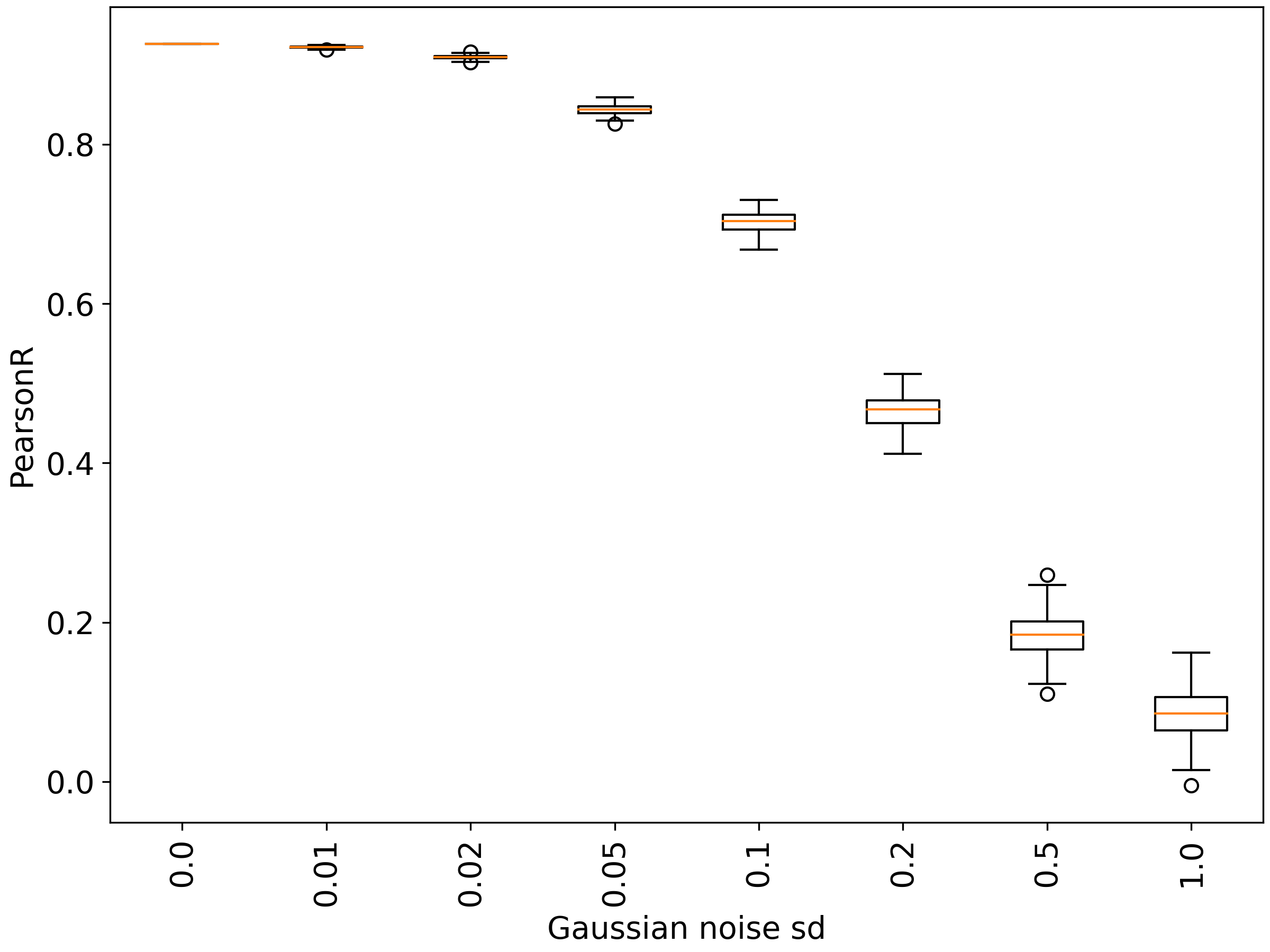

**Figure S14**. Boxplot showing distributions of Pearson correlation when adding Gaussian noise with various levels of standard deviation.

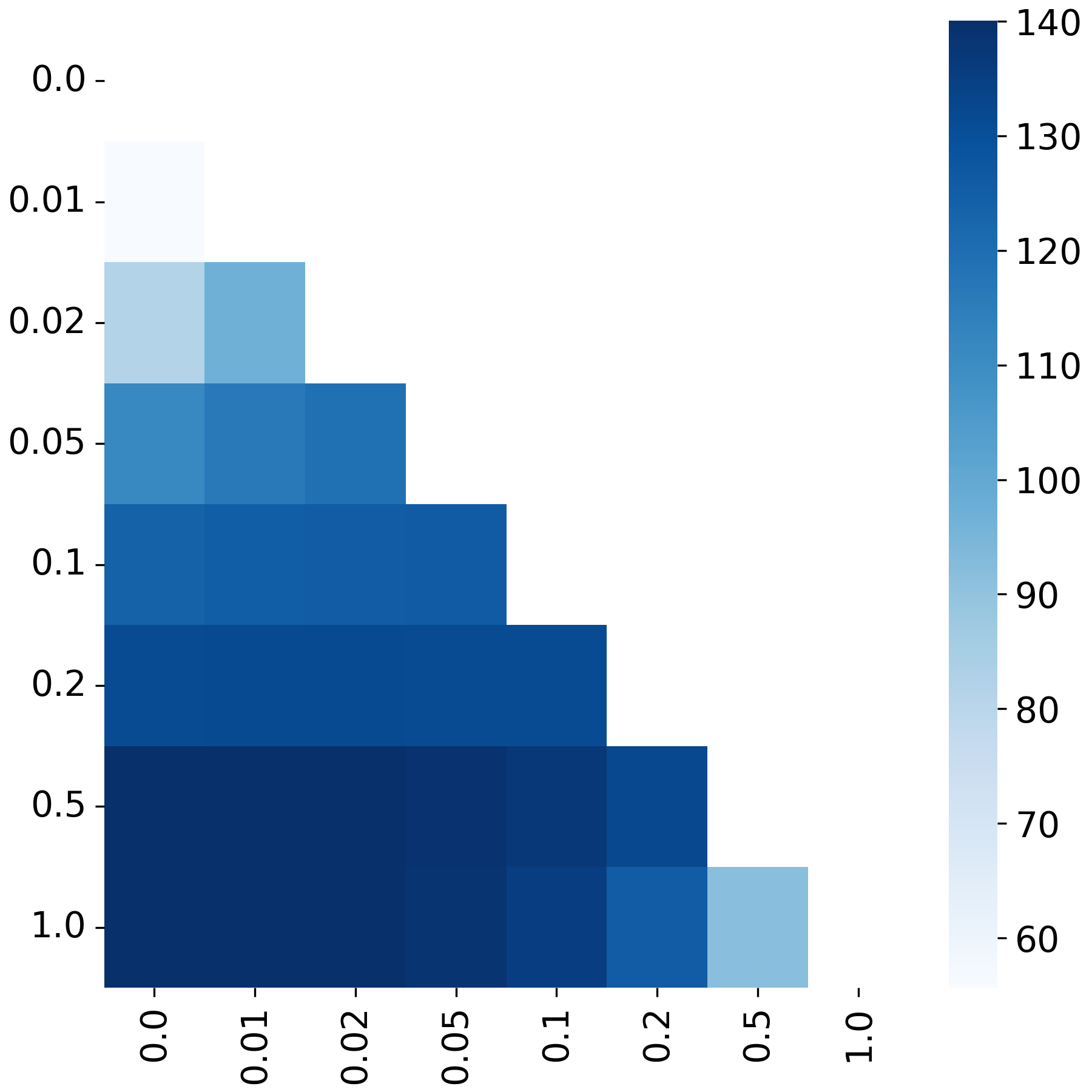

**Figure S15**. Heatmap showing pairwise t-tests of Pearson correlation when adding Gaussian noise with various levels of standard deviation. The colorbar shows -log10-transformed p-values.

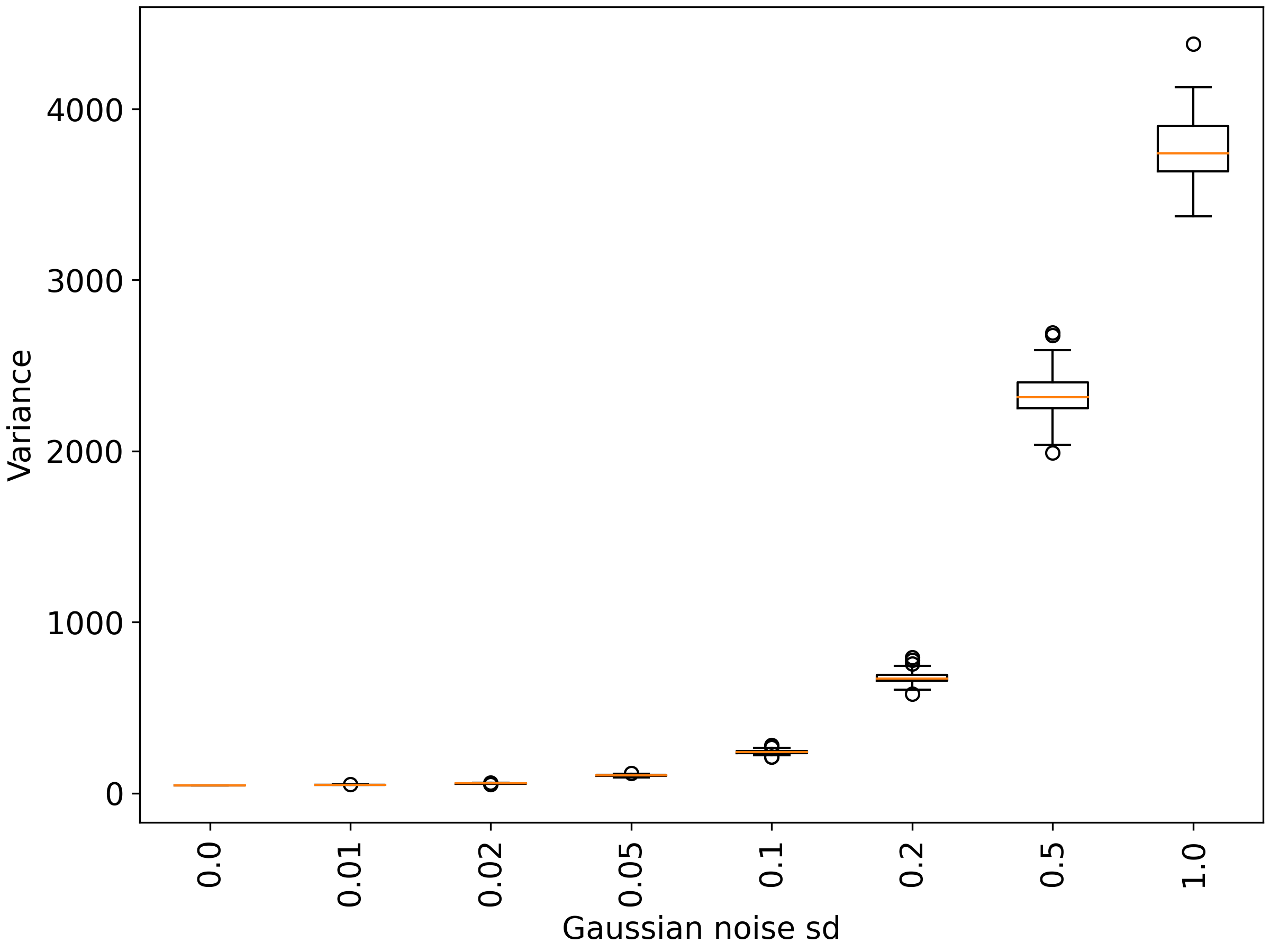

**Figure S16**. Boxplot showing distributions of variance when adding Gaussian noise with various levels of standard deviation.

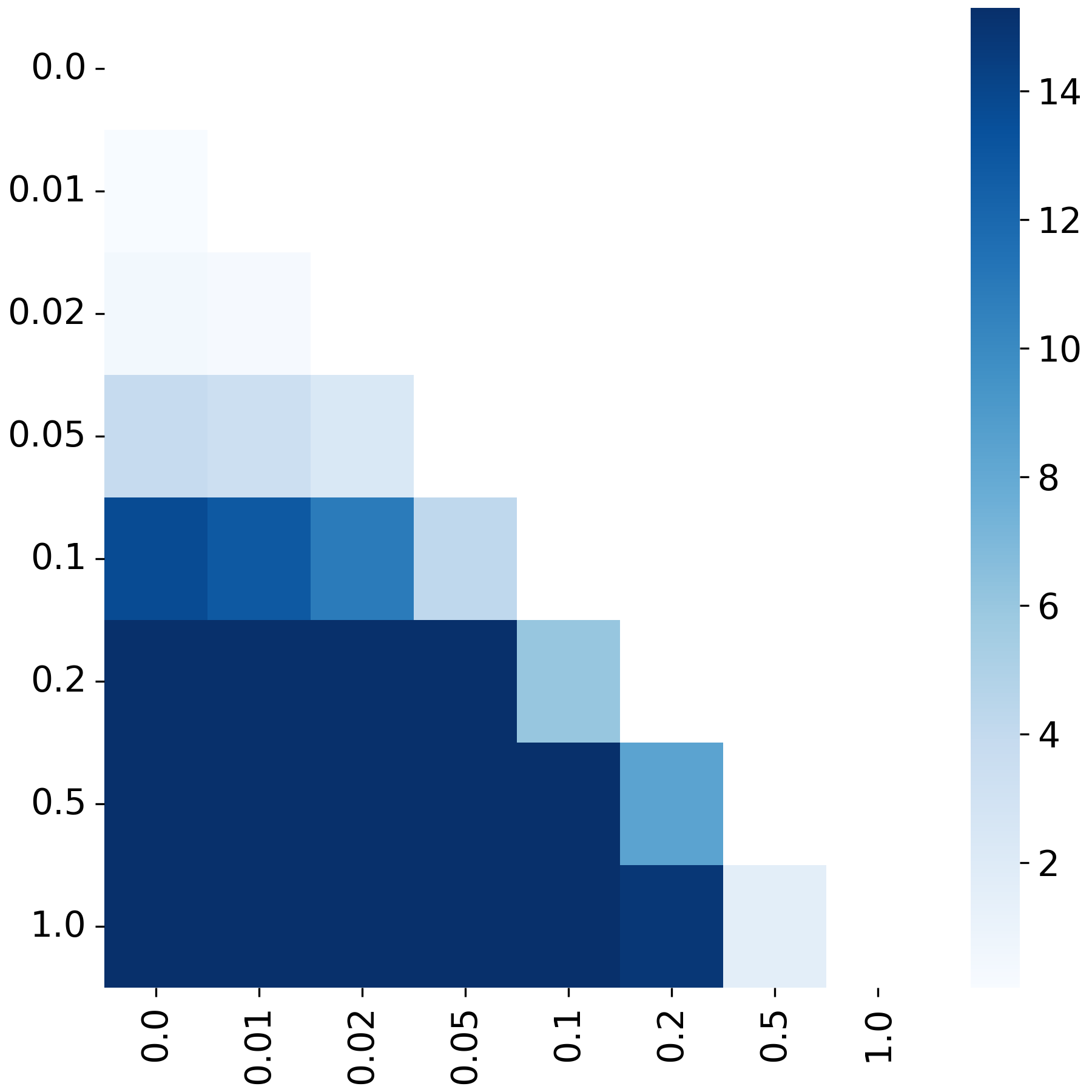

**Figure S17**. Heatmap showing pairwise F-tests of mean when adding Gaussian noise with various levels of standard deviation. The colorbar shows -log10-transformed p-values.

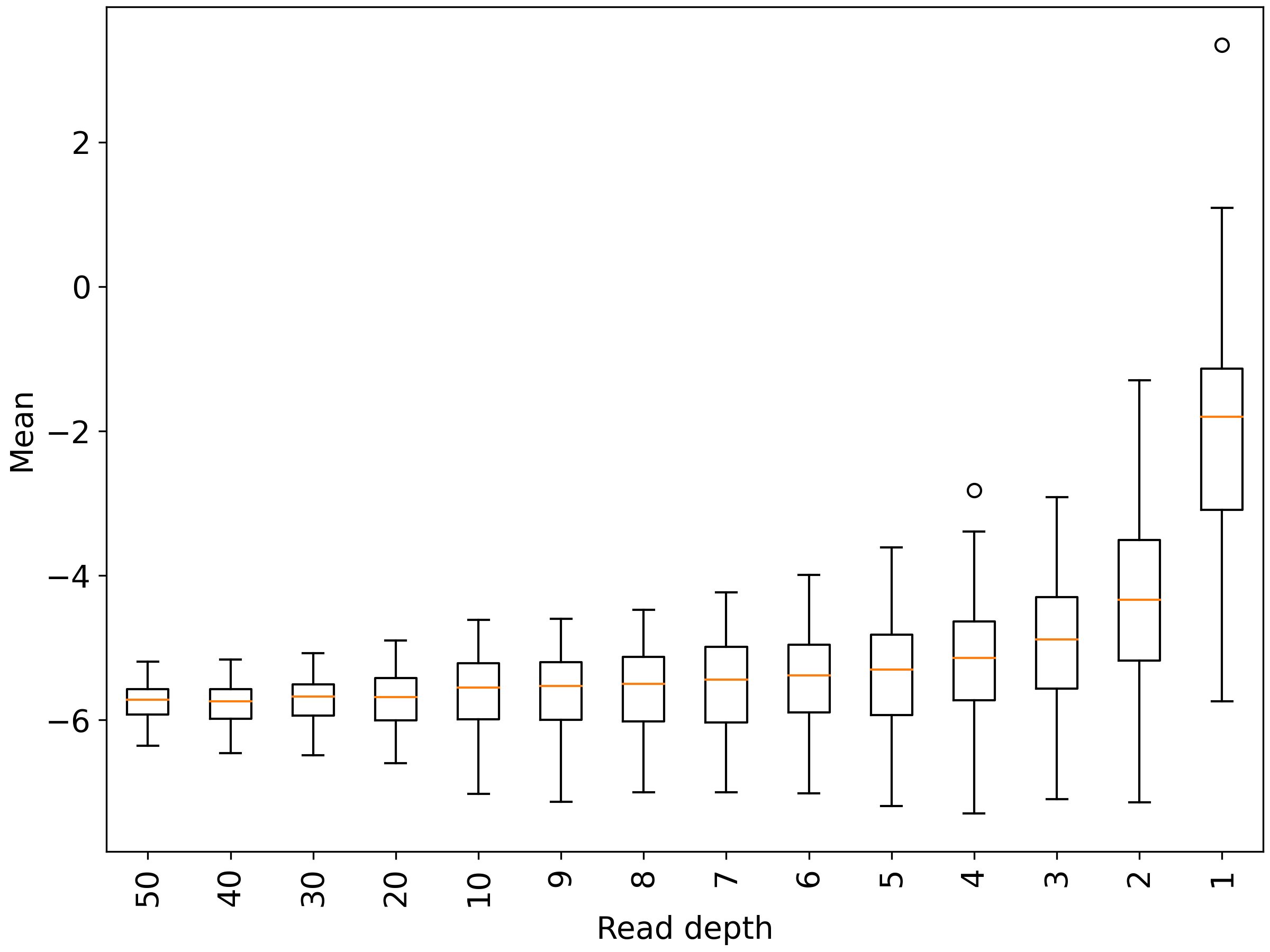

**Figure S18**. Boxplot showing distributions of mean when restricting data to a uniform read-depth level.

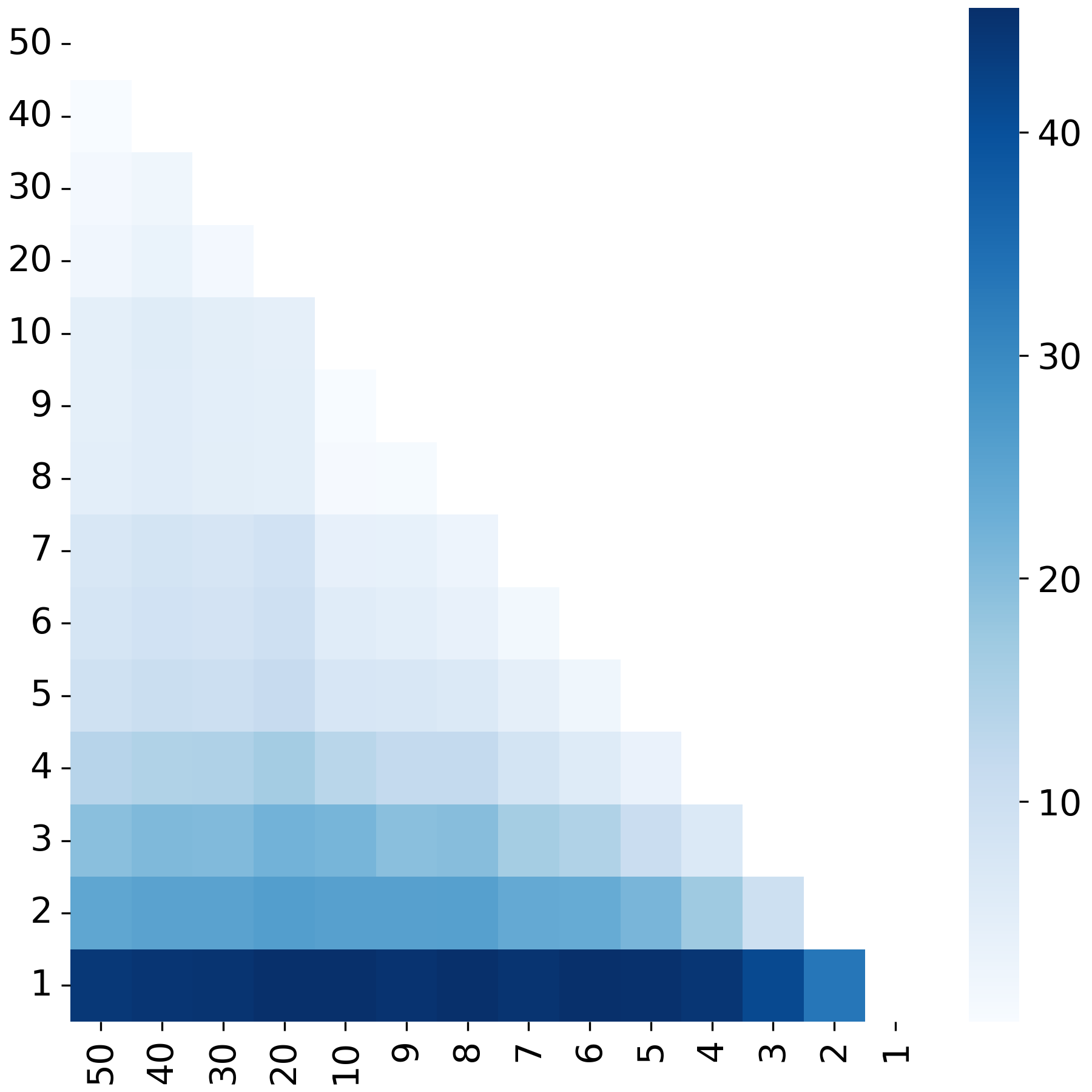

**Figure S19**. Heatmap showing pairwise t-tests of mean when restricting data to a uniform read-depth level. The colorbar shows -log10-transformed p-values.

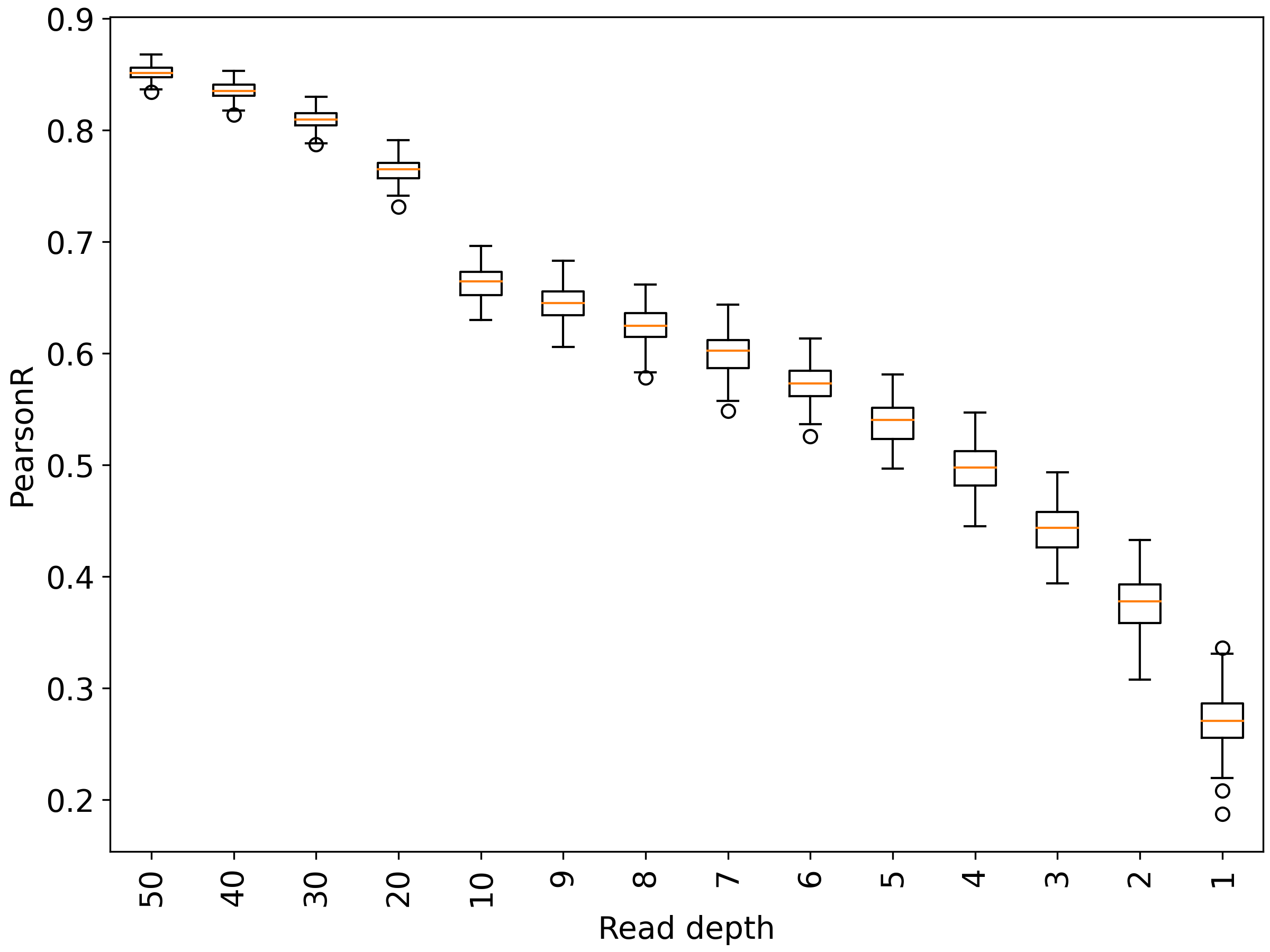

**Figure S20.** Boxplot showing distributions of Pearson correlation when restricting data to a uniform read-depth level.

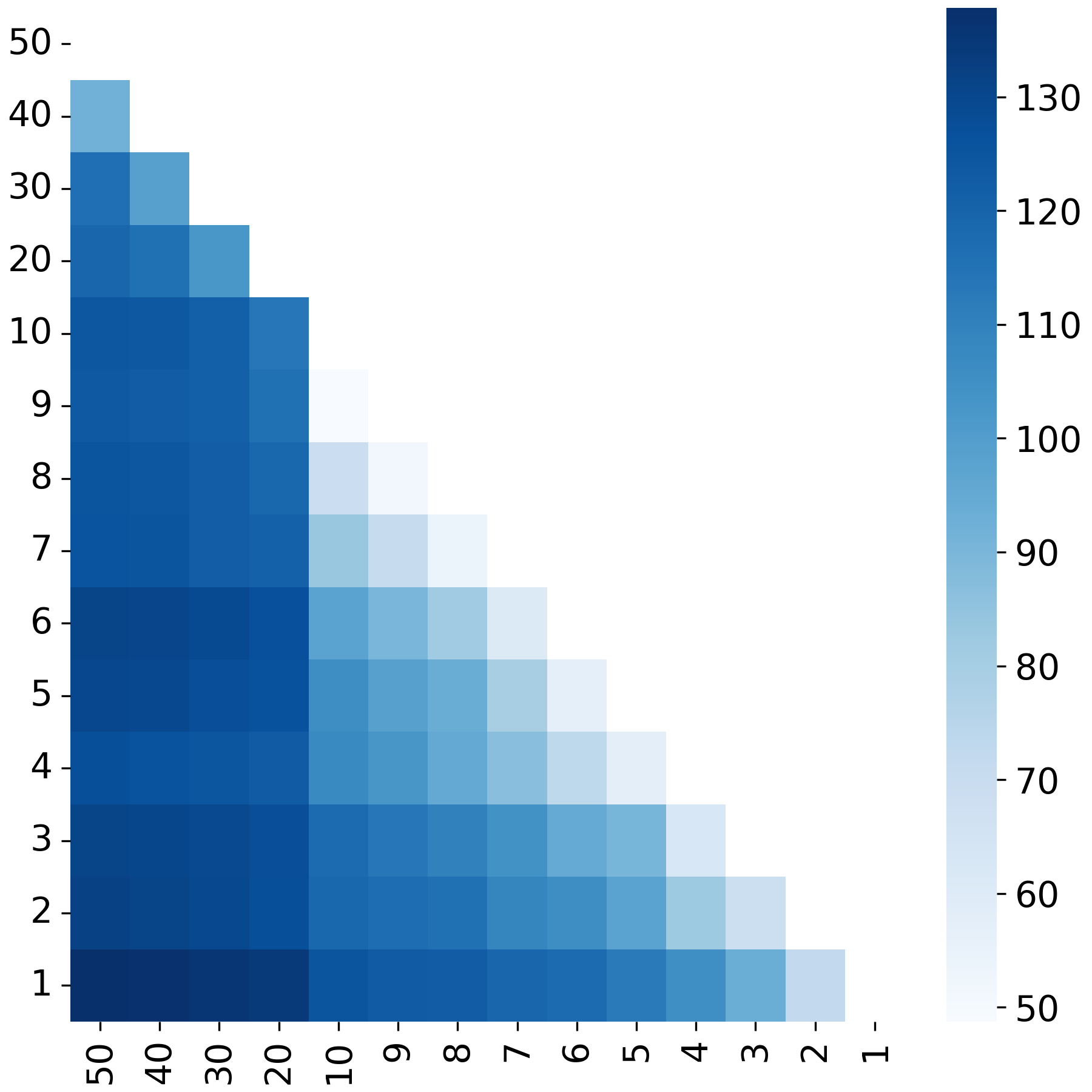

**Figure S21**. Heatmap showing pairwise t-tests of Pearson correlation when restricting data to a uniform read-depth level. The colorbar shows -log10-transformed p-values.

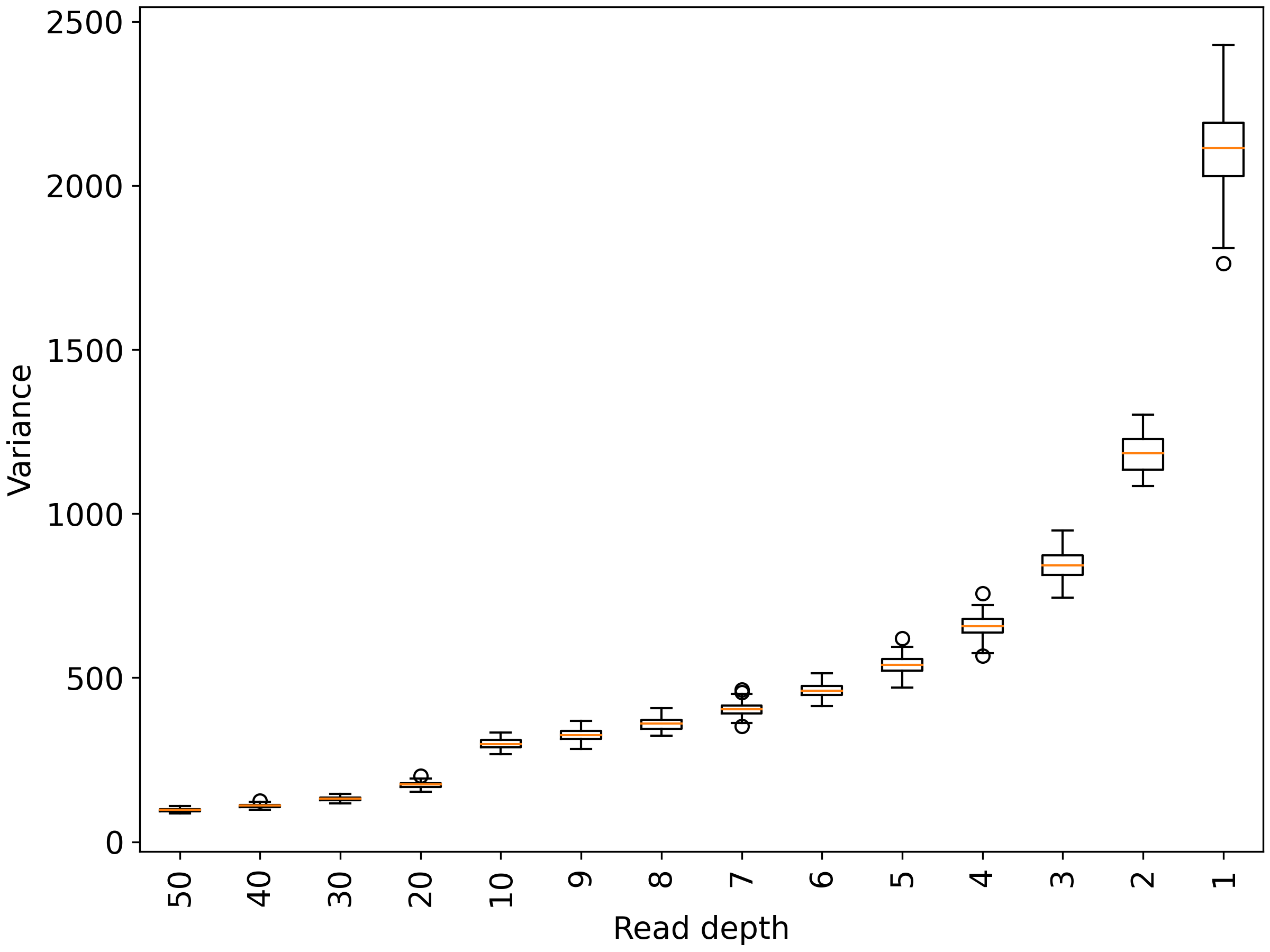

**Figure S22**. Boxplot showing distributions of variance when restricting data to a uniform read-depth level.

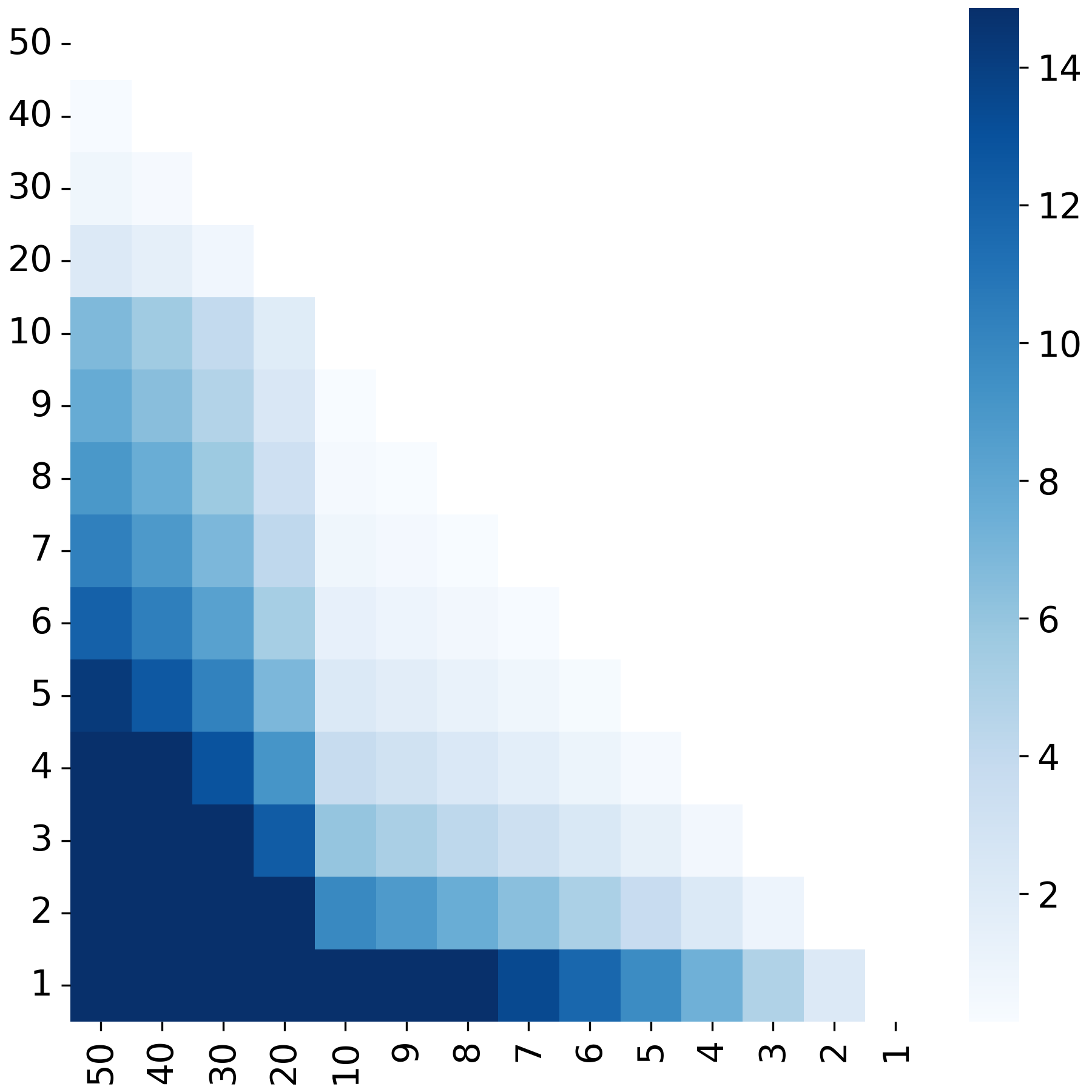

**Figure S23**. Heatmap showing pairwise F-tests of variance when restricting data to a uniform read-depth level. The colorbar shows -log10-transformed p-values.

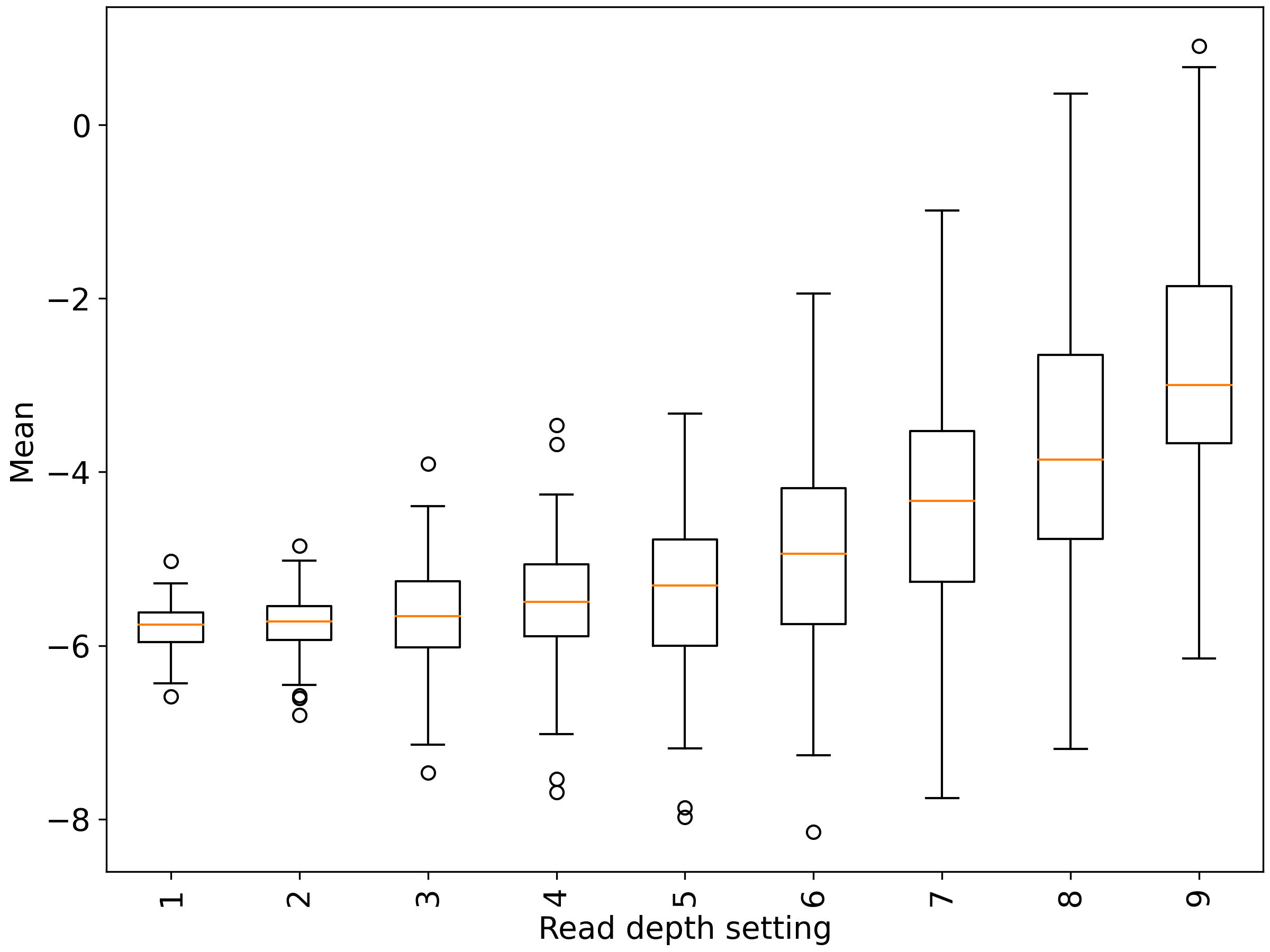

**Figure S24**. Boxplot showing distributions of mean when restricting data to a variable read-depth level. See Figure S2 for details on each read-depth distribution.

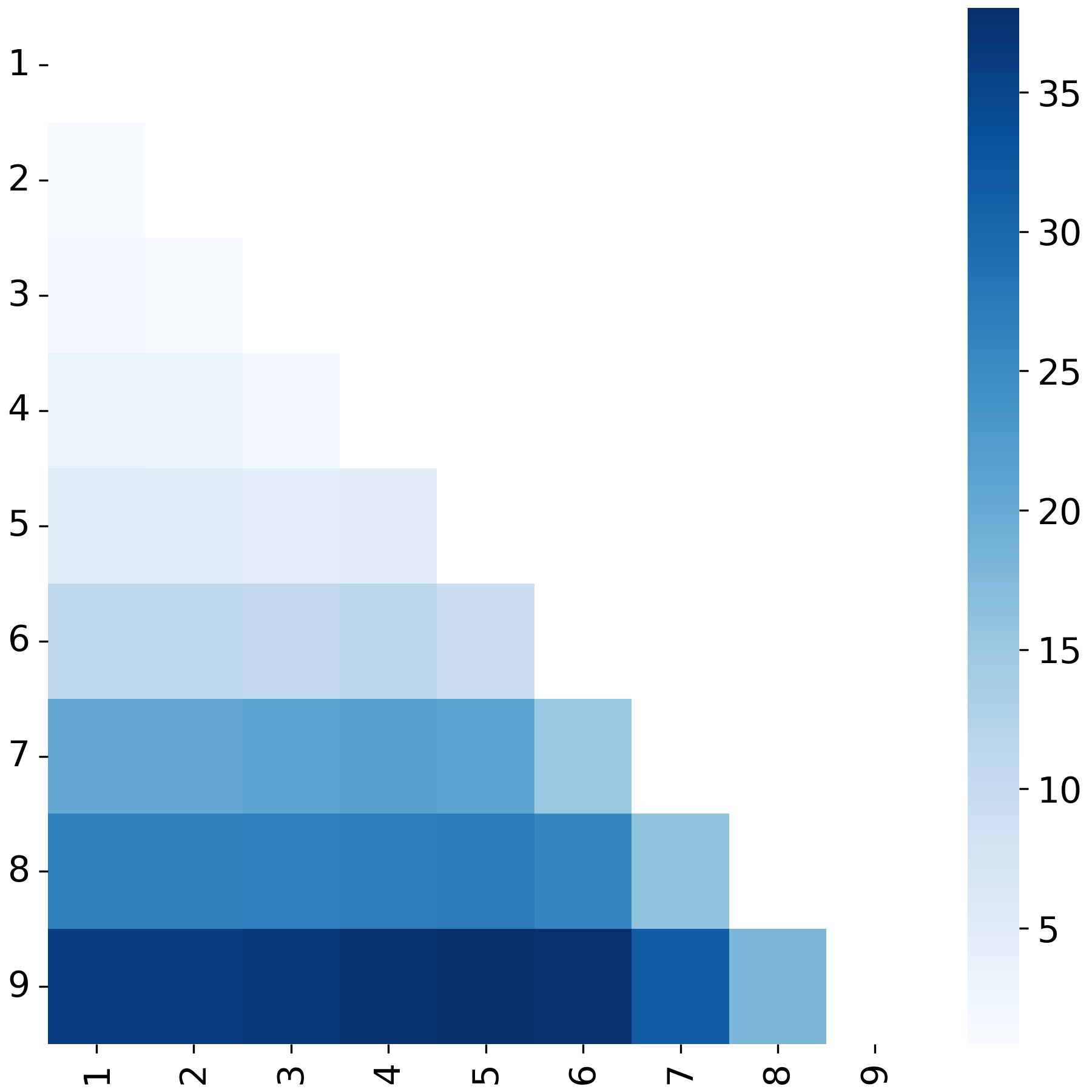

**Figure S25**. Heatmap showing pairwise t-tests of mean when restricting data to a variable read-depth level. The colorbar shows -log10-transformed p-values. See Figure S2 for details on each read-depth distribution.

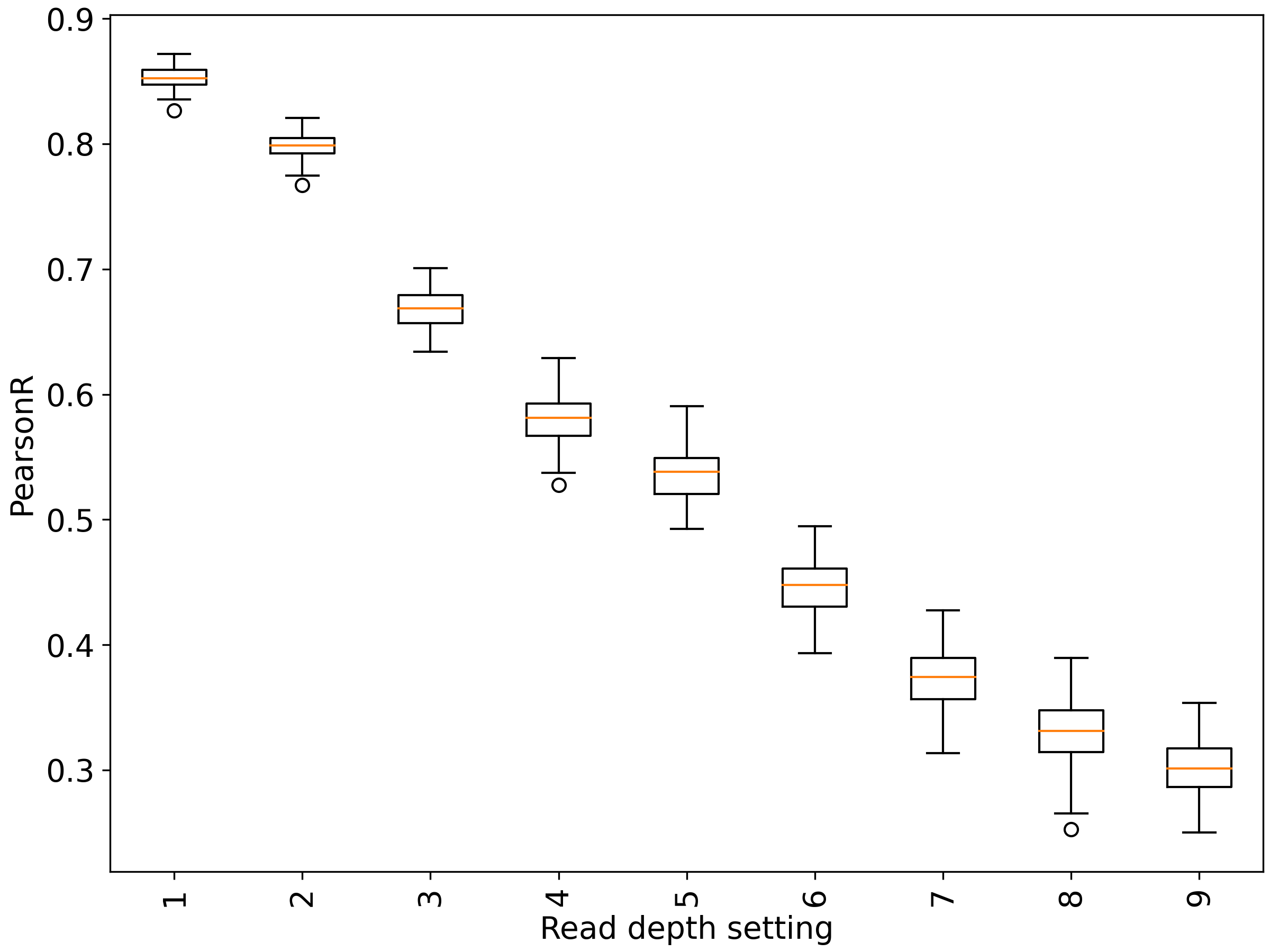

**Figure S26**. Boxplot showing distributions of Pearson correlation when restricting data to a variable read-depth level. See Figure S2 for details on each read-depth distribution.

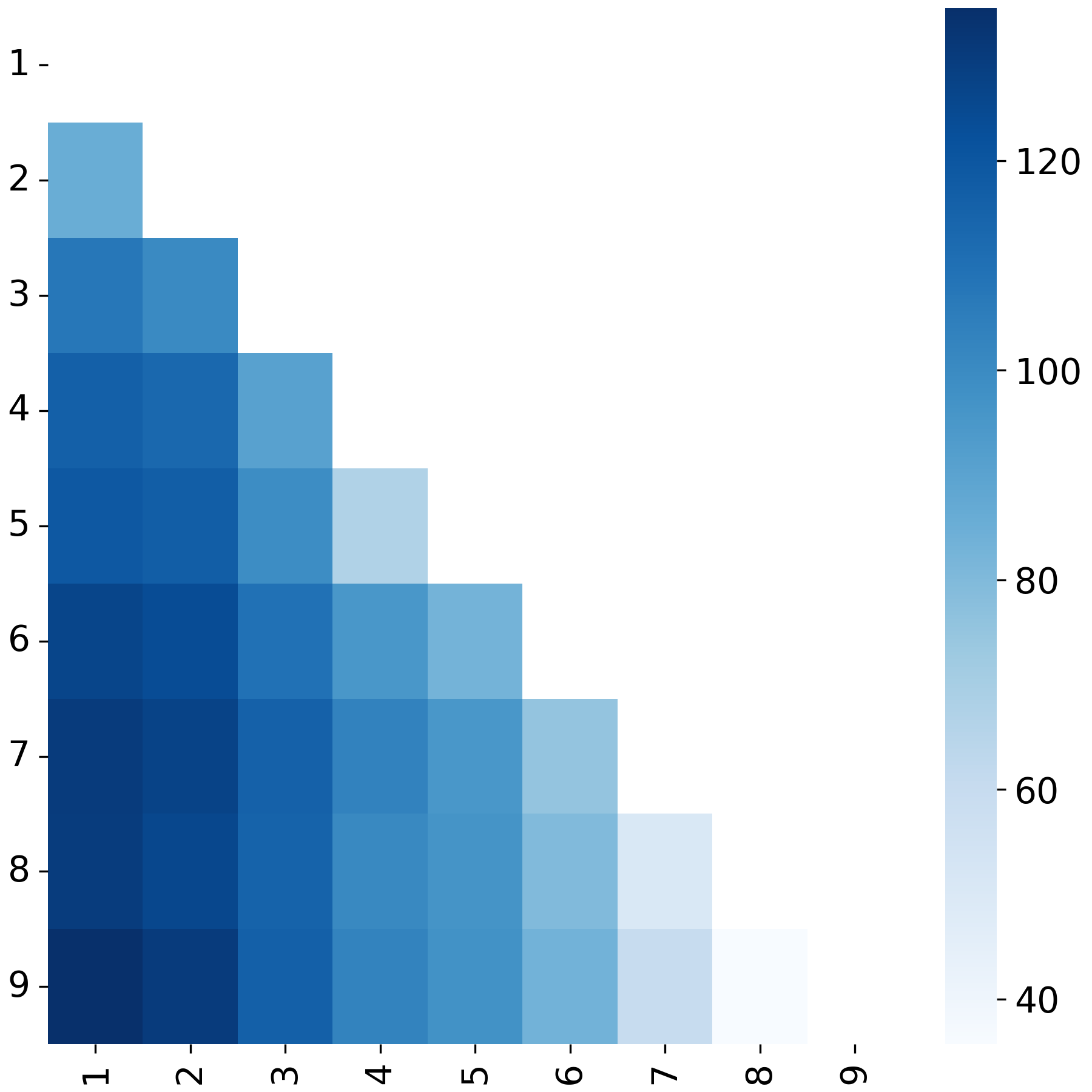

**Figure S27**. Heatmap showing pairwise t-tests of Pearson correlation when restricting data to a variable read-depth level. The colorbar shows -log10-transformed p-values. See Figure S2 for details on each read-depth distribution.

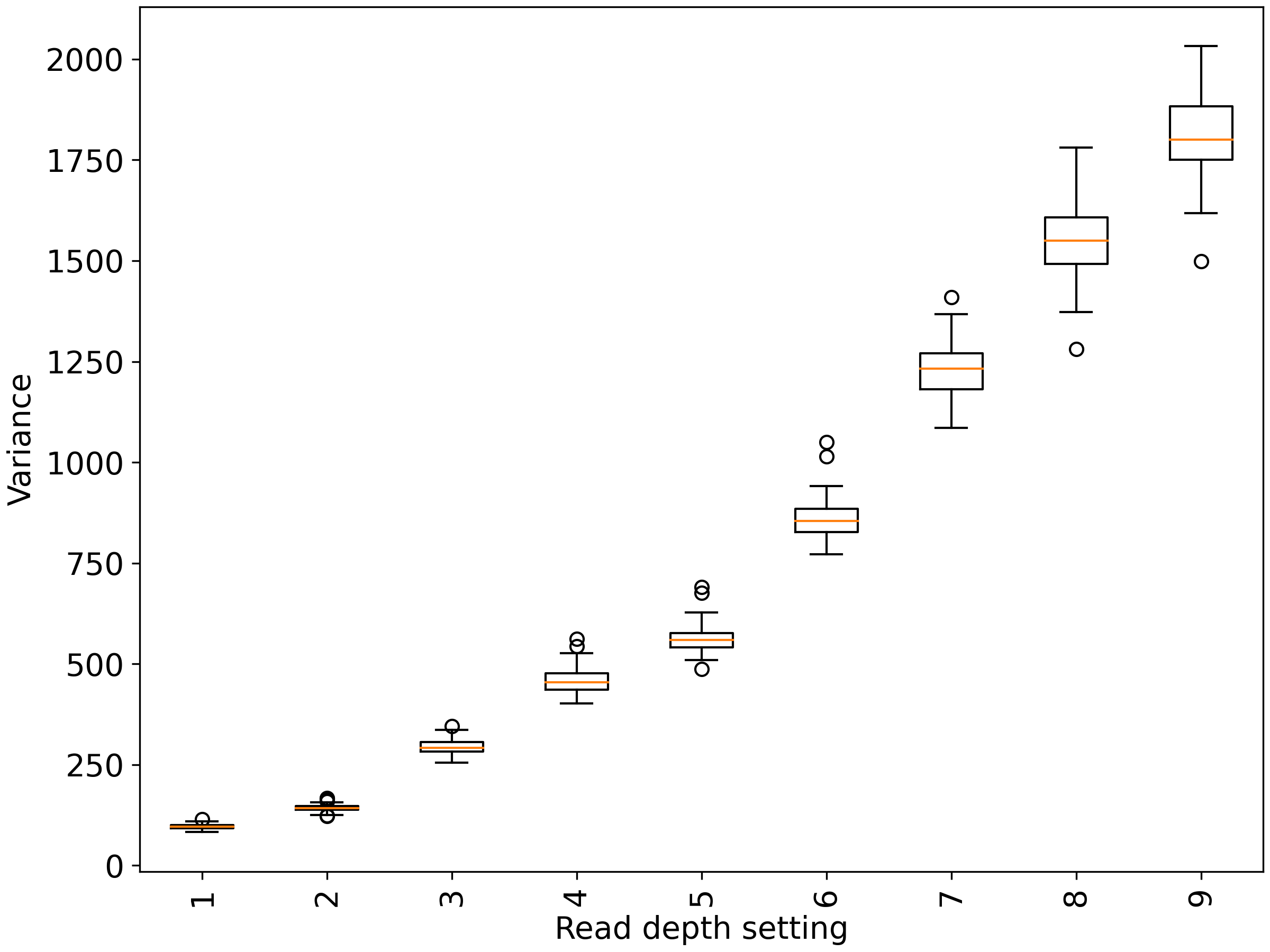

**Figure S28**. Boxplot showing distributions of variance when restricting data to a variable read-depth level. See Figure S2 for details on each read-depth distribution.

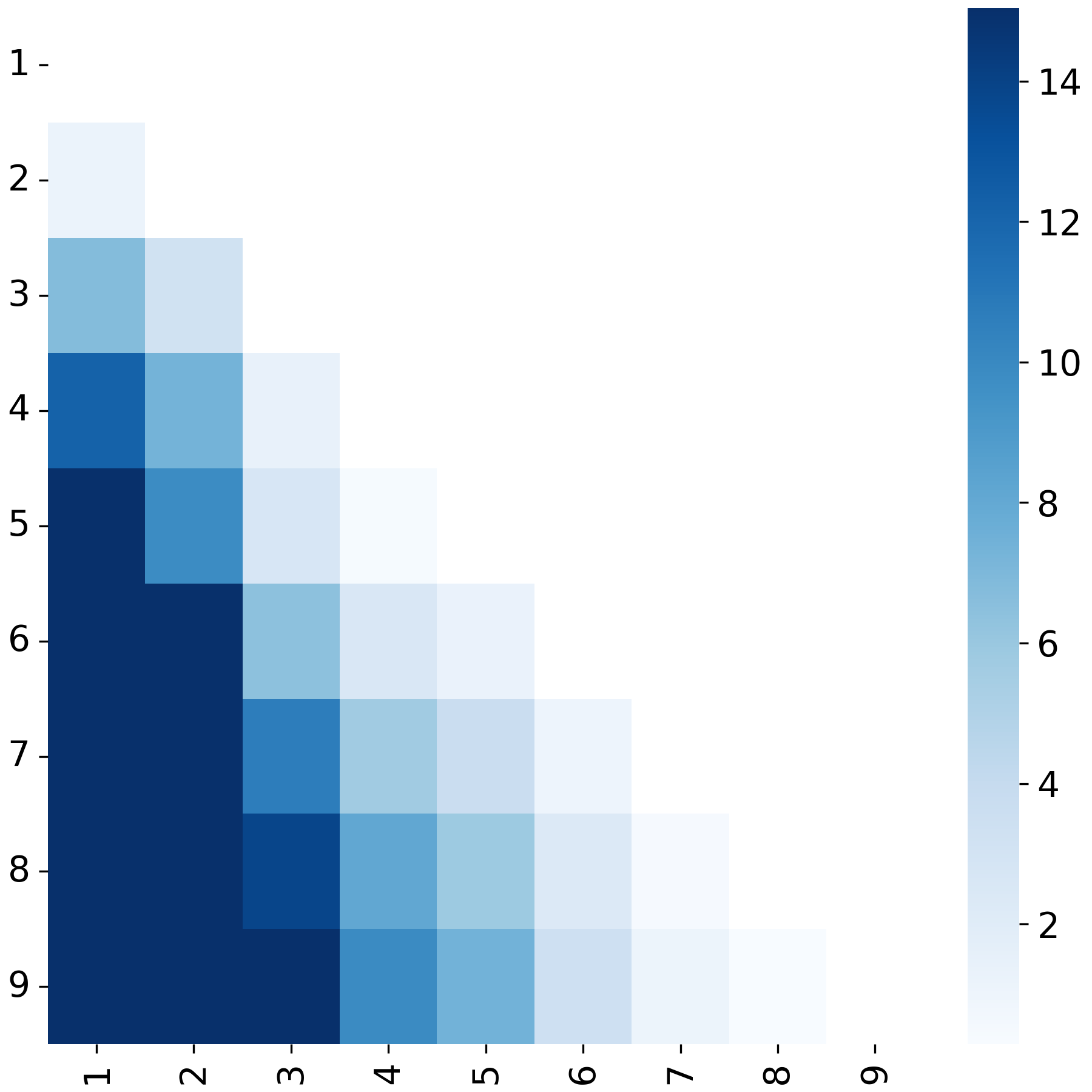

**Figure S29.** Heatmap showing pairwise F-tests of variance when restricting data to a variable read-depth level. The colorbar shows -log10-transformed p-values. See Figure S2 for details on each read-depth distribution.

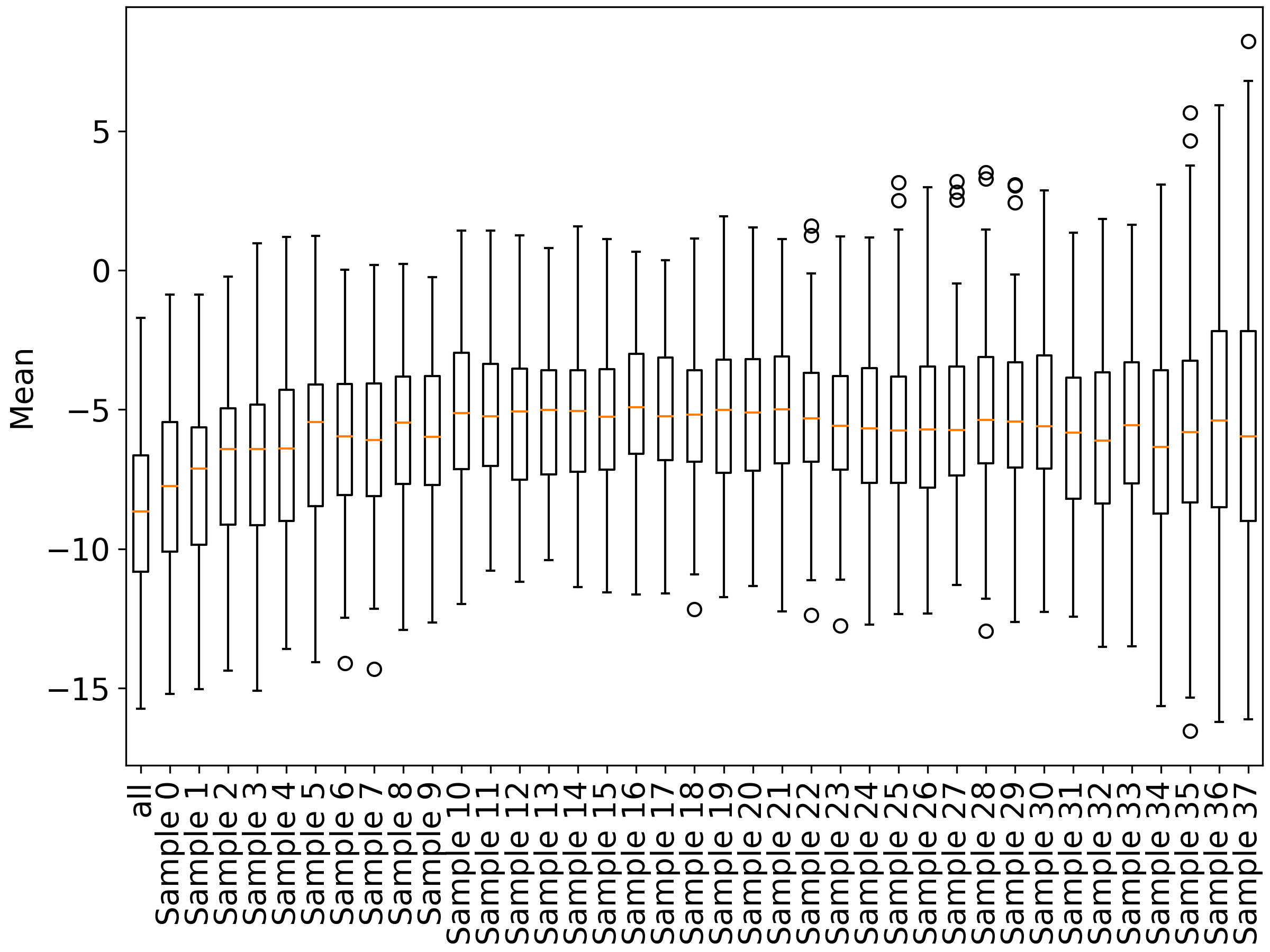

**Figure S30**. Boxplot showing distributions of mean when removing lowest-quality samples as ranked by Pearson correlation.

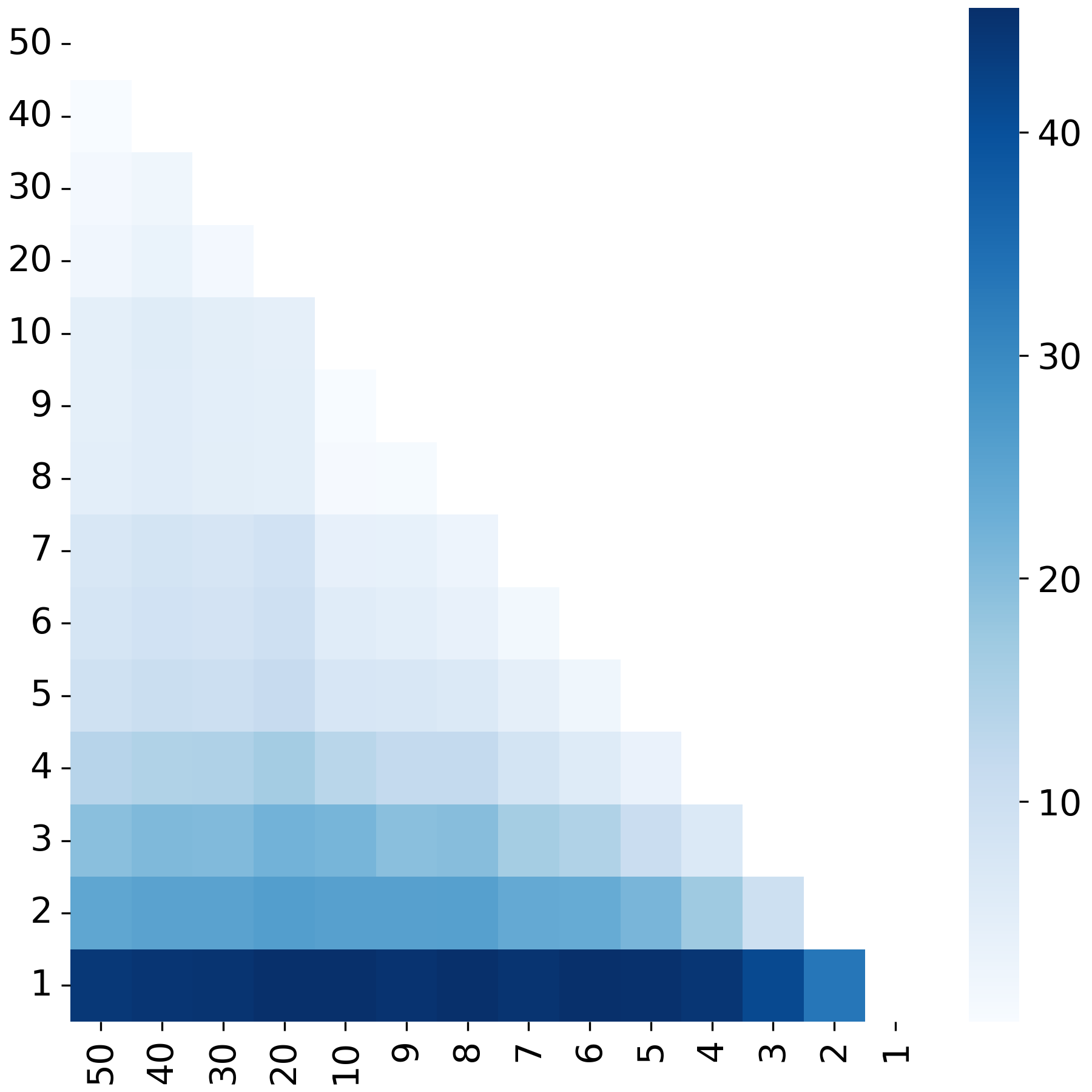

**Figure S31.** Heatmap showing pairwise t-tests of mean when removing lowest-quality samples as ranked by Pearson correlation. The colorbar shows -log10-transformed p-values.

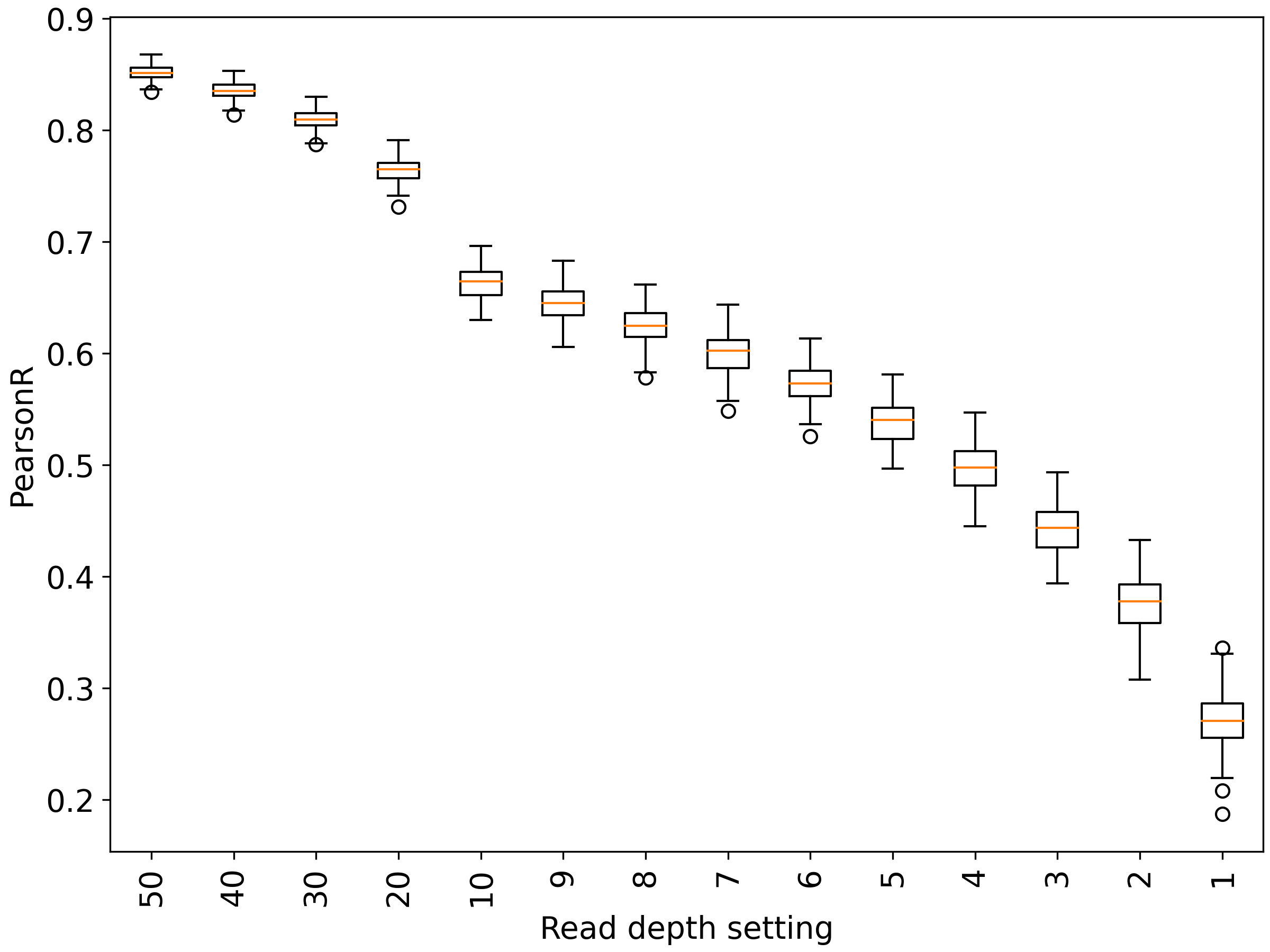

**Figure S32.** Boxplot showing distributions of Pearson correlation when removing lowest-quality samples as ranked by Pearson correlation.

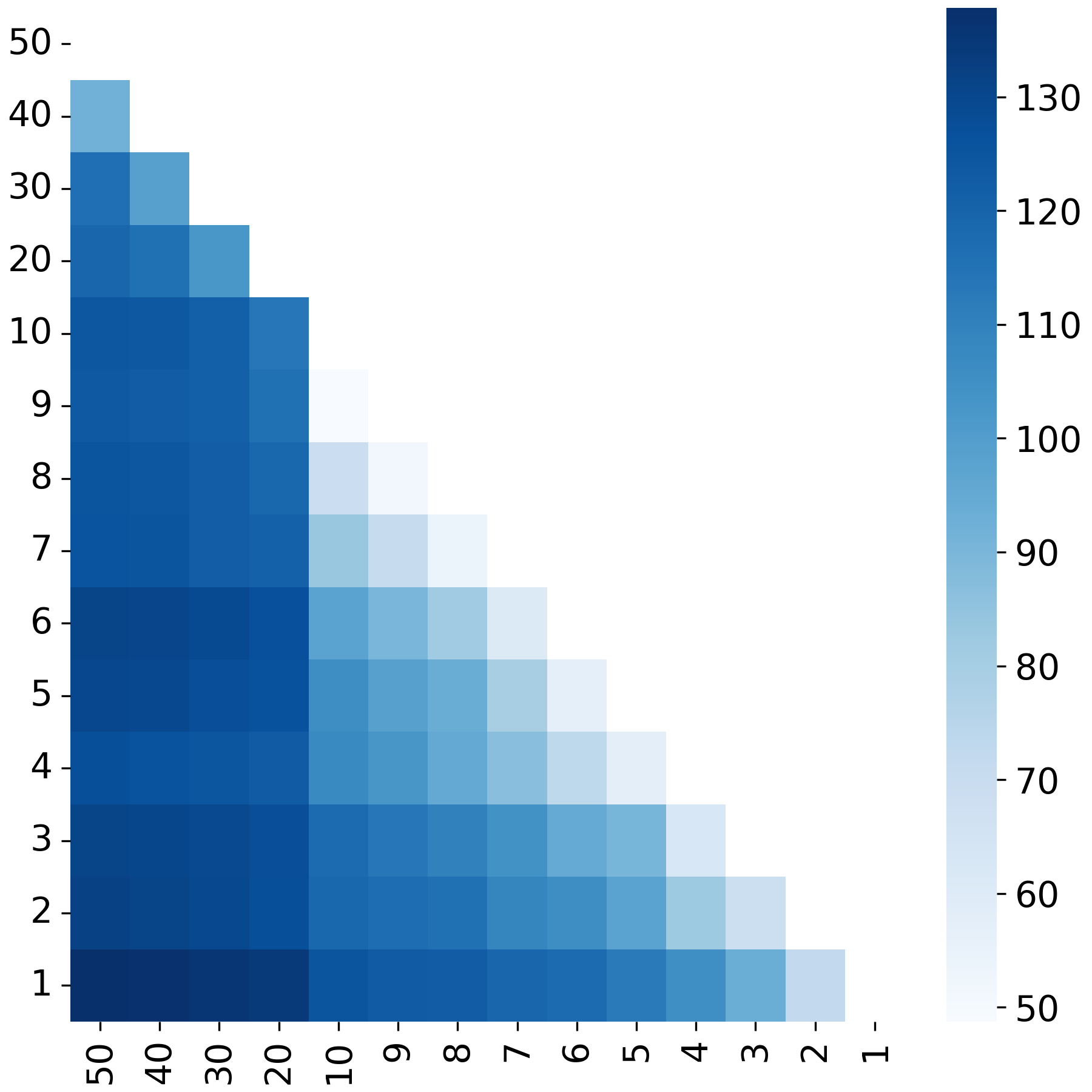

**Figure S33.** Heatmap showing pairwise t-tests of Pearson correlation when removing lowest-quality samples as ranked by Pearson correlation. The colorbar shows -log10-transformed p-values.

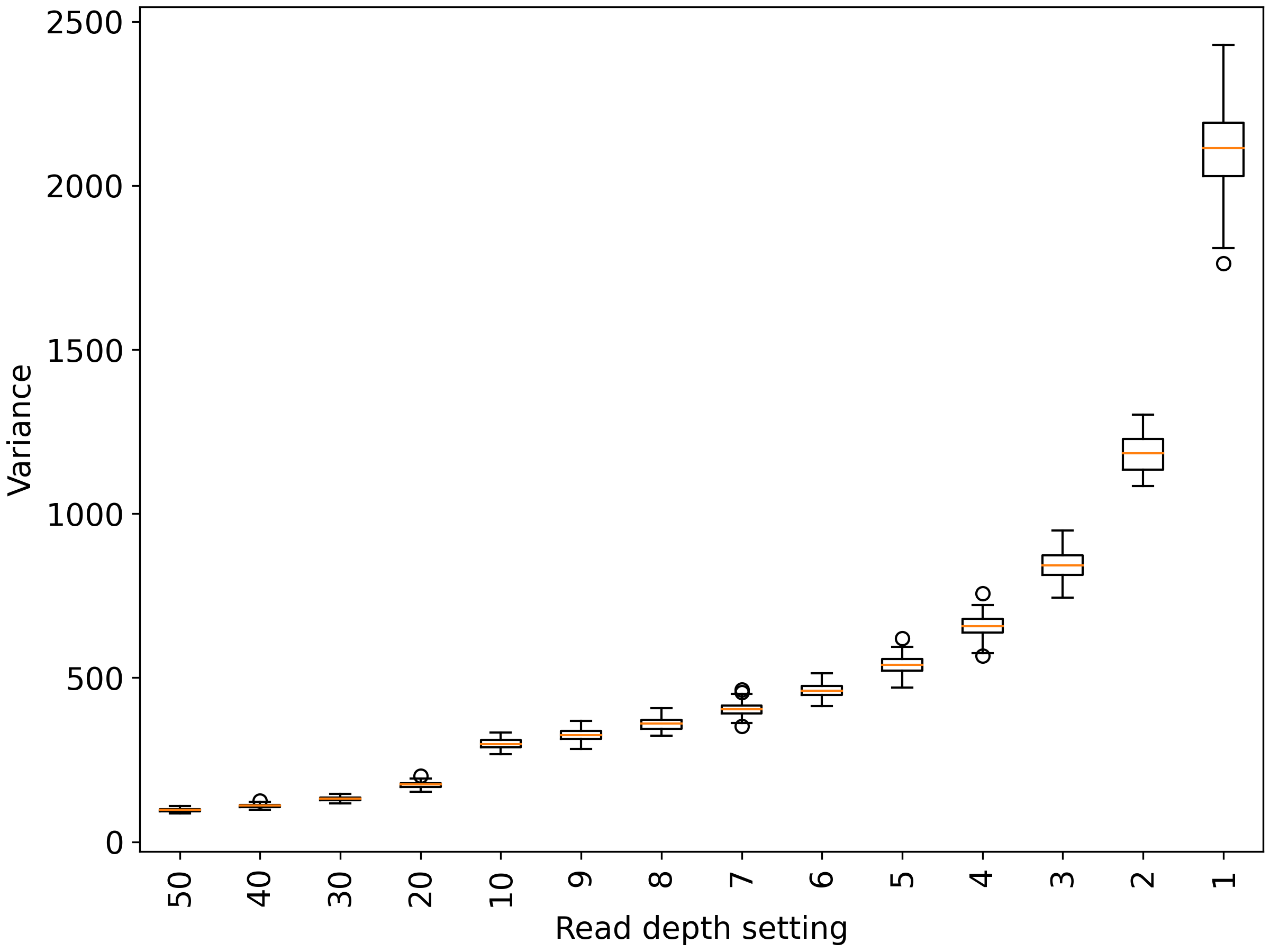

**Figure S34.** Boxplot showing distributions of variance when removing lowest-quality samples as ranked by Pearson correlation.

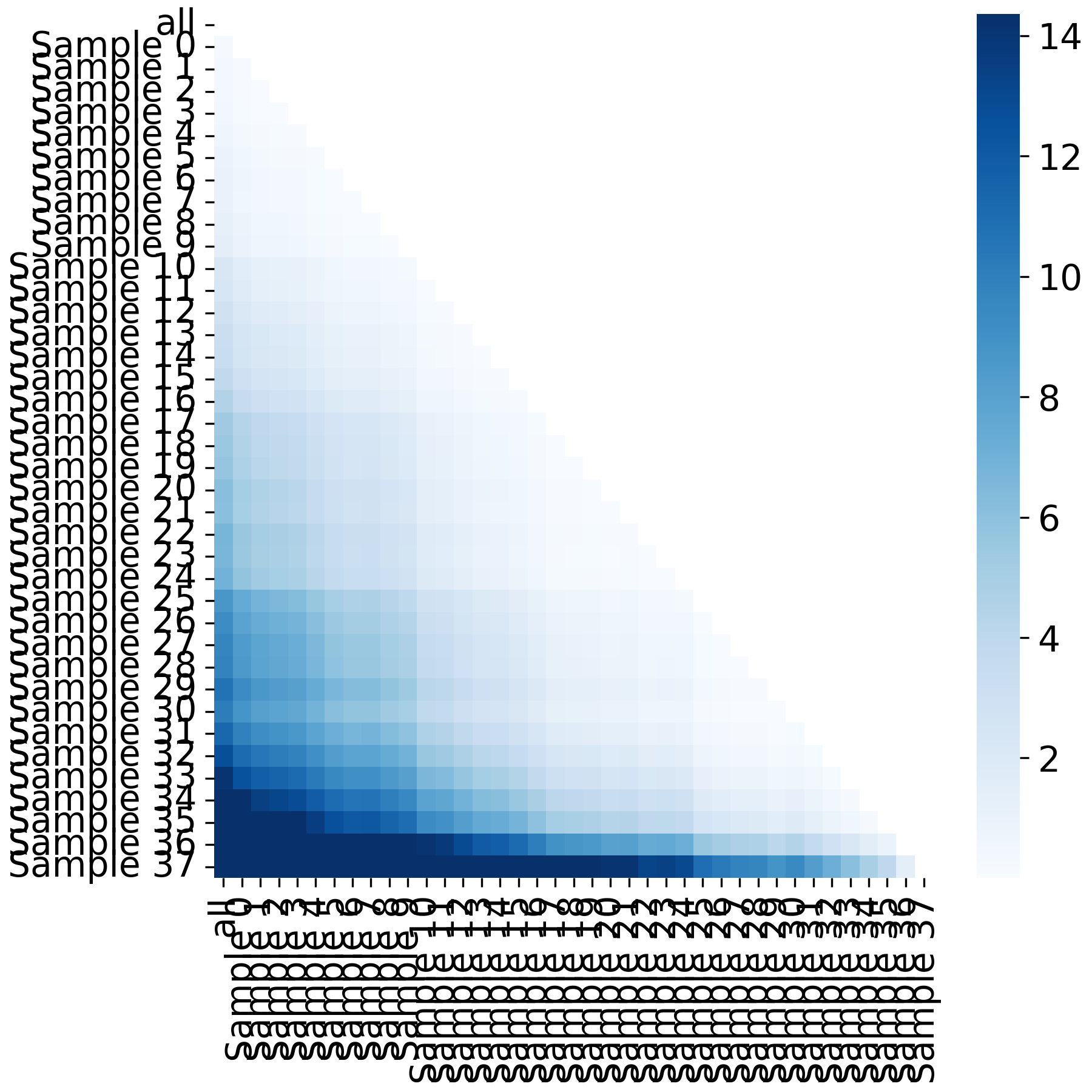

**Figure S35.** Heatmap showing pairwise F-tests of variance when removing lowest-quality samples as ranked by Pearson correlation. The colorbar shows -log10-transformed p-values.

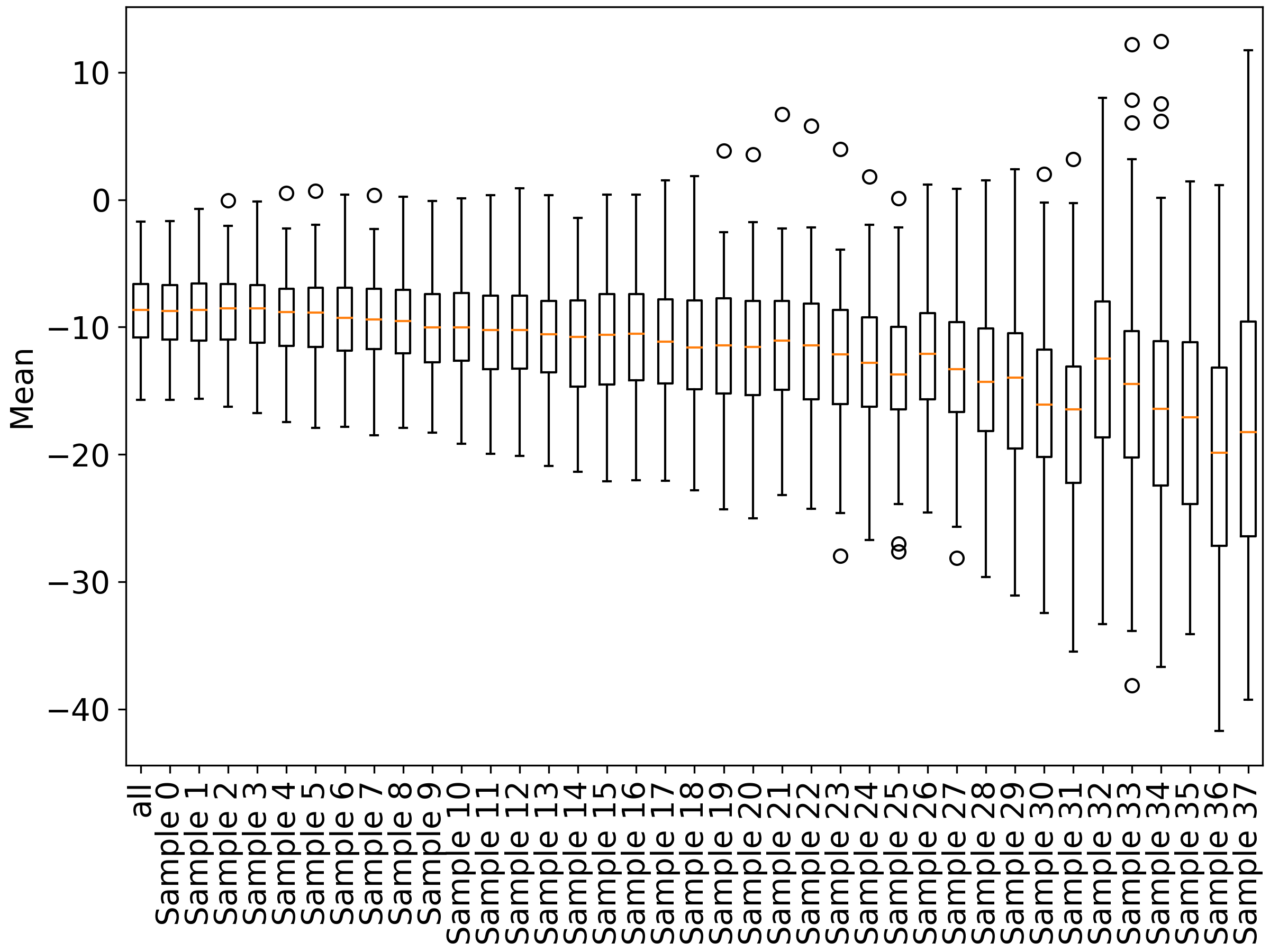

**Figure S36**. Boxplot showing distributions of mean when removing lowest-quality samples ranked randomly.

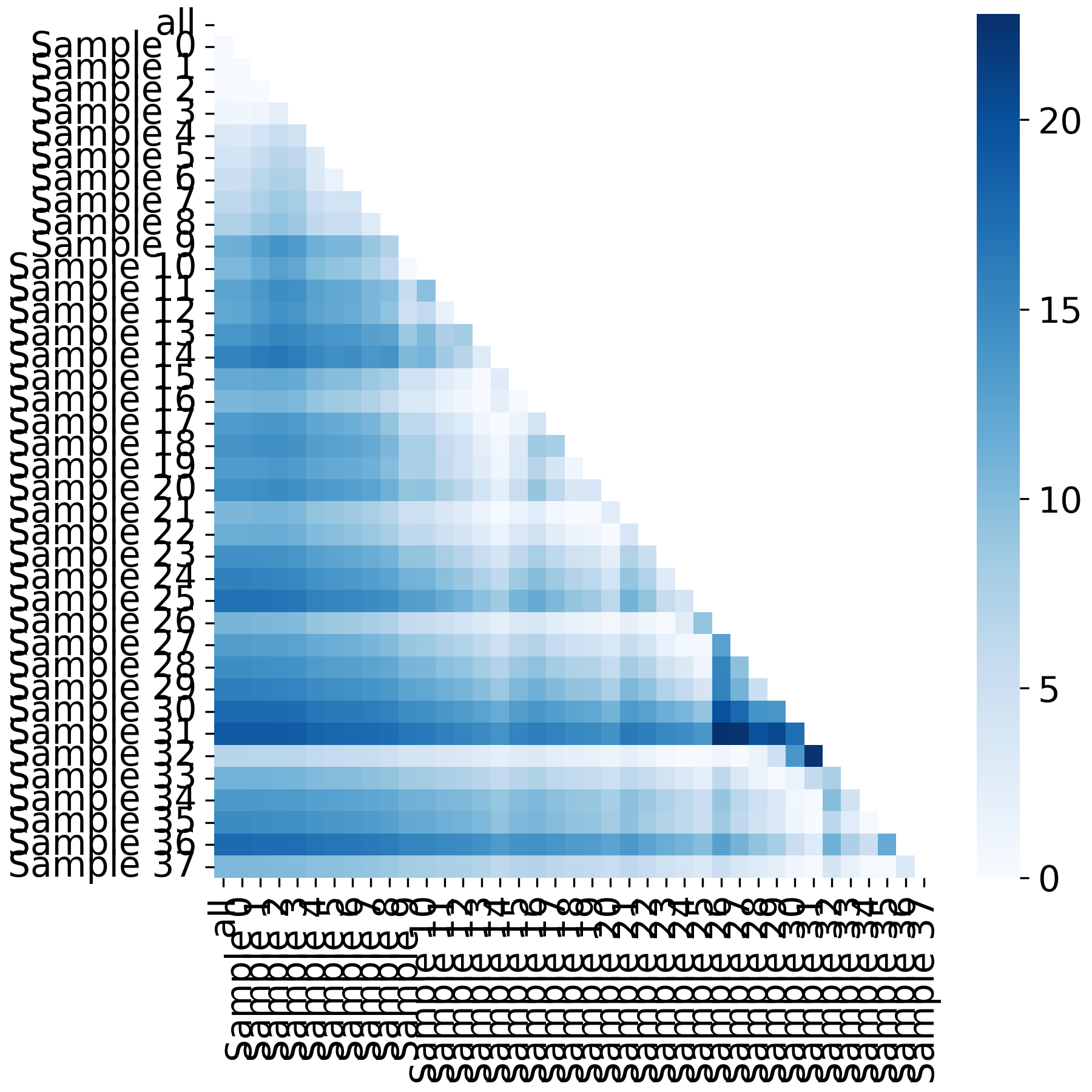

**Figure S37.** Heatmap showing pairwise t-tests of mean when removing lowest-quality samples ranked randomly. The colorbar shows -log10-transformed p-values.

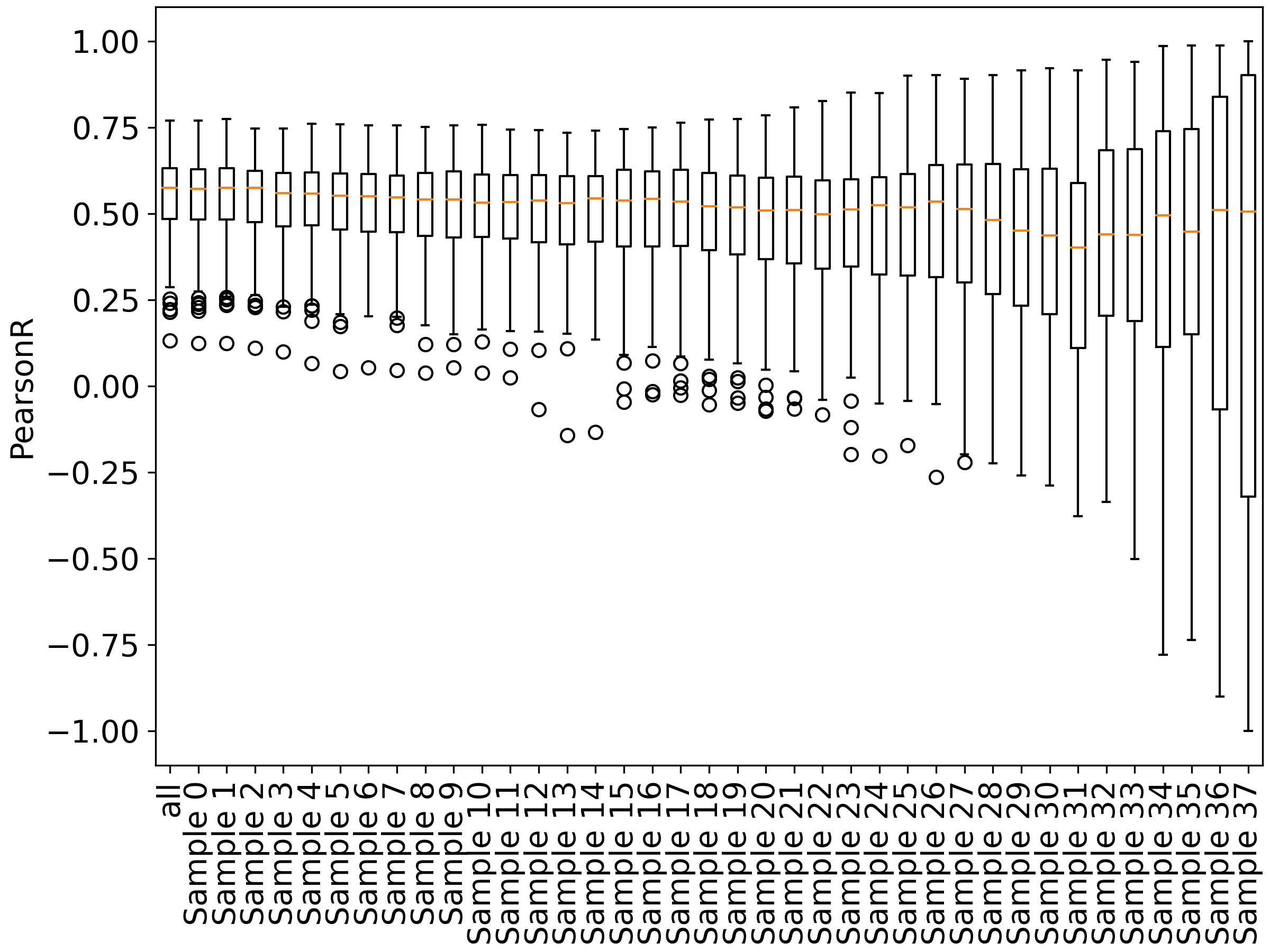

**Figure S38.** Boxplot showing distributions of Pearson correlation when removing lowest-quality samples ranked randomly.

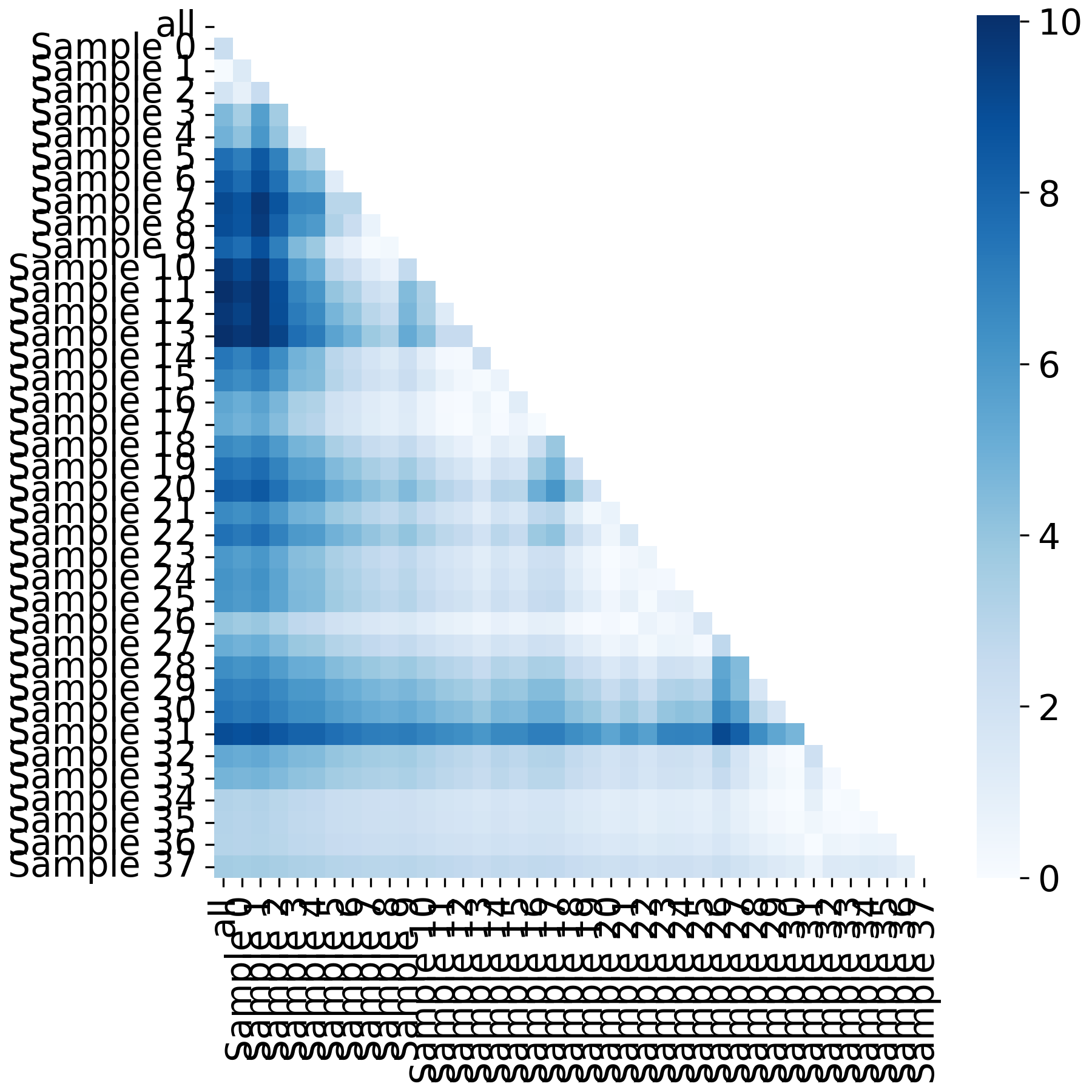

**Figure S39.** Heatmap showing pairwise t-tests of Pearson correlation when removing lowest-quality samples ranked randomly. The colorbar shows -log10-transformed p-values.

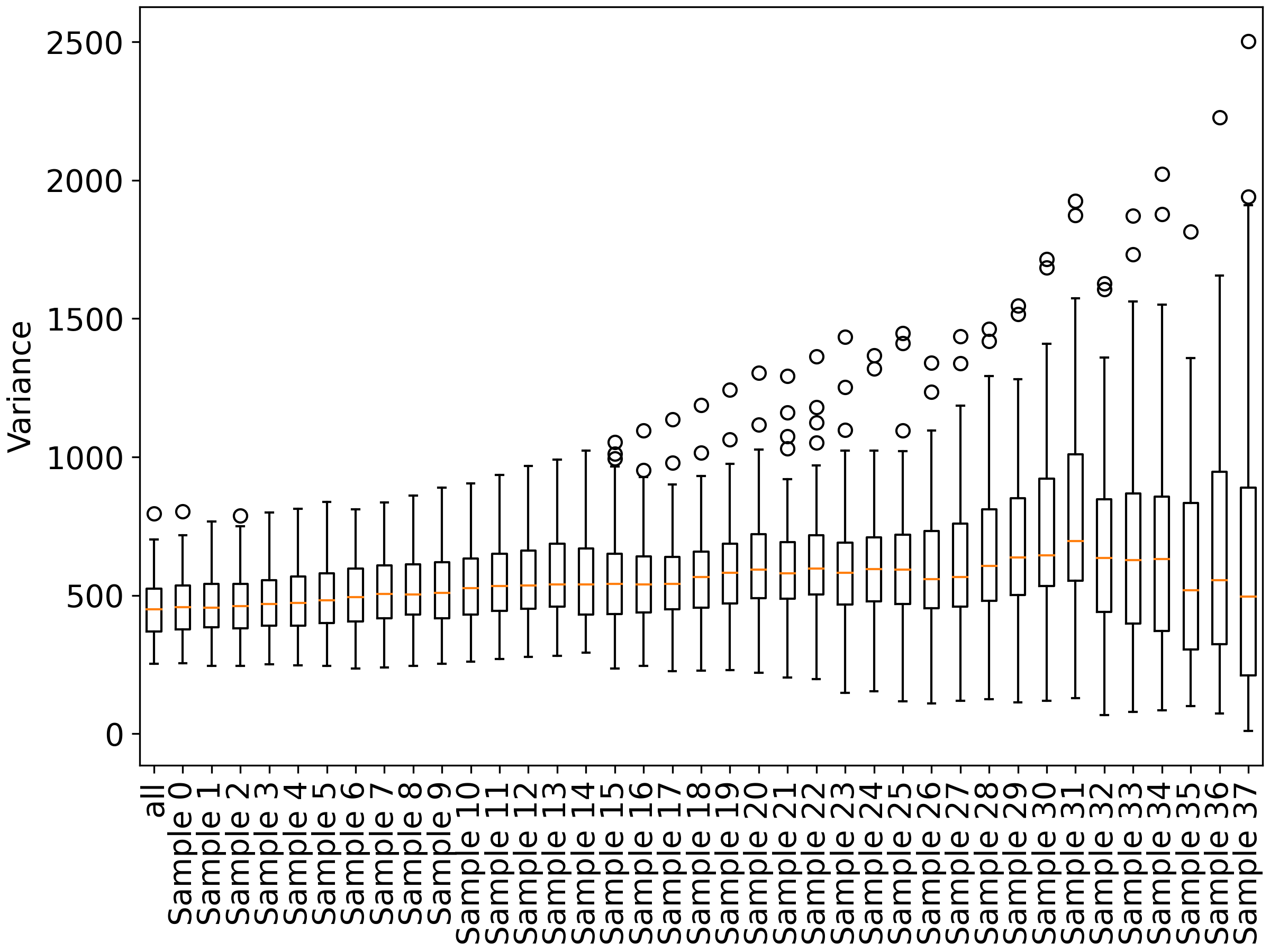

**Figure S40.** Boxplot showing distributions of variance when removing lowest-quality samples ranked randomly.

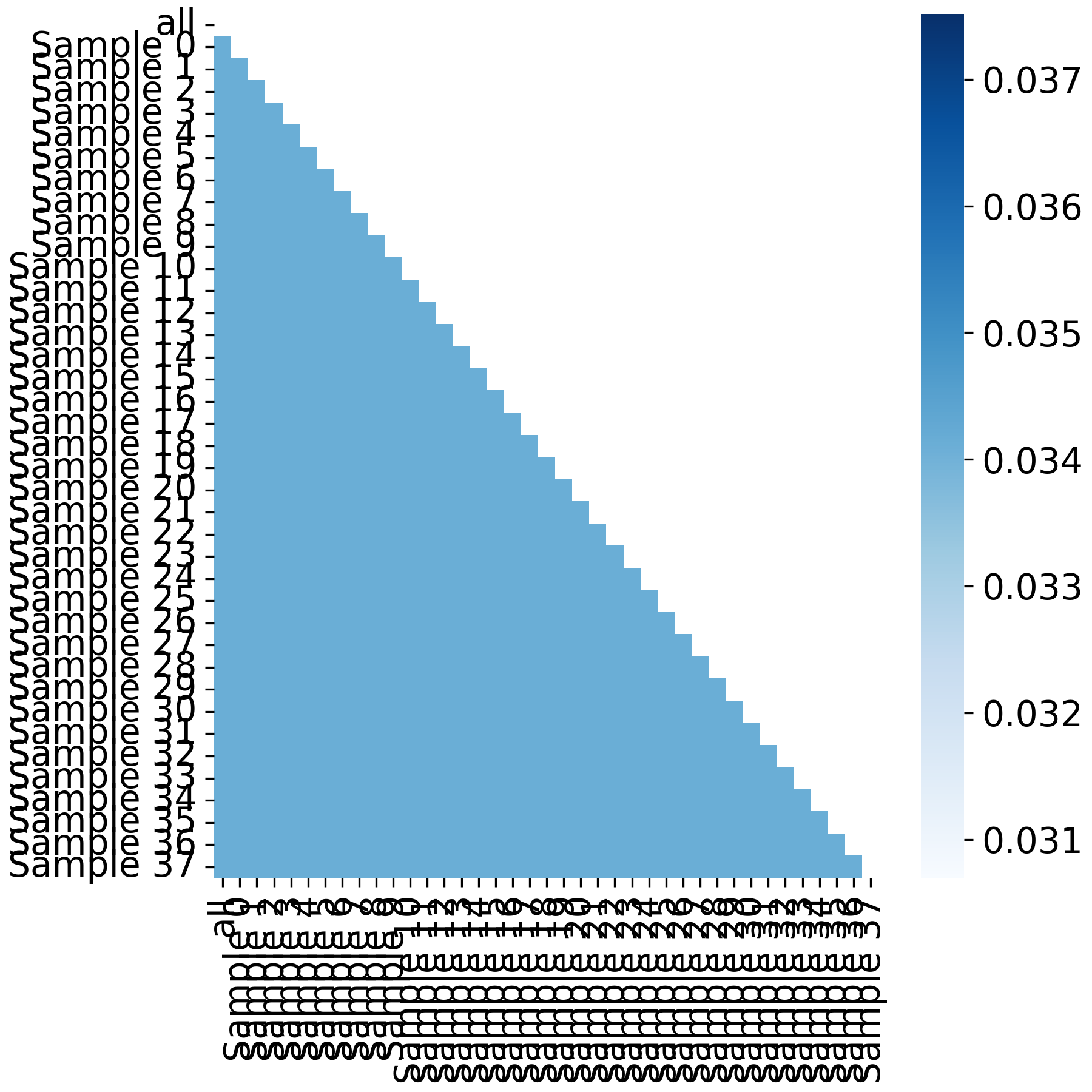

**Figure S41.** Heatmap showing pairwise F-tests of variance when removing lowest-quality samples ranked randomly. The colorbar shows -log10-transformed p-values.

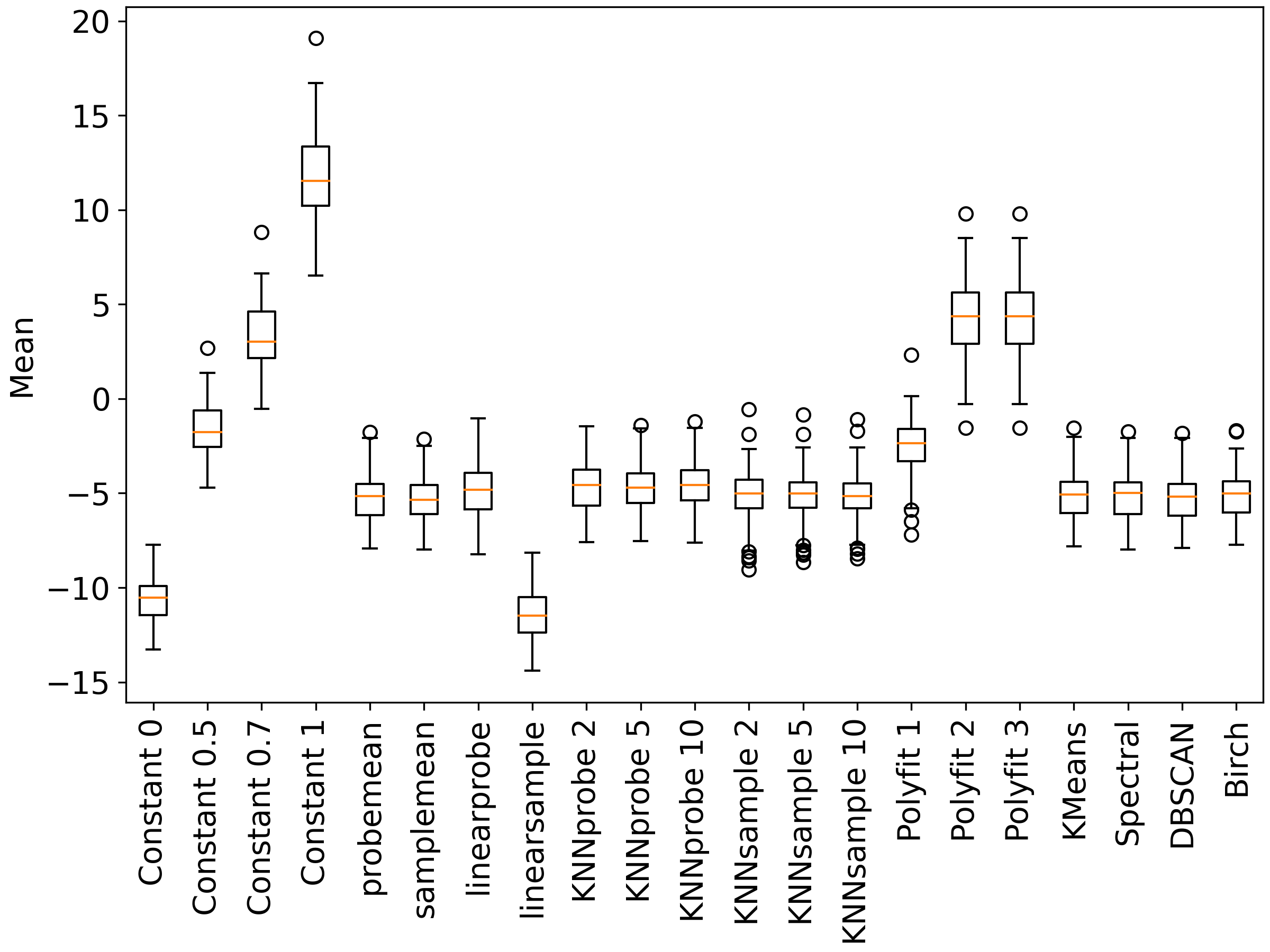

**Figure S42**. Boxplot showing distributions of mean on a moderately perturbed dataset that is imputed by various first-principle imputation tools.

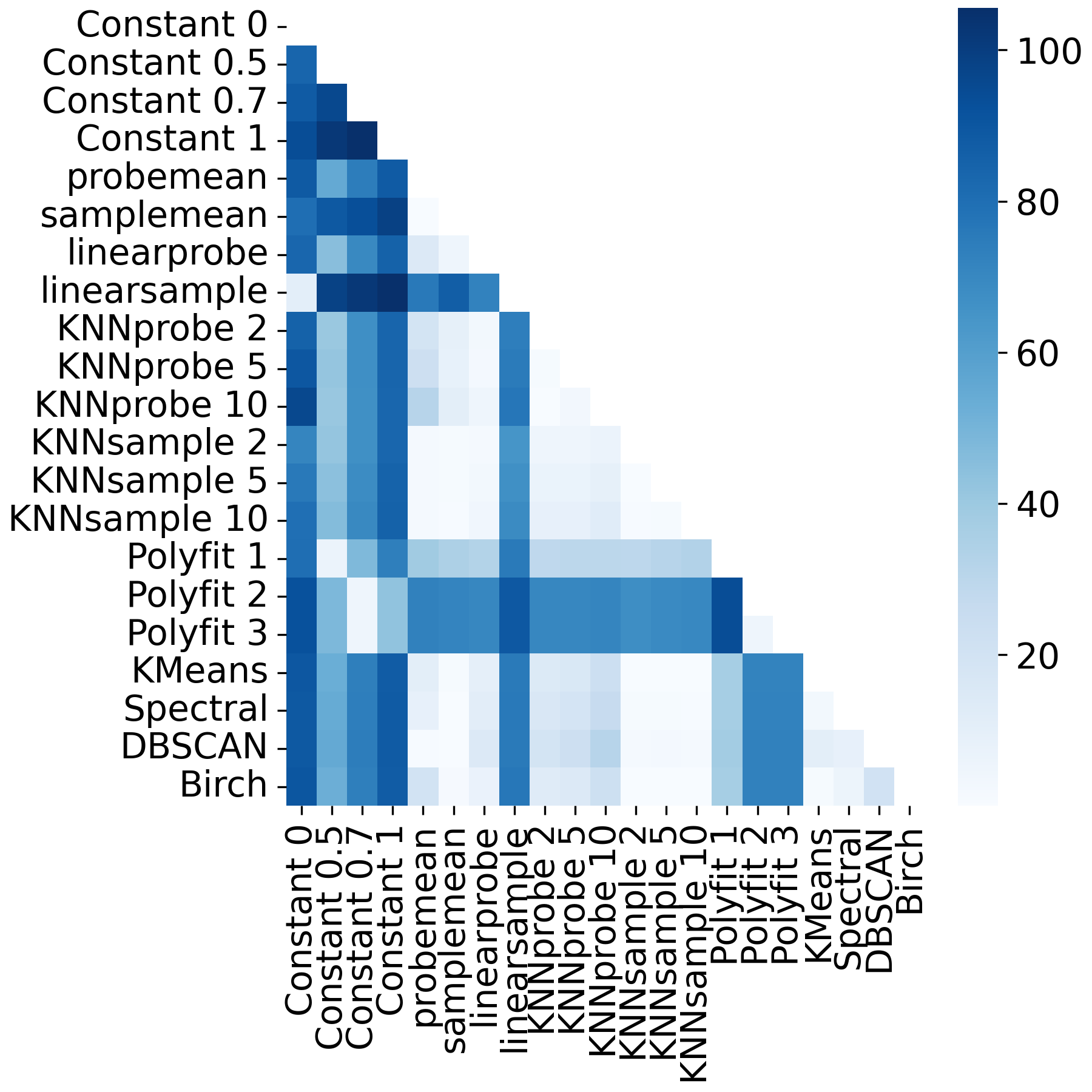

**Figure S43.** Heatmap showing pairwise t-tests of mean on a moderately perturbed dataset that is imputed by various first-principle imputation tools. The colorbar shows -log10-transformed p-values.

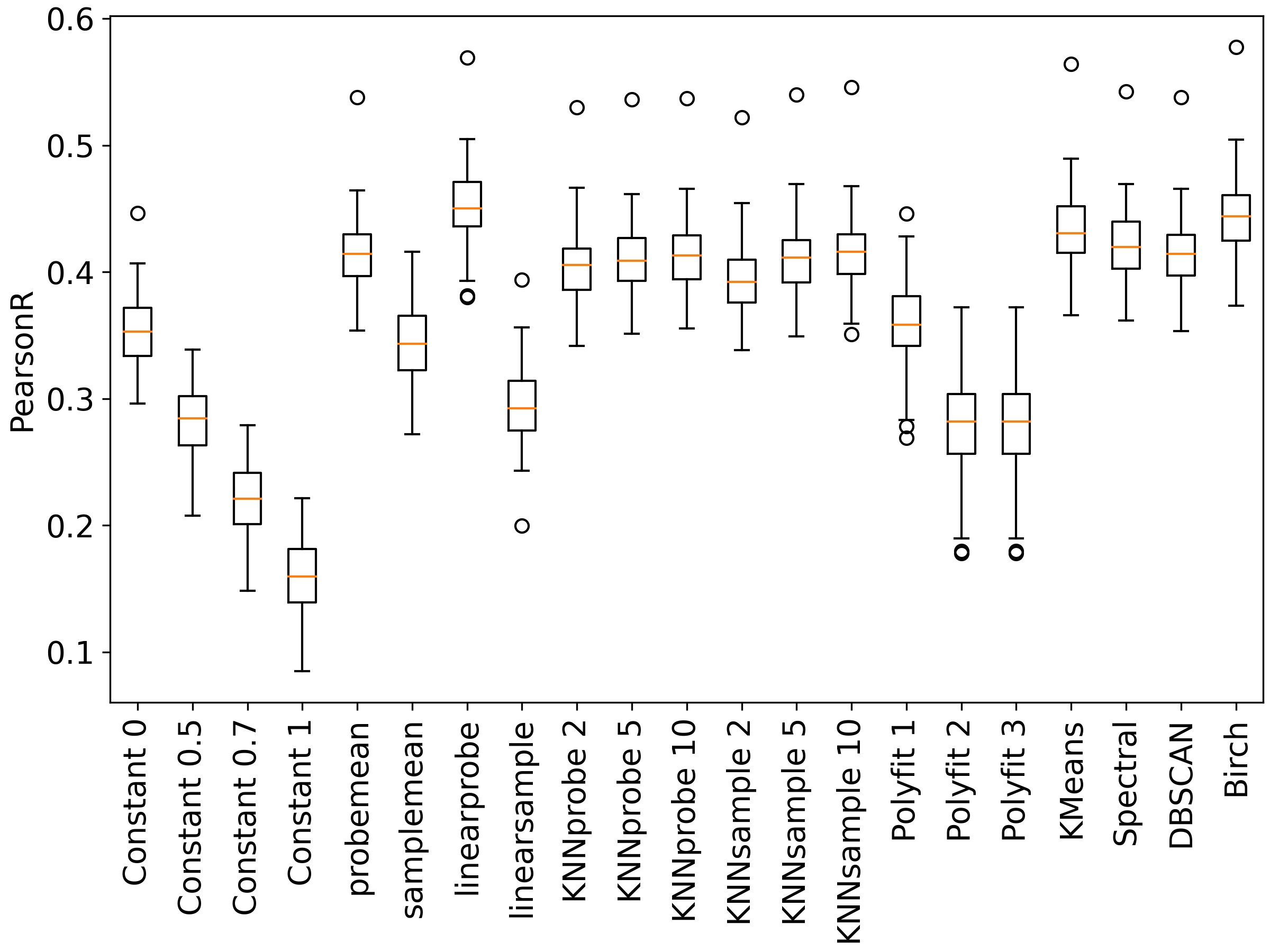

**Figure S44.** Boxplot showing distributions of Pearson correlation on a moderately perturbed dataset that is imputed by various first-principle imputation tools.

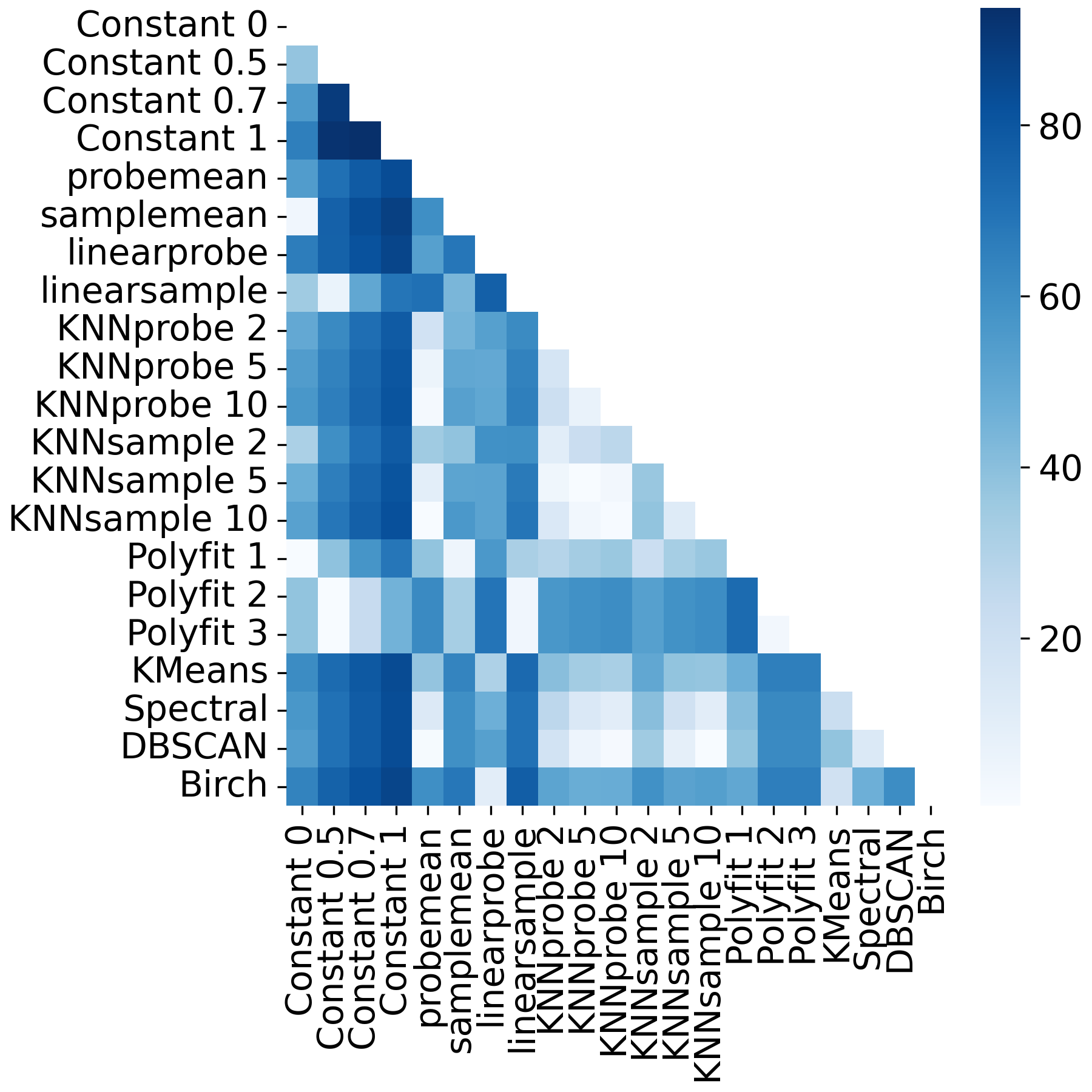

**Figure S45.** Heatmap showing pairwise t-tests of Pearson correlation on a moderately perturbed dataset that is imputed by various first-principle imputation tools. The colorbar shows -log10-transformed p-values.

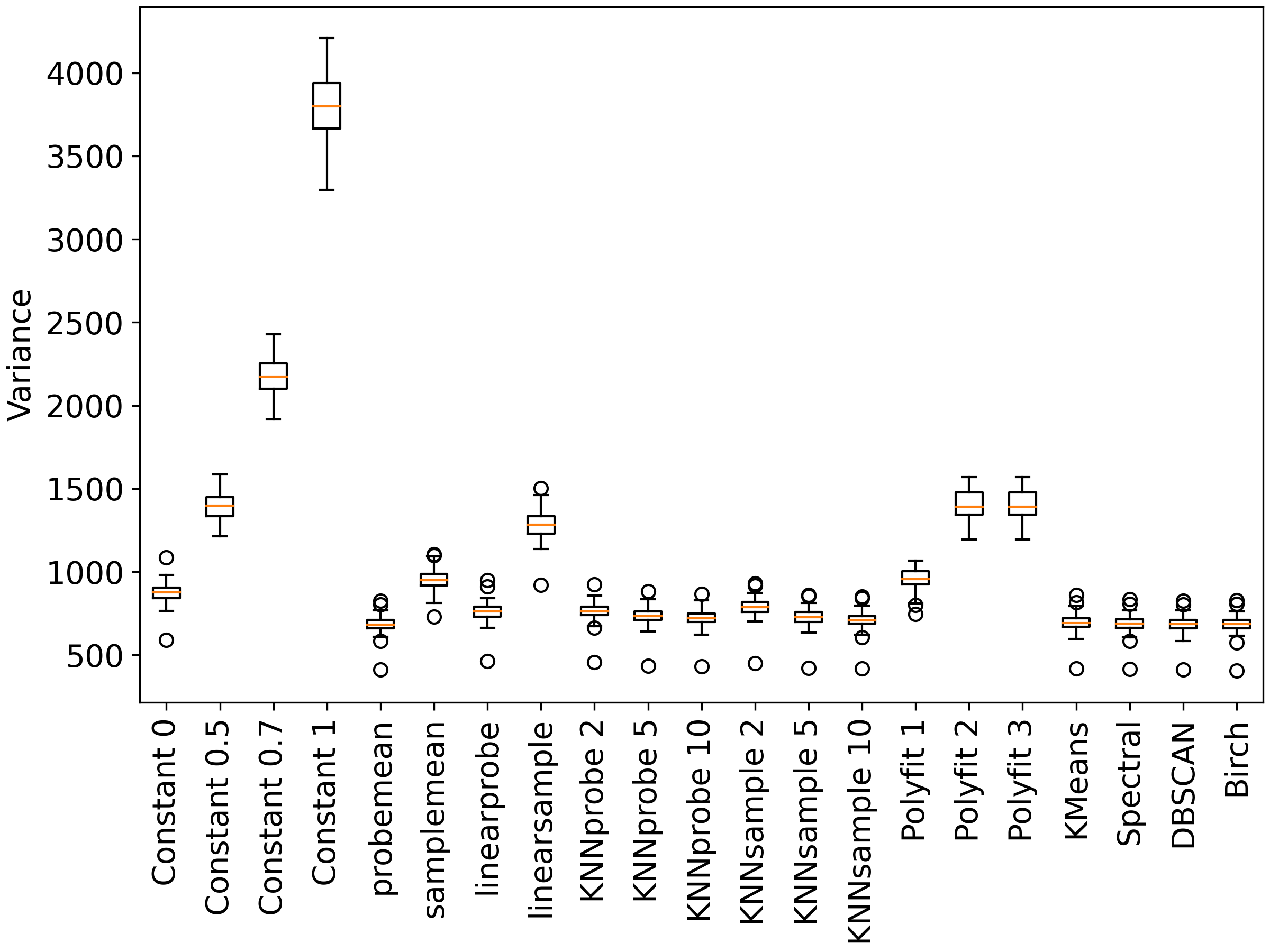

**Figure S46.** Boxplot showing distributions of variance on a moderately perturbed dataset that is imputed by various first-principle imputation tools.

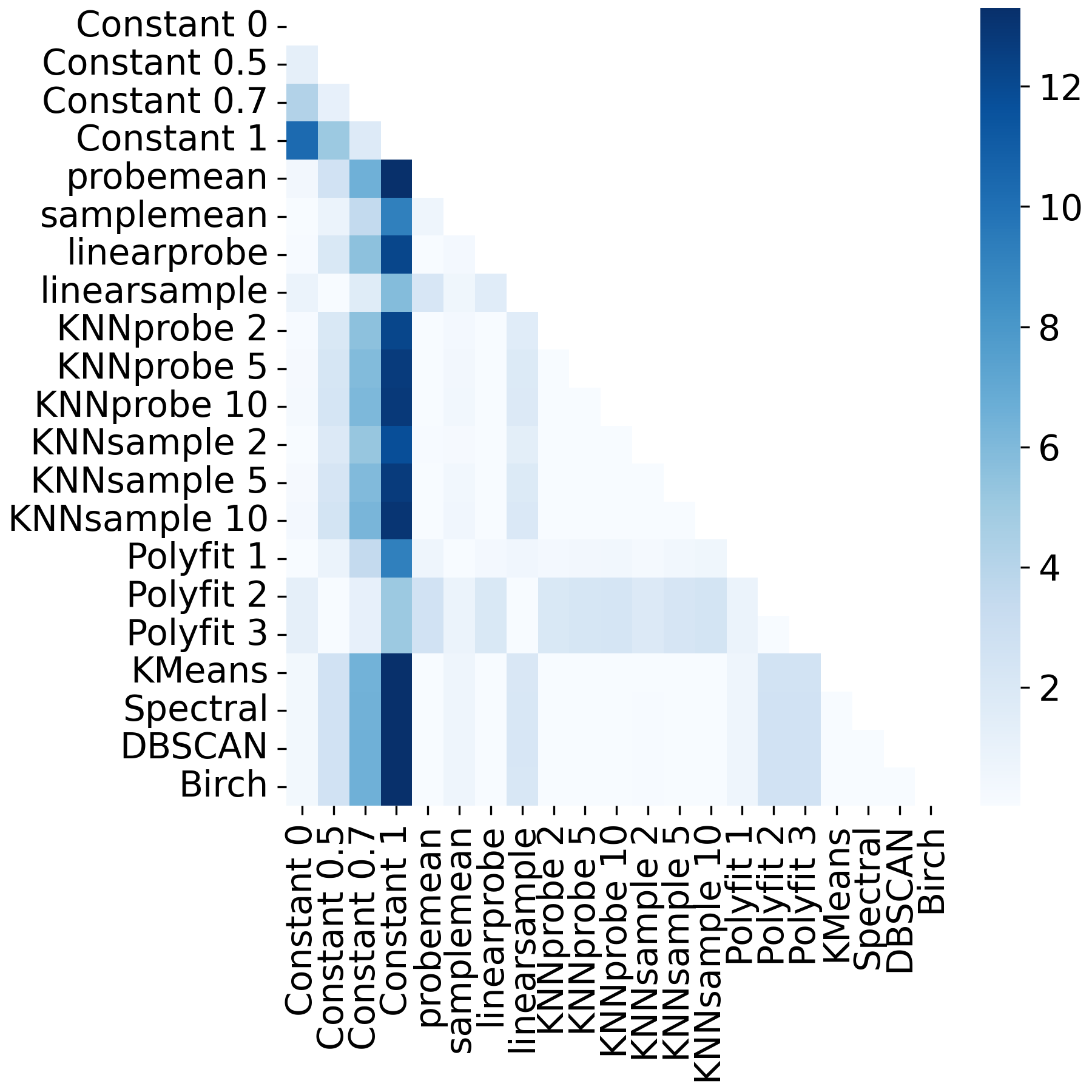

**Figure S47.** Heatmap showing pairwise F-tests of variance on a moderately perturbed dataset that is imputed by various first-principle imputation tools. The colorbar shows -log10-transformed p-values.

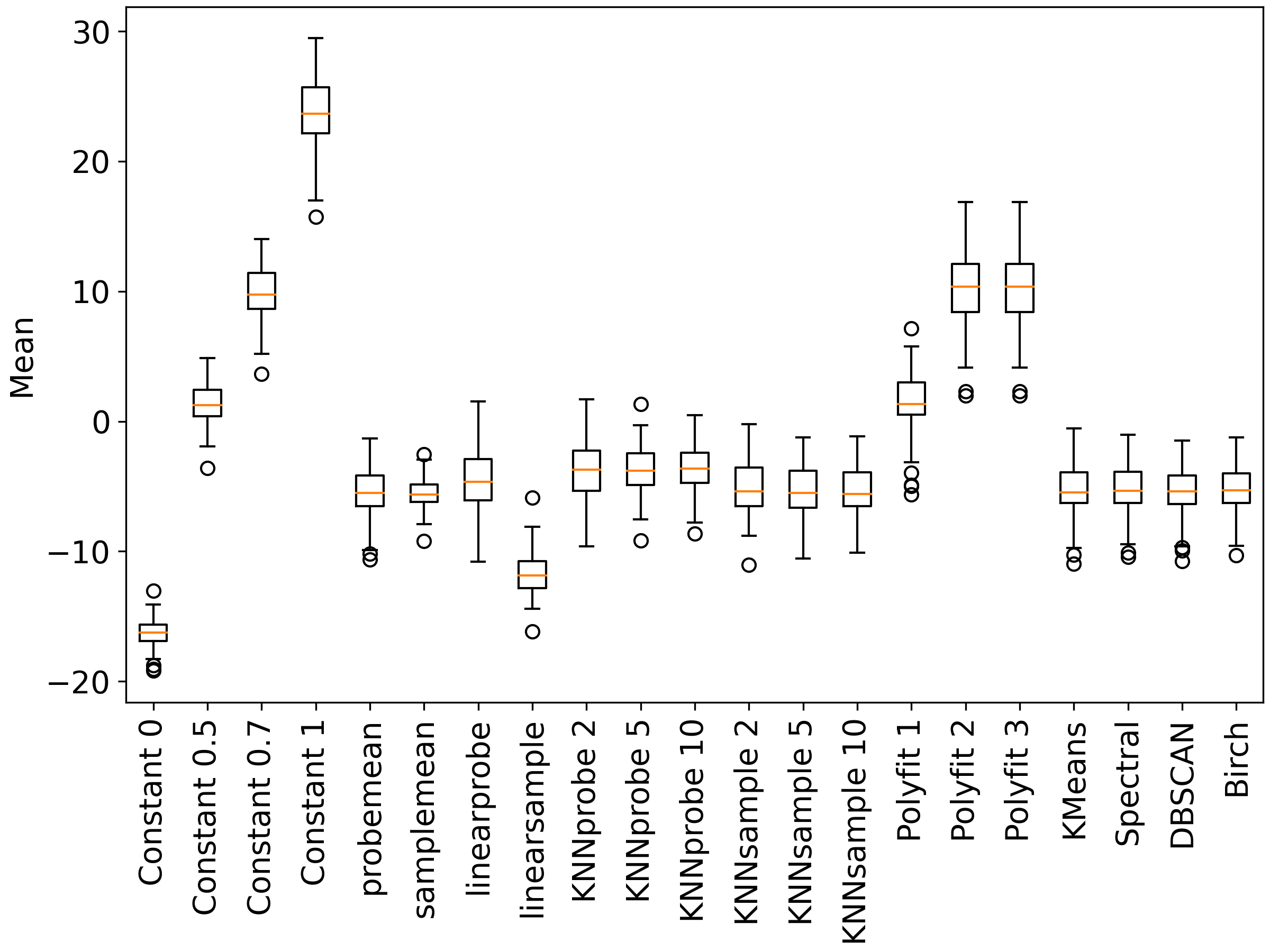

**Figure S48**. Boxplot showing distributions of mean on a heavily perturbed dataset that is imputed by various first-principle imputation tools.

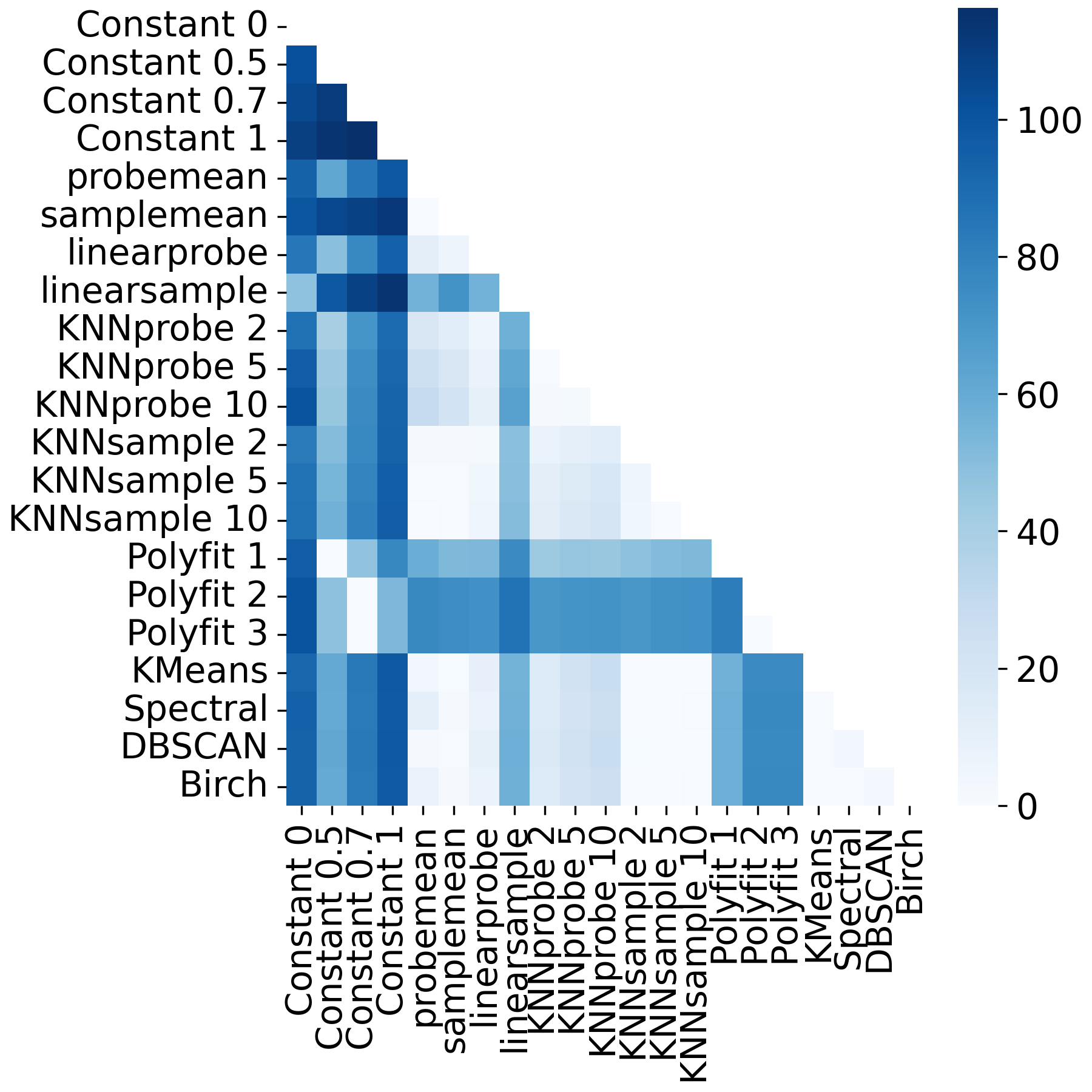

**Figure S49.** Heatmap showing pairwise t-tests of mean on a heavily perturbed dataset that is imputed by various first-principle imputation tools. The colorbar shows -log10-transformed p-values.

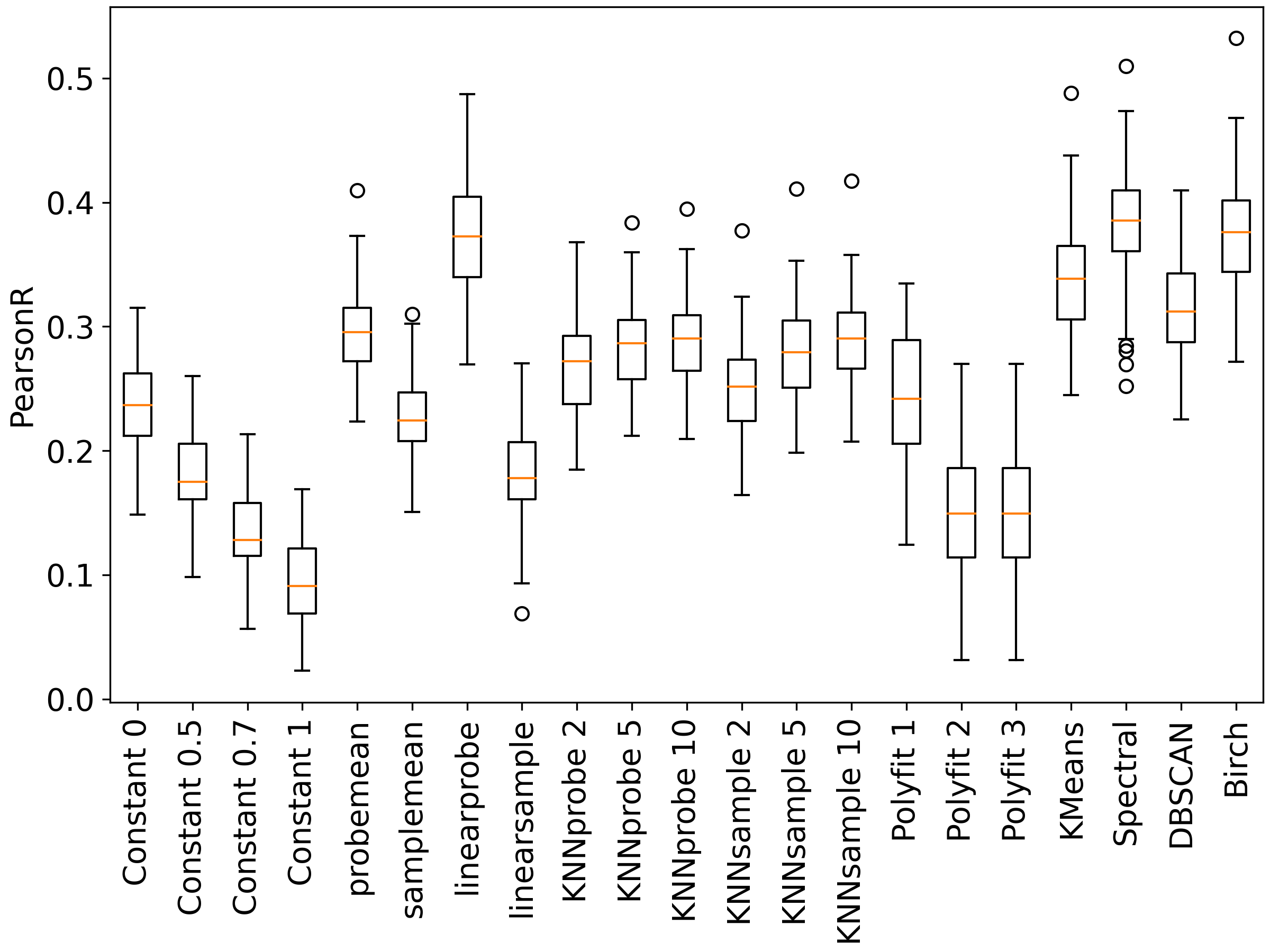

**Figure S50.** Boxplot showing distributions of Pearson correlation on a heavily perturbed dataset that is imputed by various first-principle imputation tools.

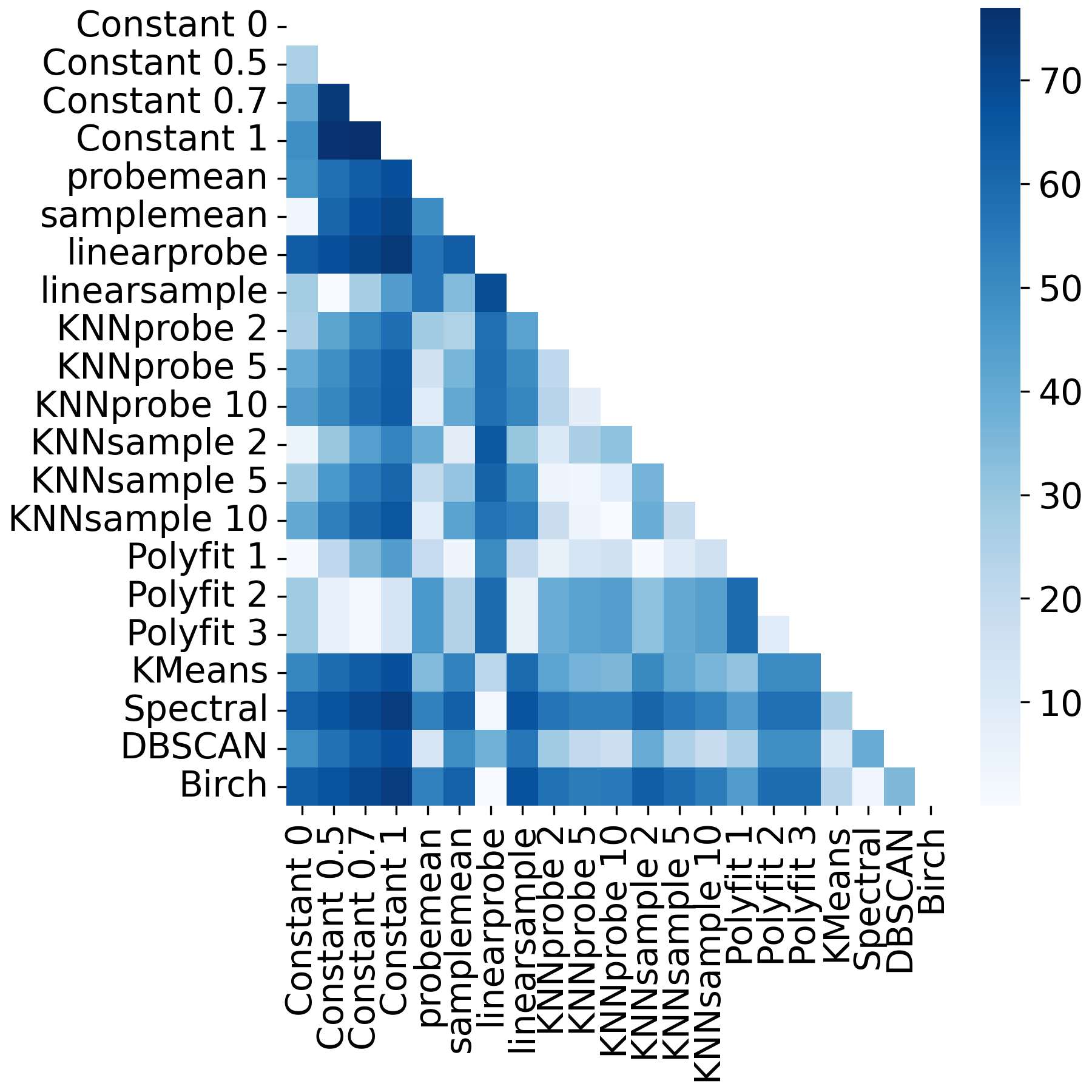

**Figure S51.** Heatmap showing pairwise t-tests of Pearson correlation on a heavily perturbed dataset that is imputed by various first-principle imputation tools. The colorbar shows -log10-transformed p-values.

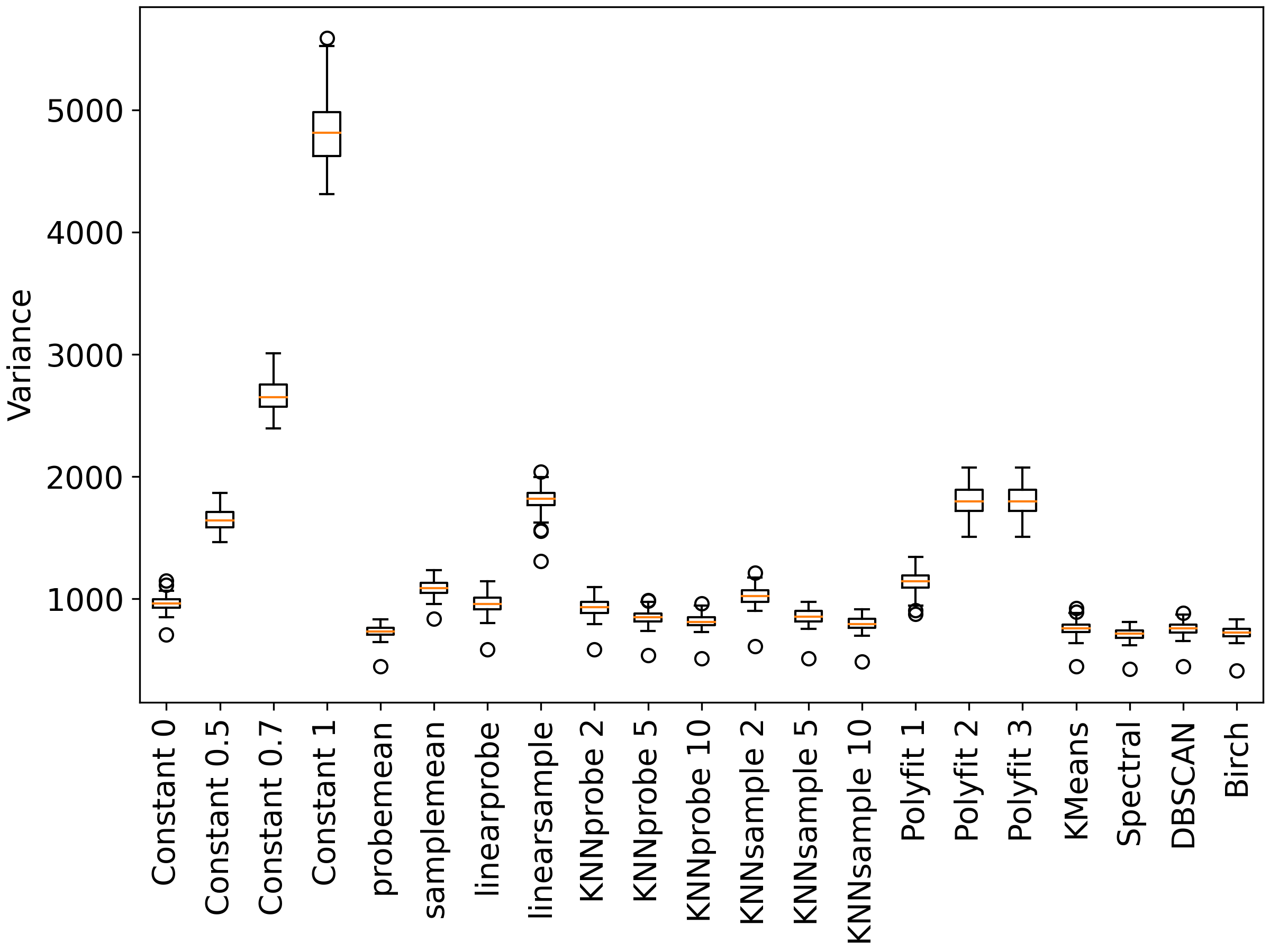
**Figure S52**. Boxplot showing distributions of variance on a heavily perturbed dataset that is imputed by various first-principle imputation tools.

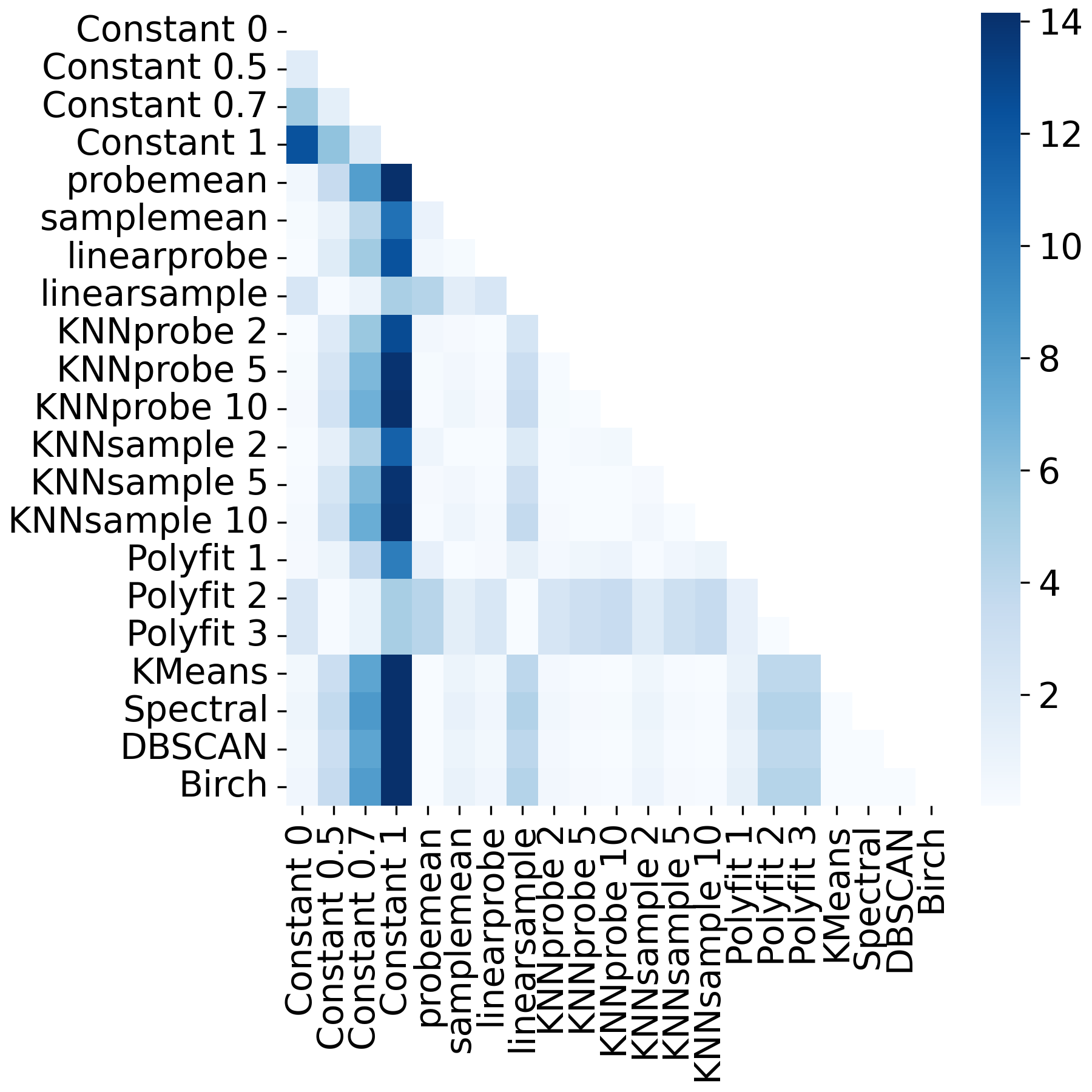

**Figure S53**. Heatmap showing pairwise F-tests of variance on a heavily perturbed dataset that is imputed by various first-principle imputation tools. The colorbar shows -log10-transformed p-values.

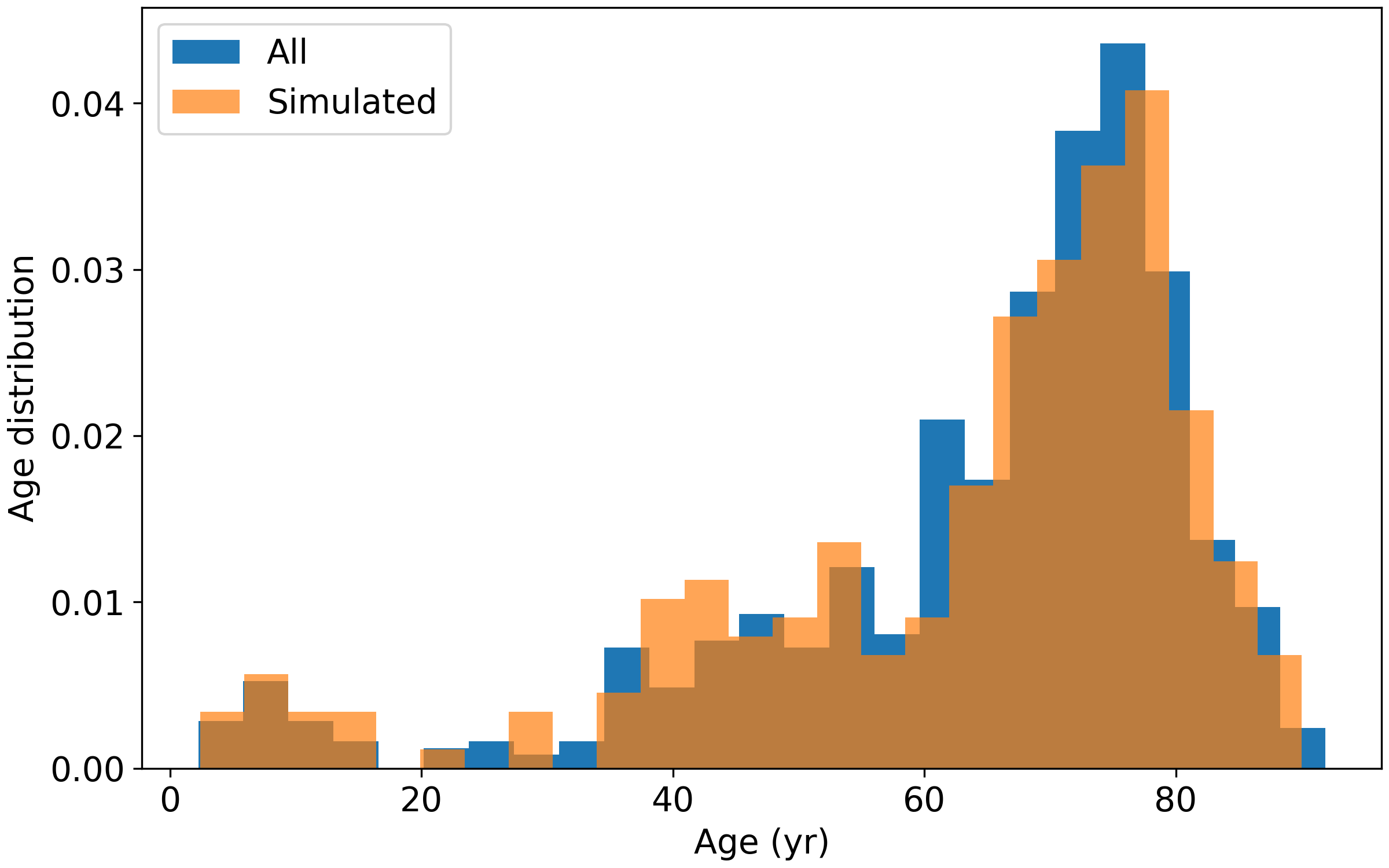

**Figure S54**. Age distributions of all samples compared to samples selected for additional imputation for testing CaMelia.
